# A transcriptional atlas of the pubertal human growth plate reveals direct stimulation of cartilage stem cells by growth hormone

**DOI:** 10.1101/2025.03.14.642964

**Authors:** Tsz Long Chu, Ostap Dregval, Farasat Zaman, Lei Li, Xin Tian, Xin Liu, Dana Trompet, Baoyi Zhou, Jussi O Heinonen, Claes Ohlsson, Lars Sävendahl, Igor Adameyko, Andrei S Chagin

**Author notes:** These authors contributed equally (Tsz Long Chu, Ostap Dregval).

## Abstract

The cartilaginous growth plate is a critical organ responsible for longitudinal bone growth. It remains open throughout life in mice but closes in humans after puberty. Growth hormone (GH) is a widely used therapy for children with growth retardation and open growth plates. However, it remains unclear whether GH directly targets human growth plates. Furthermore, while cartilage stem cells have recently been identified in mouse growth plates, their presence and GH responsiveness in human growth plates are unknown. To address these gaps, we characterized the cellular and molecular organization of early pubertal human growth plates using unique tissue samples obtained during growth-restricting surgeries. Our analysis identified two distinct populations of stem cells differing in cycling activity, molecular profiles, and regulatory factors. Quiescent stem cells were localized within a niche characterized by low Wnt and TGFβ signaling. To investigate the direct effects of GH, we developed a human growth plate explant culture system. GH directly stimulated explant growth and promoted stem cell proliferation by activating the JAK/STAT, TGFβ, and ERK pathways while inhibiting the AKT pathway. Notably, activation of the TGFβ pathway occurred in an autocrine manner. These findings provide critical new insights into human longitudinal growth and the mechanisms of GH action, with potential implications for optimizing treatments for growth disorders.

**One Sentence Summary:** This study reveals that growth hormone (GH) directly promotes proliferation within the human growth plate and activates TGFβ and ERK signaling pathways in cartilage stem cells, providing critical insights into human longitudinal growth and potential improvements in treatments for growth disorders.

## Introduction

Numerous genetic, physiological, and environmental factors influence the linear growth of children (*1*). Despite this complexity, these factors converge on a single target: the epiphyseal growth plate—a thin cartilage structure located near the ends of long bones. This organ is the primary driver of longitudinal bone growth, and its disruption, as demonstrated in genetic and surgical models in mice and humans, results in impaired bone elongation (*2-4*). Consequently, a child’s growth rate predominantly depends on the activity and functionality of the growth plate, and a wide range of growth abnormalities and disorders directly or indirectly affect this structure.

The classical view of the growth plate is that of a cartilage disc comprising chondrocytes arranged along the bone’s longitudinal axis according to their differentiation stages. At one end lies the resting zone (RZ), which houses progenitor chondrocytes. These progenitors are recruited into the proliferative zone (PZ), where they divide and assume a flattened morphology. Following several cycles of proliferation, chondrocytes exit the cell cycle and undergo hypertrophy—a process involving dramatic cell enlargement—to form the hypertrophic zone (HZ) (*5*). The progression through this differentiation trajectory is accompanied by extensive production of extracellular matrix by chondrocytes. At the end of their differentiation trajectory, hypertrophic chondrocytes die, while an extracellular matrix ultimately serves as a scaffold for new bone formation. This process, known as endochondral ossification (*6*), is responsible for the growth of long bones in all mammals (*7*).

Recent research has challenged and refined aspects of this classical model of endochondral ossification. First, many hypertrophic chondrocytes do not undergo apoptosis but instead transdifferentiate into osteoblasts and stromal cells—a process discussed extensively elsewhere (*8*). Second, stem cells were identified in the resting zone, and these cells generate all other chondrocytes in the growth plate (*9*). Their activation extends bone growth (*10*). The niche, which facilitates the maintenance and renewal of these cartilaginous stem cells, is formed by the secondary ossification center early postnatally (*11*). The niche likely provides a low-Wnt microenvironment, which is essential for the quiescence of these stem cells (*12*). However, it remains unclear whether similar cartilaginous stem cells exist in the human growth plate.

Growth hormone (GH) is the principal regulator of postnatal longitudinal growth and the primary treatment for children with growth deficiency (*13, 14*). GH responsiveness begins after infancy and persists until the end of puberty. Puberty is marked by a pronounced growth spurt driven by sex hormones in synergy with GH, after which the growth plate undergoes irreversible fusion, precluding further growth. In contrast, mice do not experience pubertal growth spurts, retain open growth plates, and continue growing into late adulthood (*4, 15, 16*).

These species differences underscore the need for human-specific studies to elucidate the clinically relevant mechanisms underlying longitudinal growth. For instance, it is not well understood why some children fail to respond to GH therapy or why responsiveness wanes over time (*17, 18*).

GH exerts its effects on the growth plate through direct and indirect mechanisms. Systemically, GH stimulates the liver to secrete insulin-like growth factor 1 (IGF1), which acts on the growth plate (*19-21*). However, liver-specific ablation of IGF1 results in only a marginal reduction in bone length (*22*). Locally, GH has been shown to directly influence the growth plate, as demonstrated in rats by the asymmetric growth of limbs following unilateral GH infusion (*23, 24*). GH promotes the proliferation of resting zone cells, including label-retaining cells within this zone (*24, 25*). These findings suggest that GH may directly target stem cells in the resting zone.

Despite these insights, critical questions remain unanswered: Does GH directly target the human growth plate? Are stem cells present in the human growth plate, and can GH specifically influence them? The molecular mechanisms of GH action within the growth plate are entirely unknown. Addressing these knowledge gaps is essential for developing more effective therapies for children with growth disorders. Given the profound differences between humans and rodents—including the absence of pubertal growth spurts and growth plate fusion in rodents—below, we explored these questions using human pubertal growth plates.

## Results

### Cellular Organization of the Pubertal Human Growth Plate

Since the human pubertal growth plate is a unique and rare tissue to obtain, we first set out to perform an extensive characterization of growth plates obtained via biopsies from pubertal children aged 12–15 (puberty stages B2–B4). These samples were collected during epiphysiodesis surgery aimed at preventing idiopathic tall stature (Fig. 1A, B), a condition more prevalent in Scandinavian countries, which have some of the tallest populations globally (*26*).

**Figure 1.**
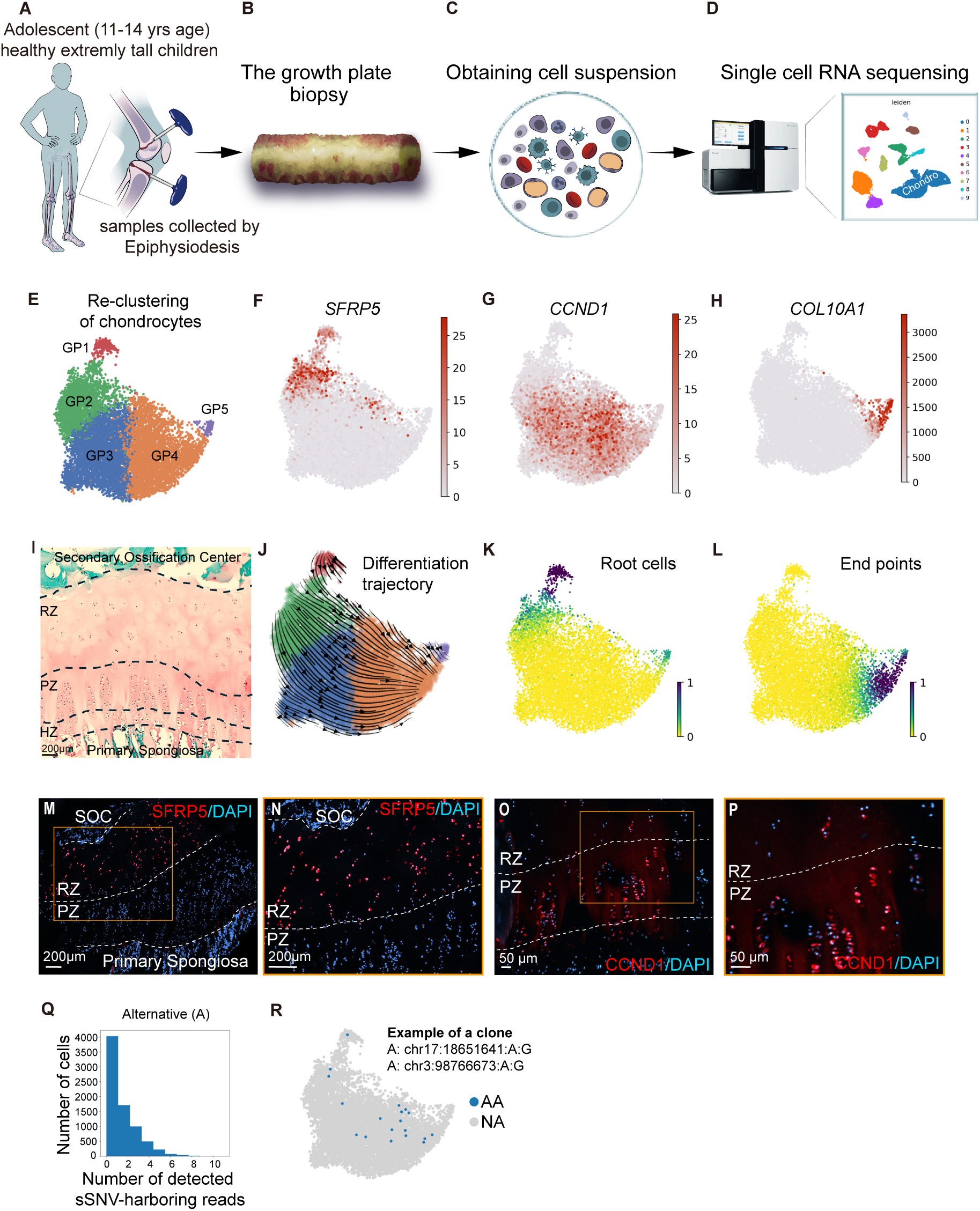
Characterization of the adolescent human growth plate at the cellular level. (A-D) Schematic representation of the experimental workflow, illustrating the collection of human growth plate biopsies during epiphysiodesis surgery (A, B), enzymatic digestion of cartilage tissue into single cells (C), and single-cell RNA sequencing (D). (B) Brightfield image showing the pubertal human growth plate biopsy, cartilage is white. (D) UMAP embedding of 24,544 cells, classified by cell type: 0 – chondrocytes, 1 – T cells, 2 – MSCs/osteoblasts, 3 – myeloid lineage, 4 – NK cells, 5 – plasma cells, 6 – naïve B cells, 7 – endothelial cells, 8 – vascular smooth muscle cells, 9 – plasmacytoid dendritic cells. (E) Re-clustering of the chondrocyte population extracted from (D) based on UMAP embedding. (F-H) Feature plots indicating expression levels of (F) *SFRP5* mRNA, (G) *CCND1* mRNA, and (H) *COL10A1* mRNA. Color intensity corresponds to raw mRNA counts per cell. (I) Representative Safranin-O/Fast Green-stained section of the human growth plate, showing key structural zones: Resting Zone (RZ), Proliferative Zone (PZ), Hypertrophic Zone (HZ), and Primary Spongiosa (PS). (J) RNA velocity analysis infers differentiation trajectories from RNA splicing dynamics, with (K) Root and (L) Endpoint cells visualized on the embedding. (M, N) Confocal images of *SFRP5* mRNA (red) and nuclei stained with DAPI (blue). (N) Magnified view of (M). SOC – Secondary Ossification Center. (O, P) Confocal images of *CCND1* protein (red) with DAPI-stained nuclei (blue). (P) Magnified view of (O). (Q) Histogram showing the number of reads harboring somatic single-nucleotide variants (sSNVs) per cell. (R) Embedding highlighting cells from one patient harboring simultaneously two sSNVs.

We dissected the cartilage from bone marrow under a stereomicroscope on ice, enzymatically digested for 30 minutes, and subjected to single-cell RNA sequencing (scRNAseq). 23,625 cells from four children passed quality control (Fig. 1C, D; Fig. S1A–C). Initial UMAP embedding revealed ten distinct cell types, including chondrocytes, endothelial cells, osteoblasts, and various hematopoietic-lineage cells (Fig. 1D; Fig. S1D).

Next, we focused on the chondrocyte cluster, comprising 9,507 cells, which showed the expression of chondrocyte-specific markers SOX9, Aggrecan, and Collagen type II (Fig. S1D–G). This cluster was computationally isolated and regressed for cell cycle and stress signatures. After optimizing the granularity of UMAP embedding by taking advantage of the literature and transcriptional differences among subclusters, we finalized the embedding with five subclusters, designated as growth plate clusters GP1–GP5 (Fig. 1E; Fig. S1H). All four samples contributed individual cell transcriptomes across these clusters; the total cell counts increased progressively from GP1 to GP5 (Fig. S1I-K).

Clusters GP1 and GP2 were enriched in the expression of progenitor cell markers SFRP5 and APOE (*27, 28*) (Fig. 1F; Fig. S1L), while Cyclin D1-positive proliferating cells were predominantly localized to GP3 (Fig. 1G). GP5 expressed the hypertrophic marker COL10A (Fig. 1H), whereas pre-hypertrophic markers IHH and MEF2C labeled GP4 and GP5 (Fig. S1M, N). The unexpectedly lower abundance of hypertrophic chondrocytes likely reflects the technical challenges of encapsulating large cells using the 10X scRNAseq platform.

Notably, the growth plate biopsies were obtained under live X-ray guidance to exclude contamination with articular cartilage. However, surgical exclusion of perichondrial cells is not entirely guaranteed. Gene expression analysis for periostin, a perichondral cell marker, confirmed the absence of these cells in our dataset (Fig. S1O).

The distribution of markers reflecting progressive development of cartilage and endochondral ossification suggests a differentiation trajectory from cluster 1 to cluster 5, consistent with the classical organization of the growth plate into the following morphological zones: (i) the resting zone, which contains stem cells, (ii) the zone of flat proliferating chondrocytes, and (iii) the zone of hypertrophic chondrocytes (*6, 28*) (Fig. 1I). However, the two clusters, expressing resting zone markers (clusters 1 and 2) came out unexpectedly.

To elucidate the hierarchical relationships between these clusters, we performed a VeloCyto analysis, a computational approach that infers differentiation trajectories based on the ratio of unspliced to mature mRNA transcripts (*29*). This analysis supported a trajectory from cluster 1 to cluster 5, identifying cluster 1 as the root cluster and cluster 5 as the terminal endpoint (Fig. 1J–L; Fig. S1 1P, R). To validate the spatial organization of these clusters, we visualized the expression of *SFRP5* and Cyclin D1 on tissue sections (Fig. 1M–P). SFRP5-positive cells clearly marked the resting zone cells, whereas Cyclin D1-positive cells were found beneath them and aligned well with the histological appearance of the human pubertal growth plate (Fig. 1I, Fig. S2A-E). This spatial arrangement corroborates the inferred differentiation trajectory observed in the dataset.

To confirm the stem cell nature of resting zone clusters, we further elucidated the genealogical, clonal connections between the clusters using the distribution of somatic single-nucleotide variants (sSNVs) in RNA reads. For this, we utilized *Monopogen*, a recently developed algorithm designed to detect and track somatic sSNVs within scRNAseq data (*30*). After filtering for potential sequencing errors, low coverage, and putative germline mutations, we identified 1,056 sSNVs. Most of these variants were present in fewer than 100 cells, although a subset exhibited higher abundance (Fig. S2F, G). As expected, the rate of sSNV detection correlated with sequencing depth (Fig. S2H). Due to the limitations of sequencing depth, each cell contained only a small number of reads at positions with either alternative (A) or reference (R) alleles (Fig. 1Q, Fig. S2I). An example of a well-covered clone highlights cells with detected reference and alternative reads for a single variant located at position chr17:18651641:A:G, where the reference allele is adenine (A) and the alternative allele is guanine (G) (Fig. S2G). This finding suggests that cells carrying this particular sSNV are genealogically related. To increase confidence in identifying clonally related cells, we further focused on cells harboring two sSNV positions that were sequenced simultaneously (Fig. 1R). Although rare, these clones were observed across all clusters, providing evidence of direct genealogical relationships between them. Notably, the frequency of sSNVs (calculated as the ratio of A to A+R) remained consistent across the growth plate (Fig. S2J), indicating no detectable accumulation of somatic SNVs within the growth plate snapshot analyzed.

The above analysis revealed two unexpected and notable features of the human pubertal growth plate: (i) the resting zone in the human growth plate is proportionally larger than anticipated based on mouse data, and (ii) it contains two transcriptionally distinct clusters of progenitor cells. These findings challenge our previous understanding derived from mouse growth plate studies and may underline known species-specific differences, such as the pubertal growth spurt and growth plate fusion observed in humans. To better understand these differences, further investigation into the quiescence, proliferative activity, and molecular properties of these progenitor cells is required and performed below.

### Molecular Organization of the Pubertal Human Growth Plate

To investigate these human-specific features of the growth plate in greater detail, we next sought to map the time course of transcriptional changes during pubertal chondrogenesis. Using DeSeq2, we identified 1,635 differentially expressed genes (DEGs) across subclusters and aligned them along the differentiation trajectory using SCFATE (*31*) (Fig. 2A; full list in Supplementary Data File 1). Known state-specific markers, such as APOE, CLU, and PTCH1 (resting zone); PTHR1 and ALPL (pre-hypertrophic zone); and COL10A1 and MMP13 (hypertrophic zone), aligned as expected along the trajectory (Fig. 2A, B). Additionally, key components of the well-known PTHrP-IHH feedback loop, which keeps the growth plate open (*6, 32*), such as PTCH1, PTH1R, and IHH (Fig. 2A), were consistently aligned, corroborating previous reports on the human pubertal growth plate (*33*).

**Figure 2.**
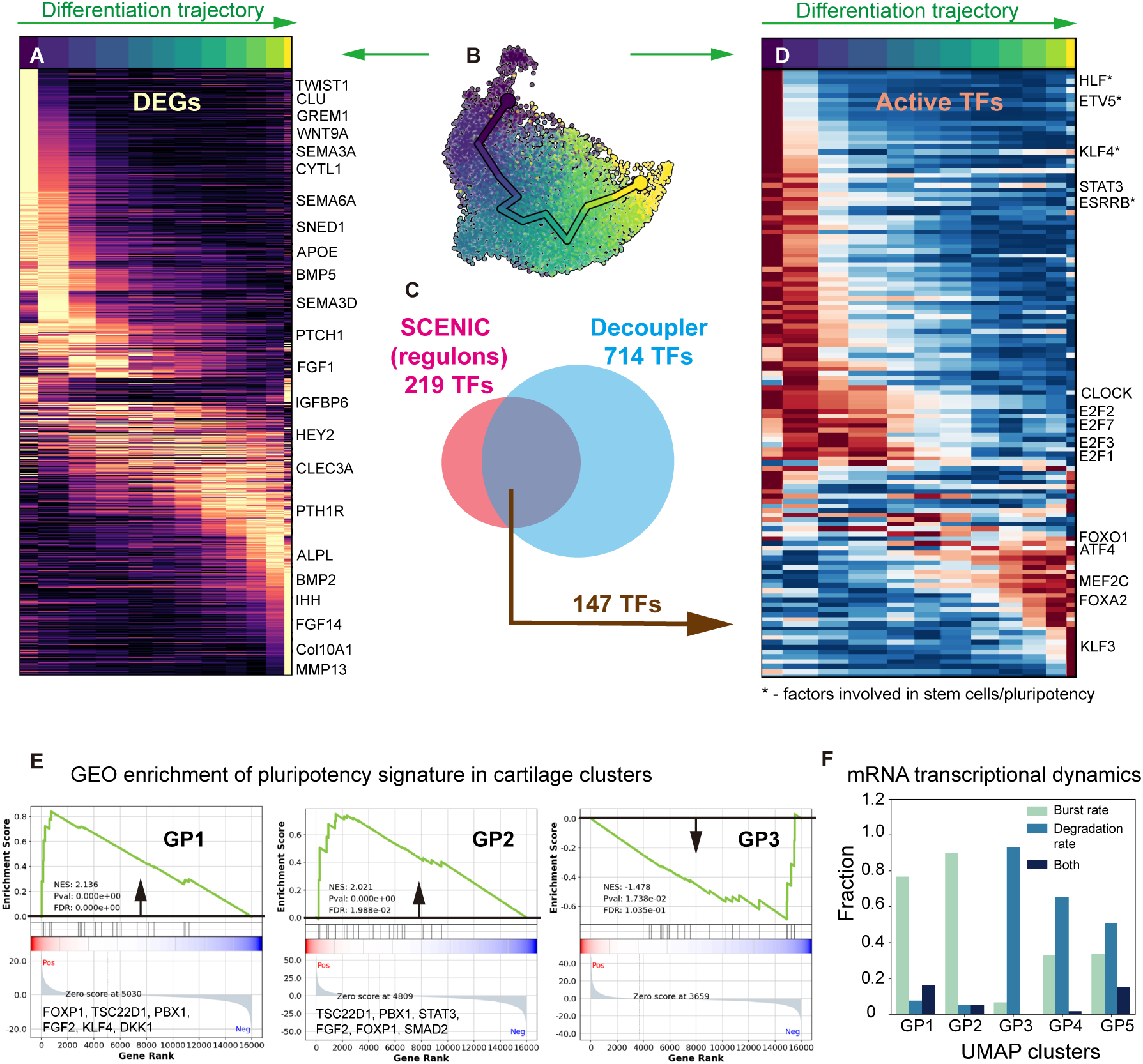
Molecular organization of the pubertal human growth plate. (A) Heatmap displaying normalized expression of genes aligned along the differentiation trajectory. Representative genes are annotated on the right, illustrating distinct patterns of gene regulation. (B) Differentiation trajectory visualized on the UMAP embedding, inferred using the SCFATE algorithm. (C) Venn diagram showing the overlap between active transcription factors (TFs) detected by Decoupler, regulons identified by SCENIC, and those identified by both algorithms. (D) Heatmap illustrating the activity of transcription factors identified by both Decoupler and SCENIC along the differentiation trajectory. Activity patterns reveal TF dynamics across differentiation stages. (E) Gene Set Enrichment Analysis (GSEA) results for the Reactome pathway “Transcriptional Regulation of Pluripotent Stem Cells,” highlighting functional enrichment in clusters GP1, GP2, but not GP3. (F) Histogram depicting relative mRNA dynamics in clusters GP1-GP5: newly synthesized mRNAs (Burst size), partially degraded mRNAs (Relative degradation), and high-turnover mRNAs (Both). The clusters GP1-GP5 correspond to those identified in Figure 1E, providing further insights into transcriptional activity and stability along the differentiation continuum. GP – Growth Plate.

The pubertal growth spurt, characterizing human but not mouse longitudinal growth, is driven by sex hormones and growth hormone (*20*), and we mapped receptors for estrogen (ESR1 (alpha), ESR2 (beta)), androgen (AR), and growth hormone (GHR). Interestingly, the expression of all these receptors was cluster-specific and predominant in chondroprogenitors, i.e., clusters GP1 and GP2 (Fig. S2K-N). We also noted that the expression of various collagens varied across zones, suggesting differential extracellular matrix composition (Fig. S2O-S). Thus, the SCFATE-arranged list of DEGs may provide a valuable tool to identify stage-specific markers for the human pubertal growth plate (Supplementary Data File 1).

To explore transcriptional regulation along the differentiation trajectory, we employed Decoupler (*34*), which predicted 713 active transcription factors (aTFs) based on expression and known target genes (Fig. 2C). SCENIC (*35*), an independent approach based on co-expression of TFs and cis-regulatory motifs (regulons), predicted 219 aTFs (with the strictest parameters, see methods), with 147 overlapping between methods (Fig. 2C). We focused subsequent analysis on these 147 TFs, given their robust identification by both approaches, and aligned them along the differentiation trajectory using SCFATE (Fig. 2D; Supplementary Data Files 2 and 3). Key TFs known to regulate chondrocyte hypertrophy in mice, such as MEF2C, ATF4, and FOXA2 (*36*), were activated at the transition to the hypertrophic stage (Fig. 2D), validating our approach. ATF4 appeared early along the differentiation as compared with MEF2C, and immunostaining showed the early expression of ATF4 compared to MEF2C (Fig. S3A-D), confirming the predicted sequence of TFs’ activation.

The transition to the proliferative stage was marked by activation of E2Fs (E2F1, E2F2, E2F3, E2F7), key cell cycle regulators (Fig. 2D). Interestingly, CLOCK, a circadian rhythm regulator, was active early in this domain (Fig. 2D). This aligns with longstanding discussions on diurnal growth in children (*37, 38*) and adds to limited direct evidence on circadian rhythm’s role in longitudinal growth (*39*).

Terminal hypertrophy was marked by aTFs such as KLF3, SRF, CEBPD, KLF12, and BACH2 (Fig. 2D; Supplementary Data Files 2 and 3), with notable enrichment of Kruppel-like factors (KLF10, 3, and 12). Among these, KLF10 is implicated in modulating chondrocyte hypertrophy in mice (*40*), while others remain unexplored.

Unexpectedly, a specific family of transcription factors, the Krüppel-like zinc finger proteins (KLFs), exhibited a distinct pattern of expression and transactivation activity along the cartilage differentiation trajectory—from early progenitors to hypertrophic chondrocytes. Notably, eight out of 17 known KLFs were identified in our cross-validated list of active transcription factors (12 when considering SCENIC analysis alone), demonstrating significant enrichment and stage-specific regulation (Fig. S3E). While some KLF transcription factors have been previously implicated in skeletal development, their roles in longitudinal bone growth remain largely unexplored (*41*). The observed stage-specificity highlights their potential pivotal roles in human bone growth, underscoring the need for further investigation.

Among those KLFs, KLF4 Yamanaka’s pluripotency factor (*42*) is selectively active in chondroprogenitors (Fig. 2D, Fig. S3E). Another well-known stemness-associated TF, ESRRB (*43*), was also active in chondroprogenitor clusters GP1 and GP2 (Fig. 2D). This prompted us to check if other TFs implicated in stem cell regulation or lineage specification are enriched in GP1 and/or GP2 clusters. It appeared that HLF and MECOM (hematopoietic stem cells), ETV5 (neuronal stem cells), POU3F1 (spermatogonia stem cells), HOXA13, FOXC1 and TBX18 (developmental lineage specification) were specifically active in either GP1, GP2 or both (Fig. 2D and Supplementary Data Files 2 and 3). This prompted us to check for pluripotency gene enrichment and, indeed, gene set enrichment analysis (GSEA) confirmed pluripotency gene enrichment in GP1 and GP2 but not in GP3 (proliferative chondrocytes) (Fig. 2E). Finally, we analyzed the splicing kinetics throughout the clusters utilizing recently developed biVI method (*44*). It revealed that cells in GP1 and GP2 clusters mostly comprise transcripts in the active synthesis, whereas the GP3 cluster mostly has transcripts, which undergo degradation (Fig. 2F). The clusters GP4 and GP5 showed relatively similar rates of newly synthesized and degrading transcripts and genes with high mRNA turnover were limited to GP1, GP2 and GP5 clusters (Fig. 2F). This observation indicates that cells in GP1 and GP2 clusters rely on different gene regulatory strategy as compared with the rest of the growth plate.

This molecular analysis offers the most comprehensive characterization of the human pubertal growth plate to date, serving as a valuable resource for diverse growth-related studies given the unique nature of the tissue analyzed. Notably, it revealed that clusters GP1 and GP2 are enriched for pluripotency-associated genes, rely on different regulatory strategies, and exhibit activated transcription factors (TFs) regulating multipotency. Although formal stemness criteria, such as self-renewal and multipotency, cannot be assessed in unmanipulated human samples, these findings—together with velocity analysis (Fig. 1J, K)—suggest that GP1 and GP2 may correspond to the human equivalents of recently identified mouse epiphyseal stem cells (*45*).

### Two Subpopulations of Chondro-Progenitors

The observed enrichment of pluripotency factors within clusters GP1 and GP2 and their different gene-regulatory strategy are highly intriguing and motivated us to investigate the differences between these clusters in greater detail. Both clusters expressed resting zone markers SFRP5 and APOE (Fig. 1F, Fig. S1L). However, PTHrP (encoded by *PTHLH*), a marker for mouse epiphyseal stem cells (*9*), was observed exclusively in GP2 but not in GP1 (Fig. 3A, B). Conversely, CYTL1, a marker for mouse skeletal stem cells derived from the growth plate region (*46, 47*), was specifically expressed in GP1 (Fig. 3C). Interestingly, among the top 25 genes most specific to mouse skeletal stem cells (root population from (*47*)), 13 were enriched in GP1 or both GP1 and GP2 (Fig. 3D). This significant overlap may highlight conserved molecular features despite differences in species, timing, and sequencing approaches.

**Figure 3.**
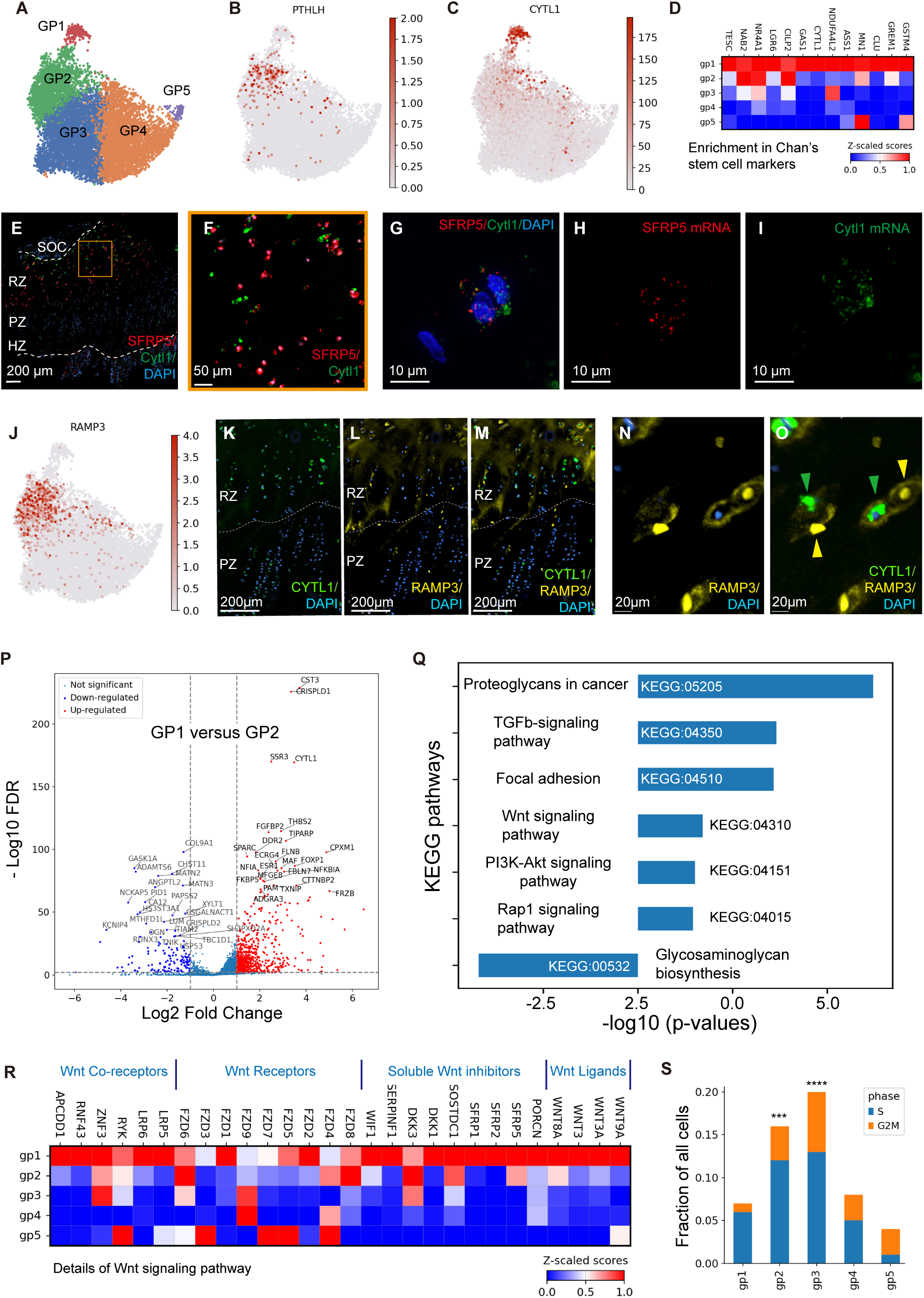
Molecular signatures of two clusters detected in the resting zone of the human growth plate. (A) UMAP clustering of human growth plate chondrocytes, reproduced from Fig. 1E for orientation. (B, C) Feature plots showing mRNA expression of (B) *PTHLH* and (C) *CYTL1*. Expression levels highlight their distinct spatial patterns in the resting zone. (D) Heatmap illustrating enrichment of genetic markers from mouse skeletal stem cells (Ambrosi TH et al., ref 45) across the clusters. (E-I) Representative confocal images displaying co-detection of *SFRP5* (red) and *CYTL1* (green) mRNAs. *RNAscope* was used for mRNA detection, with nuclei counterstained by DAPI (blue). (E) General view of staining; (F) magnified image of the resting zone; (G-I) low signal amplification and high magnification show intracellular distribution of *SFRP5* and *CYTL1* mRNA expression. (J) Feature plot showing enrichment of *RAMP3* mRNA in the GP2 cluster, emphasizing cluster-specific expression. (K-M) Confocal immunofluorescence images of *CYTL1* (green) and *RAMP3* (yellow) proteins, showing individual channels for (K) *CYTL1*, (L) *RAMP3*, and (M) merged staining. (N, O) High-magnification scans of the resting zone from (M) highlighting neighboring cells positive for either *CYTL1* or *RAMP3*. Nuclei are stained with DAPI (blue). (P) Volcano plot of differentially expressed genes (DEGs) between GP1 and GP2 clusters. The Y-axis shows -log False Discovery Rate (FDR). Genes elevated in GP1 are marked in red, and those in GP2 are marked in blue. (Q) KEGG pathway analysis of DEGs identified in (P), highlighting pathways enriched in each cluster. (R) Heatmap displaying expression of Wnt ligands and receptors associated with the KEGG Wnt pathway identified in (Q). (S) Histogram illustrating the relative proportions of cells in the S and G2M cell cycle phases across chondrocyte clusters, showing differences in proliferative activity. Data in (D) and (R) are normalized across genes to highlight cluster-specific differences, while (B), (C), and (J) depict raw mRNA counts per cell. SOC – Secondary Ossification Center; RZ – Resting Zone; PZ – Proliferative Zone; HZ – Hypertrophic Zone; PS – Primary Spongiosa. *** p < 0.005, **** p < 0.00005 by chi-squared contingency test.

Spatially, GP1 cells were located within the SFRP5-positive population in the resting zone, as verified by both immunohistochemistry and RNAscope (Fig. 3E-I). Using CYTL1 and RAMP3 as markers for GP1 and GP2, respectively (Fig. 3J), we distinguished these clusters, as detecting PTHrP in human samples proved challenging. This analysis revealed that CYTL1+ and RAMP3+ cells formed two spatially distinct populations within the resting zone, often located in close proximity (Fig. 3K- O). This spatial organization resembles dyads or chondrons, potentially derived from a single progenitor cell (Fig. S2A, B).

Differential expression analysis identified 819 and 165 significantly up- and downregulated genes, respectively, distinguishing GP1 from GP2 (Fig. 3P). Gene ontology and KEGG pathway enrichment analyses revealed glycosaminoglycan biosynthesis enriched in GP2 and six pathways enriched in GP1, including TGFβ, Wnt, and PI3K-Akt (Fig. 3Q). The TGFβ pathway, which plays a well-established role in skeletal and stem cell biology (*48*), warranted closer examination. Manual annotation separated KEGG TGFβ pathway genes into BMP-related (activating SMAD1/5/8) and TGFβ-related (activating SMAD2/3) groups (Fig. S4A, B). While TGFβ members were primarily enriched in GP1, BMP pathway genes were more prevalent in GP2 and GP5 (Fig. S4A, B). Notably, GP1 exhibited upregulation of several soluble TGFβ inhibitors, including THBS1, THBS2, THBS4, and DCN (Fig. S4A), suggesting that TGFβ signaling may be actively downregulated in this cluster. To confirm this, we re-evaluated our stringent selection of active TFs, identifying only SMAD2, with its activity lowest in GP1 and highest in GP2 (Fig. S4C-E). Furthermore, SOX4 regulon, a downstream effector of SMAD2/3, was active in the hypertrophic cluster GP5 but absent in GP1 and GP2 (Supplementary Data File 3). These results further suggest that TGFβ signaling is suppressed in GP1. Additional gene expression analysis revealed that GP1 was characterized by inhibitors such as THBS2 (7.5-fold upregulation), THBS4 (5.7-fold), and THBS1 and DCN (2.6-fold each) (Fig. S4F). Non-normalized expression data (normalized for sequencing depth but not between genes or clusters) confirmed that THBS1, THBS2, and DCN were the most highly expressed TGFβ-related genes in GP1 (Fig. S4G), underscoring active inhibition of the pathway.

The Wnt pathway (KEGG:04310), identified here, was identical to the Wnt-inhibitory environment found for the quiescent mouse epiphyseal stem cells described in prior studies (*12*). Furthermore, GAS1, a marker for mouse label-retaining epiphyseal stem cells (*12*), was specifically expressed in GP1 (Fig. 3D). Among 174 genes in the Wnt pathway, GP1 cells expressed four ligands and nine soluble inhibitors (Fig. 3R). The absence of β-catenin-related TF activity in GP1 (Supplementary Data Files 2 and 3) further suggested a Wnt- low microenvironment similar to that observed in quiescent stem cells in mice (*12*). Additionally, proliferation analysis revealed that GP1 exhibited the lowest proliferation rate compared to GP2 and GP3 (Fig. 3S), reinforcing its quiescent phenotype.

Taken together, these observations suggest that GP1 and GP2, both located within the resting zone, represent two subpopulations of epiphyseal stem cells: quiescent (GP1) and more proliferatively active (GP2). Considering their spatial arrangement, velocity analysis (Fig. 1J, K), and gradual recruitment of transcriptional machinery during the GP1-to-GP2 transition (Fig. 2D, Supplementary Data File 2), it is plausible that GP1 cells serve as a parental population for GP2 cells and may correspond to label-retaining stem cells whereas GP2 may correspond to column-generating stem cells observed in mice (*9, 11, 12*). The quiescent GP1 population resides in a Wnt- and TGFβ-low microenvironment, emphasizing its role as a reservoir of stem cell activity.

### Potential Interactions Between Different Clusters

We observed that several morphogens, including IGF1, FGFs, WNTs, BMPs, and SEMAs, are expressed in a stage-dependent manner throughout the growth plate (Fig. 2A; see also Supplementary Data File 1 for the full list of DEGs aligned along the differentiation trajectory). The long-standing hypothesis of an additional feedback loop within the growth plate (*49*), beyond the well-characterized PTHrP-IHH loop (*6*), prompted us to investigate ligand-receptor (LR) interactions between clusters in the human growth plate. While LR predictions can be inherently noisy and of limited predictive value, we conducted this analysis given the unique nature of the tissue under study.

For this analysis, we utilized the Rank Aggregate method from the LIANA package, which integrates and cross-validates predictions from multiple tools, including CellChat, CellPhoneDB, CellSignalIR, Connectome, and NATMI (*50*). From the growth plate dataset, 1,000 potential LR pairs were identified. These pairs were filtered to exclude collagen- dependent interactions and scored for specificity and magnitude. The top 60 highest-ranked LR pairs for each score were visualized (Fig. S5A, B), and the top 160 pairs for both specificity and magnitude are listed in Supplementary Data File 4. The classical PTHrP-Ihh feedback loop was among those identified in specificity-scored but not magnitude-scored interactions (Fig. S5A, B), supporting the validity of the specificity-based predictions.

Hypertrophic cells (GP5 cluster) exhibited the highest secretory activity, expressing a wide range of morphogens. GP1 cells showed the second-highest activity, while GP3 and GP4 were the least active and displayed no potential interactions when scored by specificity (Fig. S5A). Among the predicted interactions, Ihh from the hypertrophic zone targeted both the GP1 and GP2 clusters (Fig. S5A). Additionally, BMP8A and BMP8B were predicted to target predominantly GP2 and, to a lesser extent, GP1 clusters (Fig. S5A). Expression data showed that on top of BMP8A and B, hypertrophic cells expressed BMP2, BMP4, BMP6, and BMP7 (Fig. S4B, S5A, B). BMP4 was predicted to target GP2 but appeared among magnitude-scored rather than specificity-scored pairs (Fig. S5B). This aligns with earlier observations of BMP receptor enrichment in the GP2 cluster (Fig. S4B). However, additional BMPs were not captured by the LIANA algorithm, emphasizing the tentative nature of these predictions.

Beyond BMP signaling, we identified additional potential interactions involving FGFs, WNTs, Ephrins, PDGF, and Notch signaling pathways (Fig. S5A, B). Expression values for selected ligands and receptors associated with these pathways were also presented (Fig. S5C). While these predictions remain speculative, they provide valuable insights into the potential paracrine interactions shaping growth plate organization and function.

Although the predictions of individual cell-cell interactions remain speculative, the grouping of these interactions into the G5 hypertrophic cluster and G1/G2 stem cell populations likely reflects active and complex feedback signaling mechanisms between differentiated progeny and their parental stem cells.

### Growth Hormone (GH) directly stimulates the Growth of Human Pubertal Growth Plate

The above characterization provided a solid foundation for functional experiments. As highlighted in the introduction, while GH is the most commonly used treatment for children with growth deficiency, its direct effects on the human growth plate and resident stem cells, as well as the underlying mechanisms, remain largely unknown.

To address this, we employed our previously established model of the organ culture of the human growth plate (*51, 52*). In this model, growth plate biopsies from patients are sliced into 1 mm-thick sections, which can be cultured and treated (Fig. 4A). We set to treat the slices with vehicle and GH for two months to assess the direct growth-promoting effect of GH, and for 24 hours, to explore the underlying mechanisms (Fig. 4A).

**Figure 4.**
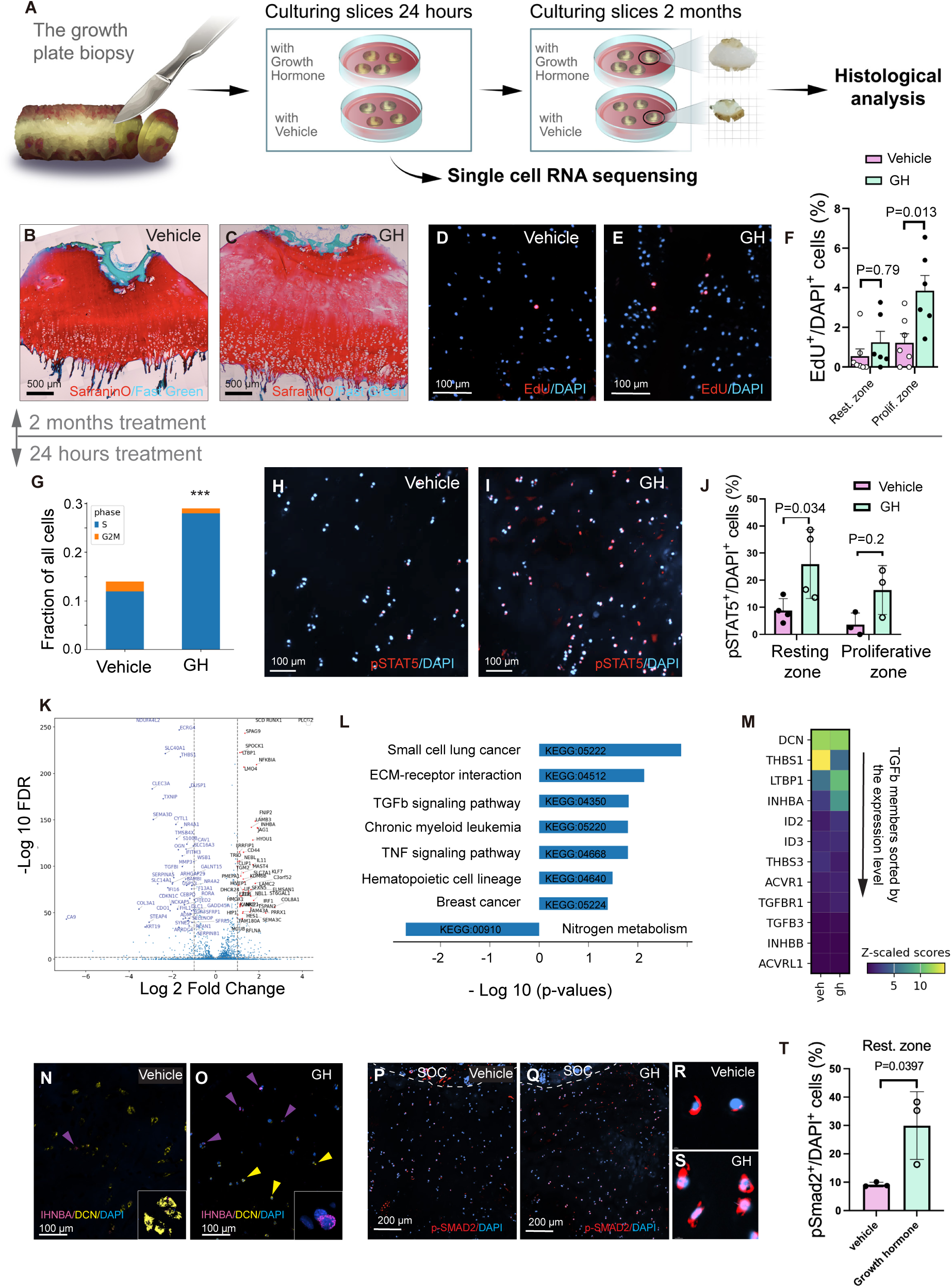
Growth Hormone (GH) directly targets stem cells and activates TGFβ signaling. (A) Schematic diagram outlining the workflow for short-term (24 hours) and long-term (2 months) organ culture of human growth plate explants treated with vehicle or growth hormone (GH). Analyses in (B-F) correspond to 2-month cultures, while (G-T) focus on 24-hour cultures. (B, C) Representative histological images of Safranin-O/Fast Green staining after 2 months of culture with (B) vehicle or (C) GH treatment. (D, E) Representative images of EdU labeling of proliferating cells (red) in (D) vehicle- and (E) GH-treated growth plate explants. Nuclei were counterstained with DAPI (blue). (F) Quantification of EdU-labeled cells in the resting and proliferative zones of vehicle (pink) and GH-treated (green) samples. Data are based on explants from 7 vehicle-treated and 6 GH-treated patients. (G) Bar graph comparing the ratio of cells in the S and G2M phases in cultured explants after short-term vehicle and GH treatment, highlighting changes in cell cycle dynamics. (H, I) Representative images of phospho-STAT5 (red) immunodetection in explants treated with (H) vehicle or (I) GH. Nuclei counterstained with DAPI (blue). (J) Quantification of phospho-STAT5-positive cells in vehicle- and GH-treated explants. Data include 4 patients per group for the resting zone and 3 for the proliferative zone. One patient’s sample was eventually lost and not used in other analyses. (K) Volcano plot of differentially expressed genes (DEGs) in the resting zone cluster between vehicle- and GH-treated samples. (L) KEGG pathway enrichment analysis of DEGs identified in (K), highlighting pathways altered by GH treatment. (M) Heatmap displaying expression of TGFβ family genes in GP2 clusters after 24-hour vehicle or GH treatment. (N, O) Representative RNAscope images showing *IHNBA* (purple) and *DCN* (yellow) mRNA expression in the resting zone of (N) vehicle- and (O) GH-treated samples. (P, Q) Immunofluorescent staining of phospho-SMAD2 (p-SMAD2, red) in (P) vehicle- and (Q) GH-treated samples. (R, S) Magnified images of p-SMAD2 staining from (P) and (Q), highlighting nuclear localization. (T) Quantification of p-SMAD2+ nuclei in the resting zone of vehicle (pink) and GH-treated (green) samples. Data are from 3 patients per group. Statistical analyses were performed using ONE-WAY-ANOVA with Turkey’s correction for multiple comparisons for (F, J), chi-squared contingency test for (G), and unpaired Student’s t-test for (T). ***p < 0.0005. SOC – Secondary Ossification Center; Rest. Zone – Resting Zone.

The long-term organ cultures revealed that human growth plate cartilage retains its integrity and activity even two months *ex vivo*, as evidenced by histology, proteoglycan abundance (SafraninO staining), proliferation, and Sox9 expression (Fig. 4B, C, Fig. S6A-O). This is not unexpected considering the avascular nature of cartilage, which is fed via diffusion *in vivo*. Bone was not expanding *ex vivo*, in contrast to cartilage, which is in line with our previous observations on long-term rodent metatarsal cultures (*53*), suggesting that the vascular supply is not required for chondrogenesis but for bone formation.

GH caused the expansion of the growth plate cartilage (Fig. 4B, C, Fig. S6A-O), albeit the effect was observed not in all patients (Fig. S6C). Whether this reflects the known variability in responsiveness to GH among children (*17*) or culture conditions remained to be elucidated but technically challenging due to the rarity of the patients.

The growth plates cultured in the presence of GH showed integral histology without vivid structural or morphological changes as compared to the control (Fig. 4B, C, Fig. S6D-O). Proliferation, while highly variable after 2 months in culture, was significantly increased by GH within the proliferative zone (Fig. 4F). The level of SOX9, a master transcription factor for chondrocytes, retained at slightly and not-significantly lower level at presence of GH (Fig. S6P, Q).

GH is a well-known anabolic hormone promoting mitochondrial biogenesis (*54*). We employed LARS2, encoding mitochondrial leucyl-tRNA synthetase essential for mitochondrial activity, as a marker for mitochondrial biogenesis. GH remarkably increased the level of LARS2 in these long-term cultures of human pubertal growth plates (Fig. S6R, S), providing a positive control of GH activity and demonstrating that promoted mitochondrial biogenesis can be among the potential mechanisms of direct GH action on the human growth plate.

We concluded that the observed responses provide a proof-of-principle that GH directly targets the human pubertal growth plate. While the mechanism may include mitochondrial biogenesis, the long-term cultures may cause effects that do not represent physiological mechanisms. To overcome these limitations, we utilized short-term cultures for elucidating the underlying mechanism(s).

### The Mechanism(s) of Direct Action of GH on Human Cartilaginous Stem Cells

We reasoned that a 24-hour period of organ culture would be sufficient to capture the initial transcriptional responses to GH treatment while minimizing transcriptional artifacts known to be associated with prolonged *ex vivo* conditions (*55*).

To validate the model on the transcription level, we first compared single-cell RNA sequencing (scRNAseq) profiles of cultured growth plate slices with those of uncultured samples. Cultured slices retained the majority of cell clusters observed in native growth plates, with the notable exception of cluster GP1 (Fig. S7A-H). Markers associated with GP1, including CYTL1 and IGF1, were either significantly reduced or absent in cultured samples (Fig. S7I, J). Despite this, the PTHrP-Ihh feedback loop—a critical regulatory mechanism— remained intact (Fig. S7G, H). Rates of proliferation were reduced in cultured samples compared to uncultured ones, while morphology was perfectly normal (Fig. S7K, L).

To address the difficulty of detecting GP1 cells in cultured samples, we employed a semi-supervised neuronal network-based model, scANVI (*56*). This method, applied to the combined vehicle- and GH-treated samples, predicted the presence of GP1 cells in cultured samples based on their uncultured counterparts (Fig. S7L-P). Although GP1 cells were detected, they were extremely scarce (Fig. S7M). While the underlying reasons for the scarcity of GP1 cells are unclear, the reduced proliferation rate in cultured conditions (Fig. S7K) may potentially hinder the distinction between quiescent and active stem cells. Consequently, the GP1 cells were not further evaluated.

Within the cultured samples, cluster c-GP2—positive for SFRP5 and PTHrP—was identified as representing cycling stem cells corresponding to the uncultured GP2 cluster (Fig. S7Q, R) Meanwhile, clusters c-GP4 and c-GP5, marked by IHH, corresponded to pre-hypertrophic and hypertrophic cells, respectively (Fig. S7S). Treatment with GH significantly increased the proportion of cells in the S-phase of the cell cycle (Fig. 4G) and elevated nuclear phospho- STAT5 levels, a direct indicator of GH signaling (Fig. 4H-J). The highest levels of phospho- STAT5 were observed in the resting zone (Fig. 4J), leading us to focus further analysis on the c-GP2 cluster.

Using Decoupler and SCENIC frameworks, as depicted in Fig. 2C, we identified 113 differentially active transcription factors (regulons) in the c-GP2 cluster upon 24h GH exposure (Fig. S8A-C). To infer potential signaling pathways downstream of the GH receptor, we utilized the CORNETO framework, which predicts intracellular signaling between inputed cell surface receptors and nuclear transcription factors (*57*). With a single known input (GH receptor) and the top 20 upregulated and downregulated regulons (Fig. S8B, C) as known outputs, this analysis predicted the activation of pathways beyond canonical JAK/STAT signaling, including MAPK1 (ERK) and GSK3B, as well as the inhibition of MAPK14 (p38α) and AKT1 (Fig. S8E). Immunohistochemical staining for phospho-ERK and phospho-AKT confirmed robust ERK activation and AKT inhibition (Fig. S8E-H), indicating that GH activates both canonical and non-canonical pathways in the human pubertal growth plate.

GH exposure for 24 hours resulted in 144 significantly up- and 120 downregulated genes (DEGs) in the c-GP2 cluster (Fig. 4K). KEGG pathway analysis of DEGs further highlighted the TGFβ signaling pathway as one of the top three most significantly altered pathways (Fig. 4L). This was particularly notable given the activity of the TGFβ/BMP pathway in native stem cells (Fig. S4). GH treatment upregulated TGFβ2, TGFβ3, INHBA (Activin A), and INHBB (Activin B), while downregulating inhibitors of the pathway, such as DCN, THBS1, and THBS4 (Fig. S9A). The most significant changes included the upregulation of TGFβ3 and Activin A, and the downregulation of THBS1 (Fig. S9B). The largest changes in the expression levels were observed for the upregulation of INHBA and LTBP1 and the downregulation of DCN and THBS1 (Fig. 4M). RNAscope analysis of tissue sections corroborated these results, showing increased expression of Activin A and decreased expression of DCN upon 24 hours of GH treatment (Fig. 4N, O). These findings suggest activation of the SMAD2/3 signaling pathway.

To confirm this, we examined the nuclear localization of SMAD proteins, observing increased levels of SMAD2/3 (indicative activation of TGFβ signaling) and decreased levels of SMAD1/5 (indicative inhibition of BMP signaling) in the resting zone upon GH treatment (Fig. 4P-T, Fig. S9C-G).

Collectively, these findings suggest that GH directly promotes the proliferation in the human pubertal growth plate and regulates both canonical and non-canonical signaling pathways in the epiphyseal stem cells. Non-canonical pathways include activation of ERK1/2 and TGFβ and inhibition of AKT. Activation of the TGFβ pathway is driven by autocrine secretion of Activin A, accompanied by the downregulation of soluble TGFβ inhibitors upon GH exposure. These results provide novel insights into the mechanisms of GH action.

## Discussion

Here, we characterized the cellular and molecular organization of the human pubertal growth plate at an unprecedented level, identified two populations of stem cells therein, demonstrated that growth hormone (GH) can promote human growth directly, and showed that GH triggers non-canonical signaling pathways and activates the local TGFβ signaling pathway in an autocrine manner. Our initial goal was to test if GH directly targets the human growth plate and, if so, to evaluate the underlying mechanisms. Early in the study, we realized that existing knowledge about the cellular and molecular organization of human growth plates was insufficient to achieve this objective, prompting an in-depth analysis of the intact growth plate. We discuss our findings in two parts: the general characterization of the human growth plate and the effect of GH on this structure.

### Cellular and Molecular Characterization of the Human Pubertal Growth Plate

Exploring the cellular and molecular machinery driving children’s growth is crucial due to species differences between mice and humans, especially for addressing growth disorders. While embryonic human limb development has been thoroughly characterized recently (*58, 59*), data from mice suggest substantial differences in mechanisms of longitudinal growth between neonatal and postnatal periods (*11*). Furthermore, responsiveness to GH is acquired postnatally (*14*). This provides a rationale for in-depth characterizing the pubertal human growth plate.

Our analysis identified numerous genes and transcription factors marking different stages of chondrocyte progression, revealing novel expression patterns. While some genes are known to be specific to various growth plate zones, many of these patterns were previously unexplored. Furthermore, active transcription factors identified by both SCENIC (*35*) and DECOUPLER (*34*) analyses bring our understanding of molecular machinery to a novel level. These complementary methods are based on different biological and computational approaches. By integrating these methods, our dataset provides a robust and comprehensive list of transcription factors active within the human pubertal growth plate. Aligning these transcription factors along the differentiation trajectory provides valuable insights into the regulatory mechanisms driving transitions between chondrogenic stages. The mechanisms underlying transitions between the stages remain poorly understood in both mice and humans. In mice, the transition from the proliferative to the hypertrophic stage is the most extensively studied and involves transcription factors such as ATF4 and MEF2C (*36, 60*). Similarly, we identified the activation of these factors during the proliferative-hypertrophic transition in the human pubertal growth plate, mirroring the mouse process. Additionally, our analysis revealed several novel transcription factors active during this transition. Additional transcription factors involved in further differentiation highlight the complexity of this process. The transcription factors regulating the transition from the resting to the proliferative stage remain poorly characterized, even in mice. Our findings provide valuable insights into this critical step of human postnatal chondrogenesis and provide a robust resource for tissue engineering and genetic studies, e.g., chondrodysplasia research.

A striking feature of human growth plates is the abundance of resting zone cells, which comprise nearly half of the structure in pubertal children. This challenges the long-standing hypothesis that growth ceases due to the exhaustion of chondro-progenitors (*61*). The presence of stem cells, as shown in mice (*9, 62, 63*), and proposed here for humans, suggests that growth cessation might involve active change of the stem cell niche, potentially driven by elevated sex hormones or GH during puberty. For example, we observed that stem cells require a TGFβ-low microenvironment, while GH activates local TGFβ signaling, offering a plausible mechanism. Unfortunately, technical limitations precluded us from formally testing this hypothesis, as the most quiescent root population was not detected under culture conditions. The limited availability of patient material also precluded the isolation and formal testing of these cells for stemness, such as self-renewal and cartilage formation under the kidney capsule in mice. Nonetheless, numerous indirect observations strongly support the stemness of resting zone cells. These include differentiation trajectory analysis, genetic links to other cell populations, the presence of established and novel stem cell markers, slow cycling activity, transcriptional similarity to skeletal stem cells reported in mice, enrichment of pluripotency-associated genes, and active stem cell transcription factors. Collectively, these findings support the hypothesis that human growth is driven by stem cells, consistent with emerging evidence that stem cells play a central role in mouse longitudinal growth (*10, 45*).

Interestingly, we identified two transcriptionally distinct populations within the resting zone. This phenomenon, unreported in mice, may reflect species differences or the unsorted nature of our dataset. Mouse scRNAseq data often involve sorting strategies, which, while reducing heterogeneity, introduce bias and require extended preparation times. Notably, 24 hours in organ culture erases these transcriptional differences while preserving morphology. Spatial analysis revealed that these populations coexist in close proximity, suggesting a genealogical relationship. Velocity and Monocle analyses support this relationship. Velocity and gradual recruitment of additional transcription machinery suggest cluster GP1 as a parental cluster for GP2. GP1 cells are quiescent and reside in a WNT-low environment, resembling label-retaining quiescent stem cells in mouse growth plates (*12*). Thus, GP1 and GP2 likely represent two transcriptional stages of stem cells: one quiescent and the other more active. Such dual stem cell populations—quiescent and clone-making—are uncommon and were reported before for tissues like the intestine and hair follicles (*64*). However, these two populations may also perform complementary functions, as shown in skull sutures, where one population facilitates fusion, whereas another keeps the suture generating osteoblasts (*65*). Similarly, human growth plates, which fuse unlike those in mice, might rely on a specialized cellular population for this process.

### The Growth Hormone’s Direct Action and Underlying Mechanisms

GH is the primary treatment for children with short stature, though its efficacy varies among individuals and diminishes over time (*14, 17*). This variability underscores the need to explore the mechanisms underlying GH action. While the systemic effects of GH via liver-produced IGF1 are well established, its direct impact on the growth plate has been less studied. Evidence for a direct effect is based on studies where GH was locally infused into one leg of hypophysectomized rats, demonstrating localized growth stimulation (*23, 24*), though these experiments do not rule out the possibility that GH’s direct effects require the presence of other factors, such as IGF1. Moreover, species differences raised questions about whether human growth plates respond directly to GH.

Ethical constraints prevent direct experiments involving local GH infusion in children. To address these questions, we employed organ cultures of human pubertal growth plates. These experiments demonstrated that GH can directly stimulate the growth of the human pubertal growth plate, which can be seen after two months of treatment. Notably, in a typical clinical setting, children are treated with GH for several years. It has to be emphasized that responsiveness to GH is acquired postnatally—at 2-3 years of age in children and 2-3 weeks in rodents (*66*). Neonatal rat metatarsal bones do not respond to GH *ex vivo* (unpublished observation), highlighting the importance of using pubertal growth plates for testing direct GH action. Although this organ culture model has been validated in previous studies (*51, 52*), it has limitations. Cartilage, as an avascular structure, typically performs well in vitro, as shown in five-month-long cultures of rat metatarsal bones (*53*). However, the subchondral bone does not form *ex vivo*, as seen in metatarsal cultures (*53*). Furthermore, serum-free conditions may lack the nutrients required for sustained cartilage growth. Perhaps the most significant limitation is the inability to replicate the pulsatile secretion of GH, which may lead to receptor downregulation and reduced responsiveness with time. Indeed, while 3 out of 5 patients showed a growth-promoting response, the absence of a response in 2 patients may reflect either intrinsic patient variability or the limitations of the culture conditions. Despite these challenges, our findings provide proof-of-principle that GH can directly stimulate human growth plate cartilage.

To explore the mechanisms of GH action, we mimicked a single pulse of GH by overnight culture of growth plates. This approach revealed increased cell proliferation following GH treatment. GH receptors were detected in both stem cells and proliferative cells, and activation of the JAK/STAT pathway—a canonical GH receptor signaling pathway (*14, 67*)—was predominantly observed in stem cells, suggesting these cells are primary targets of GH. This finding aligns with studies in rats, where GH’s effects are localized to resting zone cells (*24*). Further molecular analyses of stem cells revealed two key observations. First, GH signaling in stem cells involves non-canonical pathways, including activation of MAPK ERK1/2 kinases and inactivation of AKT kinase. While these pathways have been observed in other systems (*67*), their role in GH-mediated longitudinal growth is novel. These findings are particularly relevant in the context of therapies for achondroplasia, such as CNP analogs, which inhibit ERK1/2 (*68*). Monitoring potential interactions between such therapies and GH action will be critical.

A particularly intriguing observation was the stimulation of the TGFβ signaling pathway in human cartilaginous stem cells by growth hormone (GH). The TGFβ pathway is well-known for its role in maintaining stem cell quiescence across various organs (*69*). In this study, we observed that quiescent cartilaginous stem cells reside in a microenvironment characterized by low TGFβ signaling. The GH-mediated elevation of TGFβ signaling may disrupt this specialized microenvironment, potentially inducing stem cell activation. Consistent with this, we observed increased proliferation of stem cells following GH treatment, though it remains unclear whether this response originated from quiescent or already activated stem cells. Although direct regulation of the TGFβ pathway by GH is not extensively documented, one recent study demonstrated that GH stimulates TGFβ signaling in mouse podocytes by upregulating TGFβ1 expression (*70*), supporting the possibility of a similar interplay. Here, the GH-dependent downregulation of local TGFβ inhibitors further suggests that this interaction may occur through autocrine regulation of TGFβ modulators. Interestingly, neutralizing antibodies targeting the TGFβ pathway are currently under clinical investigation for treating fibrodysplasia ossificans progressiva (*71*). Monitoring GH levels and growth rates in these patients could provide valuable insights into the interaction between GH and TGFβ pathway.

### Limitations of the Study

The primary limitation of this study is the rarity of children undergoing epiphyseodesis surgery. This procedure is performed predominantly in Sweden and the Netherlands, nations among ten tallest globally (*26*), where surgical termination of normal growth is sometimes considered. Our clinical center serves all of Scandinavia, yet we can obtain only 3–5 samples annually. This scarcity highlights the uniqueness of the tissue studied but imposes certain constraints. For instance, we combined data from both boys and girls and were unable to categorize patients as GH-responsive or non-responsive, limiting our ability to address this critical distinction in detail. Although the patients in our cohort were generally normal, characterized by idiopathic tall stature and tall parents, the possibility of an underlying genetic cause cannot be excluded entirely despite all the patients being examined by the experienced pediatric endocrinologist. Thus, extrapolation of our findings to the general population should be approached with caution.

Functionally, our study was confined to *ex vivo* experiments. We minimized potential limitations by employing organ cultures and reducing culture time to the practical minimum, but results should still be interpreted carefully. One notable drawback of these cultures is the inability to transcriptionally identify the cluster of quiescent stem cells. Consequently, we could not explore their regulation by GH. Based on morphological analysis, we hypothesize that these cells remain present but lose their quiescence during overnight culture. Alternatively, the GP2 cluster of active stem cells may lose the appropriate microenvironment, i.e., the niche, which can lead to the death of these cells or loss of their unique transcriptional signature. Both options would preclude identifying this cluster transcriptionally. Despite these limitations, our findings offer valuable insights into human growth plate biology.

## Conclusion

Our study provides a comprehensive characterization of the cellular and molecular organization of the human pubertal growth plate and uncovers novel mechanisms of GH action. By identifying stem cells as primary targets and uncovering novel signaling mechanisms, including non-canonical pathways and TGFβ pathway activation, this study not only enhances our understanding of growth regulation but also has implications for optimizing GH therapy and exploring its interactions with emerging treatments. Altogether, the findings offer a solid foundation for improving therapies for growth retardation and understanding congenital skeletal abnormalities. Furthermore, the insights gained here hold promise for advancing tissue engineering and exploring the broader applications of GH as an anabolic agent.

## Data availability statement

All data and code are available via the link https://www.ncbi.nlm.nih.gov/geo/query/acc.cgi?acc=GSE288028 Original images and tissue sections can be obtained from the corresponding author upon reasonable request.

## Acknowledgments

We express our gratitude to Olga Kharchenko for her professional illustrations. We also acknowledge SciLifeLab for facilitating the single-cell RNA sequencing and providing access to the associated computational resource, UPPMAX. Additionally, we thank the University of Gothenburg, Karolinska Institutet, and the Medical University of Vienna for their premises and infrastructure that supported this work.

This study was financially supported by: the Swedish Research Council (grant 2020-02298 to ASC; grant 2020-02025 to LS), the Novo Nordisk Foundation (grant NNF21OC0070314 to ASC.), the Swedish state under the ALF agreement between the Swedish government and county councils (ALFGBG-966178 to ASC; ALF-Stockholm RS2021-0855 to LS)

We are sincerely grateful for this generous support, without which this research would not have been possible.

## Author contributions

Conceptualization: ASC

Methodology: NTLC, OD, FZ, IA, LL, XT, XL, DT, BZ, JOH

Investigation: ASC, NTLC, OD

Visualization: ASC, NTLC, OD, DT

Funding acquisition: ASC, LS

Project administration: ASC

Supervision: ASC, LS

Writing – original draft: ASC

Writing – review & editing: ASC, IA, NTLC, CO

## Supplementary materials

Supplementary Material File 1: Figures S1-S9, Methods, Tables S1, S2

Supplementary Data file 1 (all DEGs aligned along the differentiation trajectory)

Supplementary Data file 2 (all active TFs aligned along the differentiation trajectory)

Supplementary Data file 3 (the same active TFs with their activity shown on UMAP)

Supplementary Data file 4 (top 160 LR pairs by specificity and by magnitude)

## Supplementary Materials

**The PDF file includes**

Figs. S1 to S9 with Figure legends

Tables S1 to S2

Materials and Methods

**Other Supplementary Material for this manuscript includes the following**

Supplementary Data File 1

Supplementary Data File 2

Supplementary Data File 3

Supplementary Data File 4

**Figure S1.**
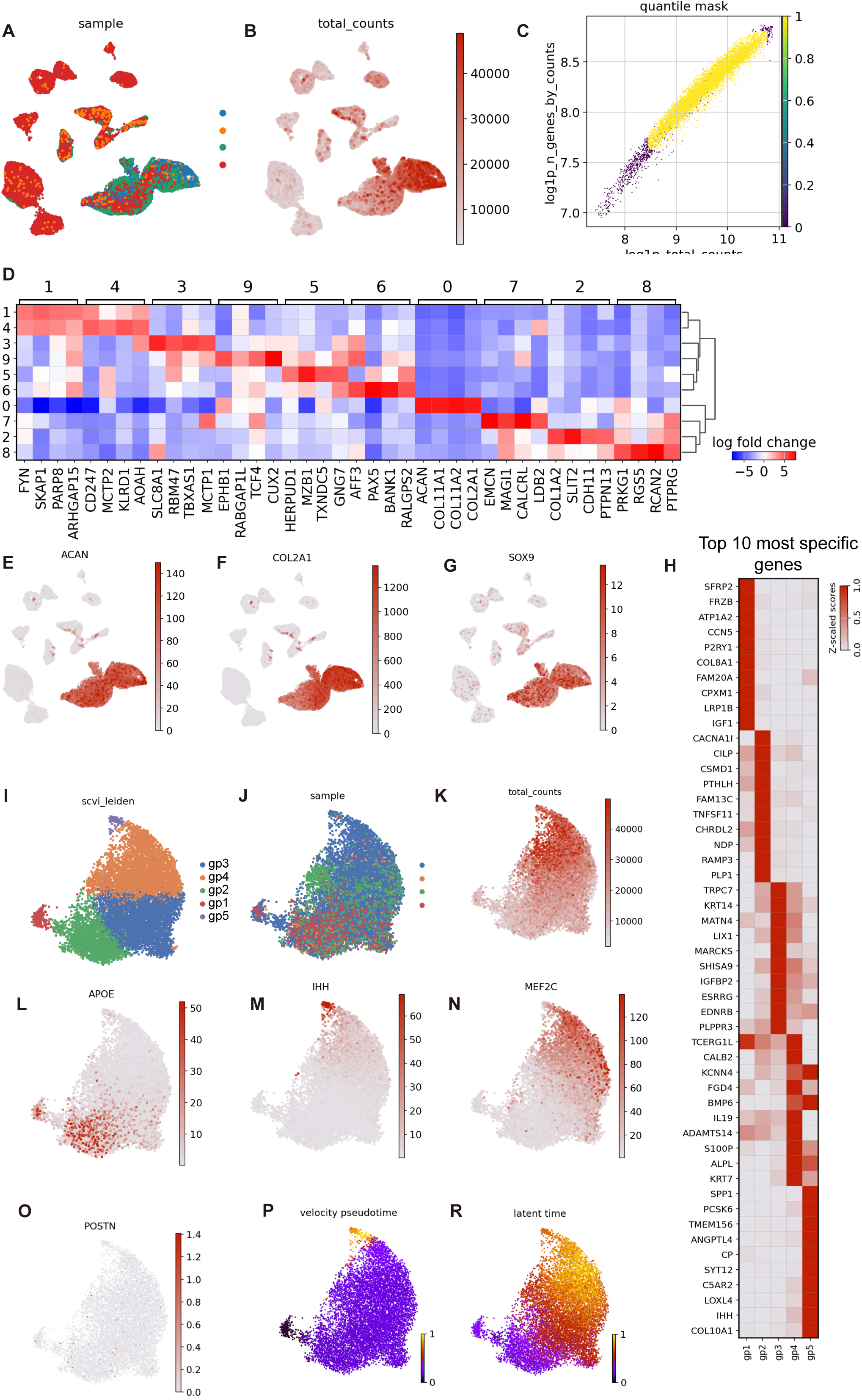
Cellular heterogeneity and molecular characteristics of the human growth plate. (A) UMAP clustering showing patient representation across clusters, with four patients color-coded to assess distribution and variability. (B) Total RNA transcripts per cell visualized on the UMAP plot, indicating transcript abundance across clusters. (C) Scatter plot of the number of genes versus total gene counts per cell on a logarithmic scale, highlighting transcriptional activity and sequencing quality. (D) Heatmap displaying the top 4 differentially expressed genes (DEGs) for each cell cluster in the UMAP embedding from Fig. 1D, providing insights into cluster-specific gene signatures. (E-G) Feature plots illustrating chondrocyte-enriched expression of (E) *ACAN* (Aggrecan), (F) *COL2A1*, and (G) *SOX9* mRNAs, key markers of cartilage development and maintenance. (H) Heatmap showing the expression of the top 10 DEGs across reclustered human chondrocytes. (I) UMAP clustering of human growth plate chondrocytes, reproduced from Fig. 1E for orientation, demonstrating subpopulation diversity. (J) Patient representation across UMAP clusters of human growth plate chondrocytes, showing contributions from each patient. (K) Total RNA transcripts per cell in sequenced chondrocytes, emphasizing transcript abundance across the dataset. (L-O) Feature plots displaying expression of (L) *APOE*, (M) *IHH*, (N) *MEF2C*, and (O) *Periostin* mRNAs in human growth plate chondrocytes, highlighting distinct spatial and functional expression patterns. (P, R) Pseudo-time (P) and latent time (R) predicted by VeloCITO plotted on UMAP embedding, revealing dynamic gene expression trajectories and temporal progression in chondrocyte differentiation.

**Figure S2.**
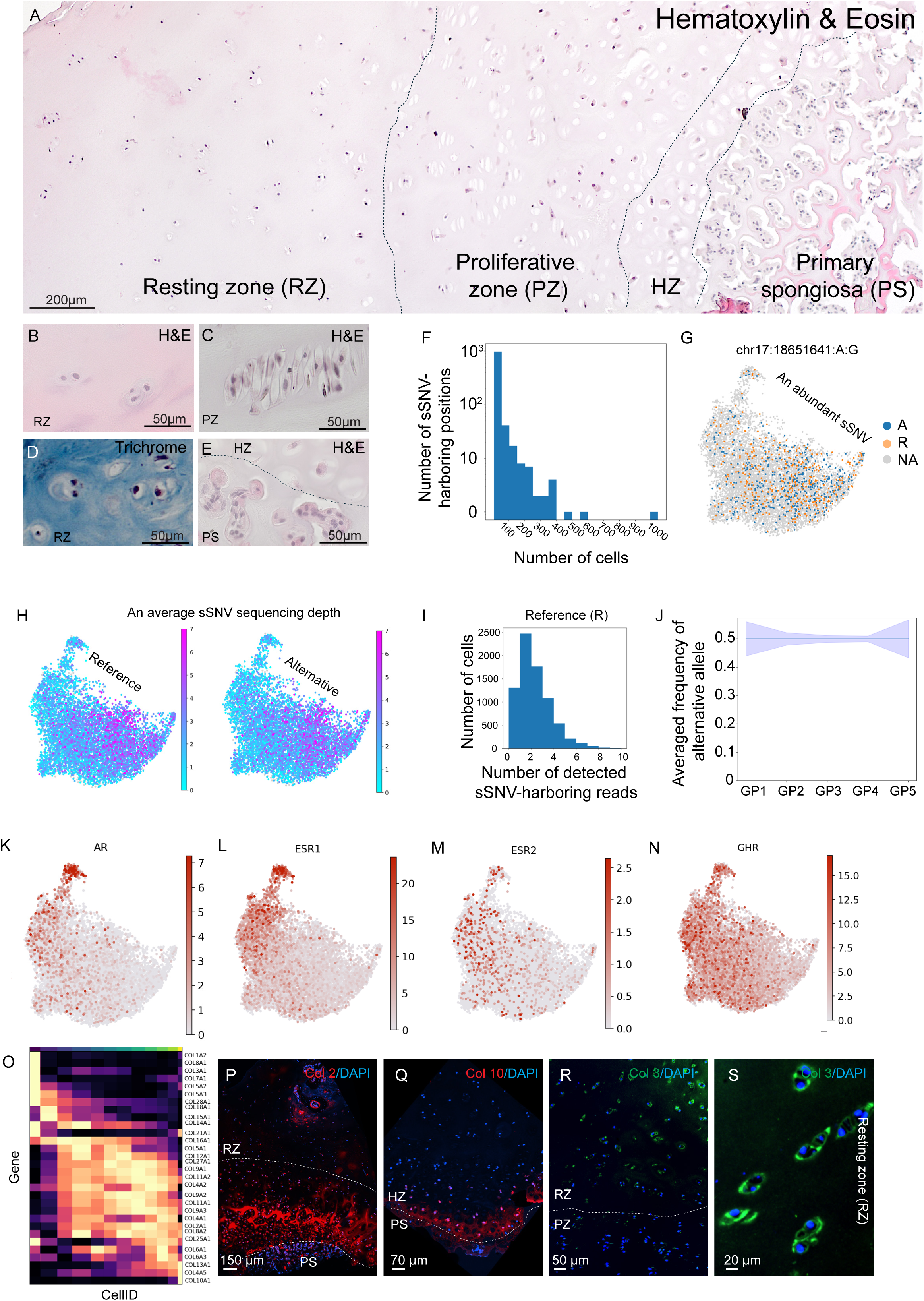
Further characterization of the pubertal human growth plate. (A-E) Representative histological images of human growth plates stained with Hematoxylin and Eosin (A, B, C, E) or Trichrome (D). Images include low magnification (A) and high magnification views of specific zones: (B, D) resting zone, (C) proliferative zone, and (E) hypertrophic zone. (F-J) Analysis of somatic single-nucleotide variants (sSNVs) with Monopogene. (F) Histogram showing the distribution of sSNVs per cell recovered in the analyzed dataset, highlighting genetic heterogeneity. (G) UMAP embedding illustrating the distribution of a single abundant sSNV from one patient throughout the entire UMAP embedding. Orange dots represent cells harboring reads with the reference nucleotide (adenine at position chr17:18651641), blue dots indicate cells harboring reads with the alternative nucleotide (guanine at this same position), and gray dots denote all other cells. (H) UMAP embeddings showing the total number of detected reference (left panel) and alternative (right panel) sSNVs per cell across the dataset. (I) Histogram showing the number of reads harboring the reference nucleotides per cell. (J) Relative abundance of alternative nucleotides per cluster normalized for sequencing depth, demonstrating no changes in the number of sSNVs across cellular populations. (K-N) Feature plots showing expression of mRNAs encoding receptors for pubertal hormones: (K) Androgen Receptor (*AR*), (L) Estrogen Receptor Alpha (*ESR1*), (M) Estrogen Receptor Beta (*ESR2*), and (N) Growth Hormone Receptor (*GHR*), indicating hormone signaling potential in the growth plate. (O) Heatmap displaying variation in collagen genes along the differentiation trajectory visualized in Fig. 2B. (P) Representative immunofluorescence image showing Collagen Type II (red) with nuclei counterstained by DAPI (blue). (Q) A representative image of Collagen Type X mRNA (red) visualized by Hybridization Chain Reaction (HCR), illustrating its spatial distribution. (R, S) HCR images of Collagen Type III mRNA (green), showing expression at the resting-to-proliferative zone border (R) and within the resting zone at high magnification (S). RZ – Resting Zone; PZ – Proliferative Zone; HZ – Hypertrophic Zone; PS – Primary Spongiosa.

**Figure S3.**
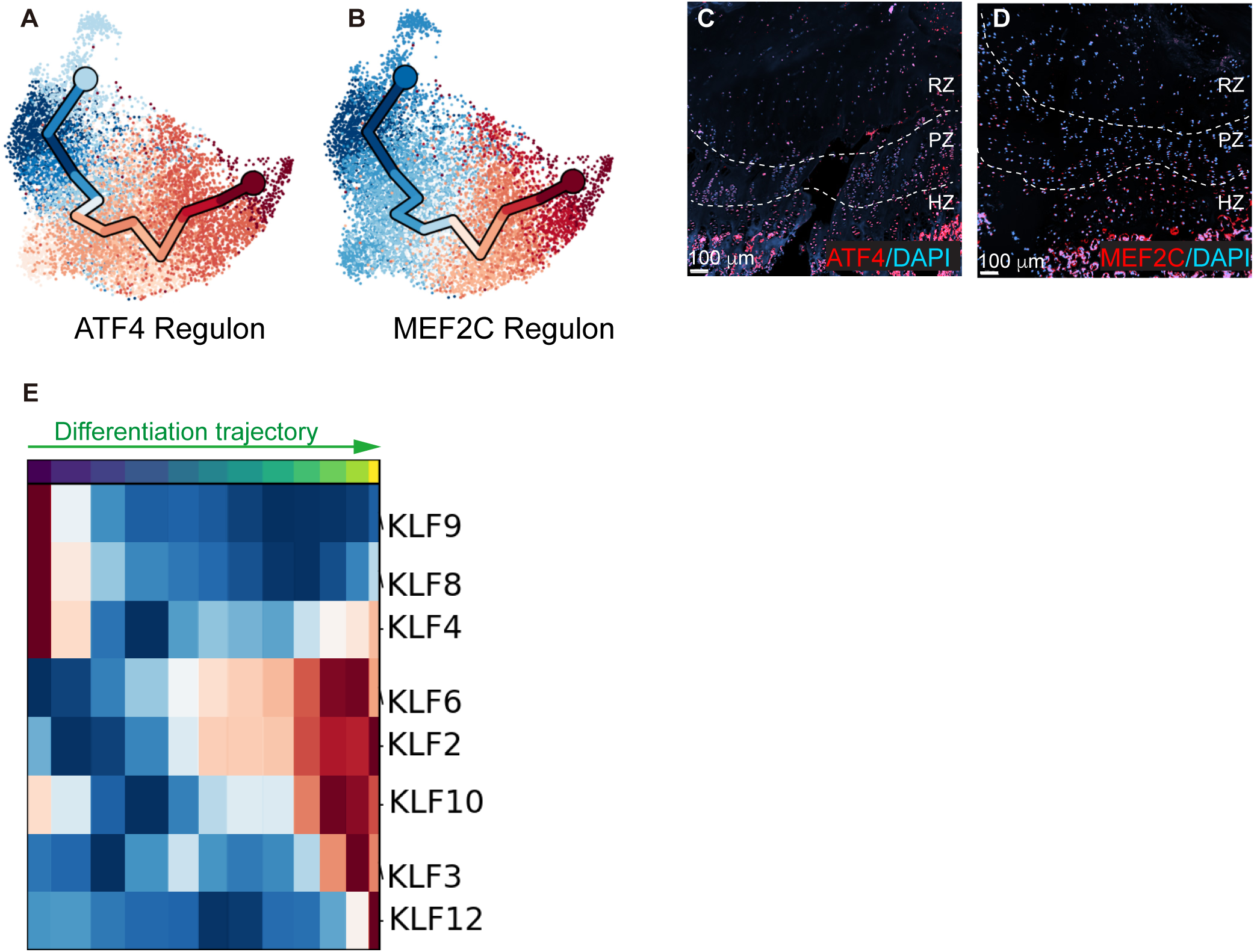
Examples and validation of selected regulons. (A, B) UMAP embeddings illustrating regulon activity of (A) *ATF4* and (B) *MEF2C* predicted by SCENIC. Regulon activity is shown along the differentiation trajectory defined in Fig. 2B, highlighting zone-specific activity patterns. (C, D) Representative images of immuno-detected expression of (C) ATF4 and (D) MEF2C in growth plate sections, showing protein localization corresponding to predicted regulon activity. Nuclei were counterstained with DAPI (blue). (E) Heatmap showing activity of transcription factors of the KLF family along the differentiation trajectory from Fig. 2B, emphasizing dynamic changes in KLF regulon activation across growth plate zones. RZ – Resting Zone; PZ – Proliferative Zone; HZ – Hypertrophic Zone; PS – Primary Spongiosa.

**Figure S4.**
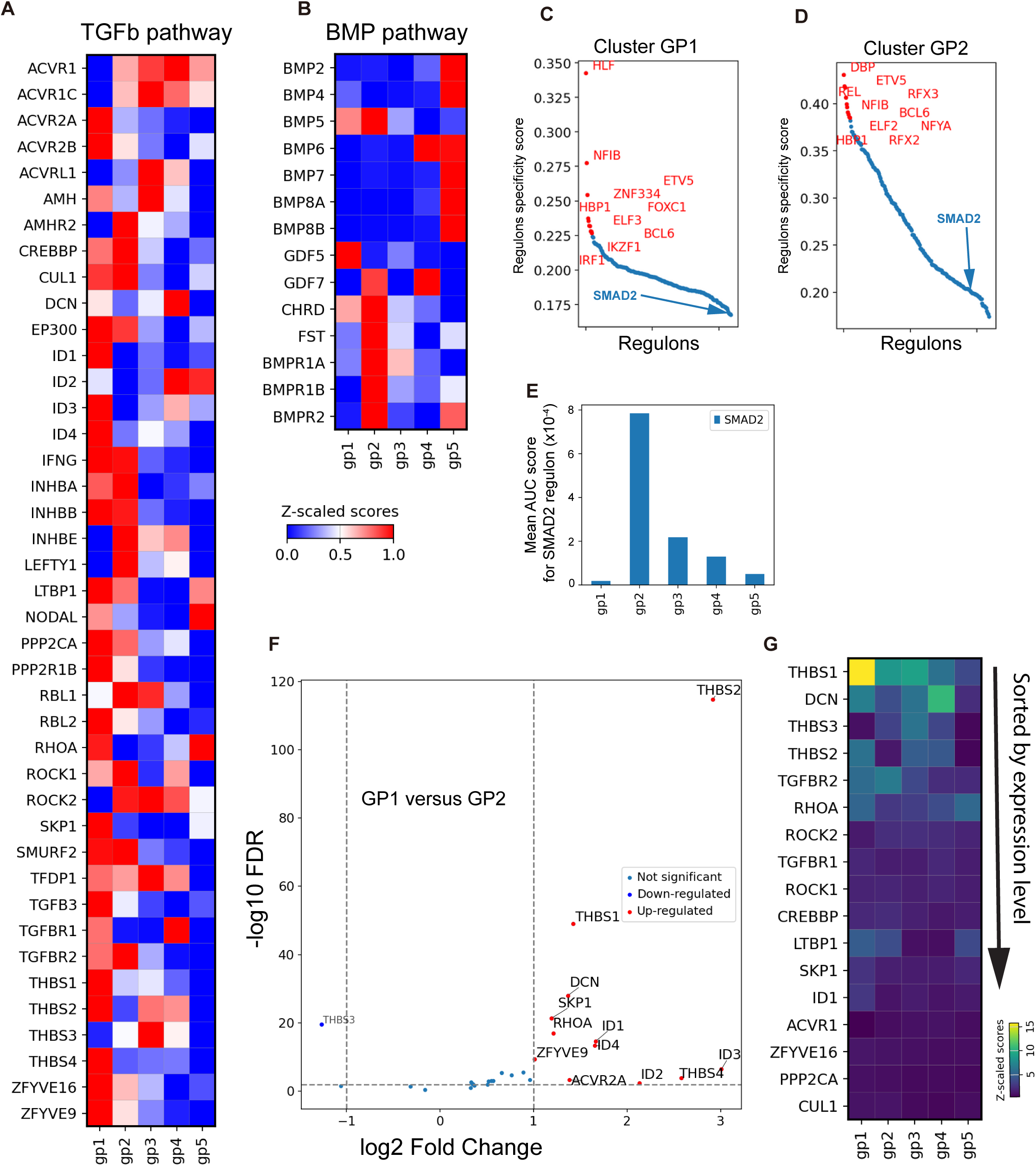
TGFβ pathway in cartilaginous stem cells. (A, B) Heatmaps showing scaled expression of genes related to the (A) TGFβ pathway and (B) BMP pathway across chondrocyte subsets. Gene expression is scaled between genes to facilitate comparisons of distribution across clusters. (C, D) Scatter plots ranking specificity of regulons predicted by SCENIC in the (C) GP1 cluster and (D) GP2 cluster. (E) Histogram depicting the mean AUC activity score of the *SMAD2* transcription factor predicted by SCENIC across chondrocyte subsets, revealing differences in pathway activity. (F) Volcano plot comparing TGFβ pathway genes from (A) between GP1 and GP2 clusters. Red dots indicate genes significantly elevated in GP1, while a single gene, *THBS3* (blue dot), is significantly elevated in GP2. (G) Heatmap displaying the most highly expressed genes from (A) without scaling between genes, providing a direct comparison of relative expression levels within the TGFβ pathway family.

**Figure S5.**
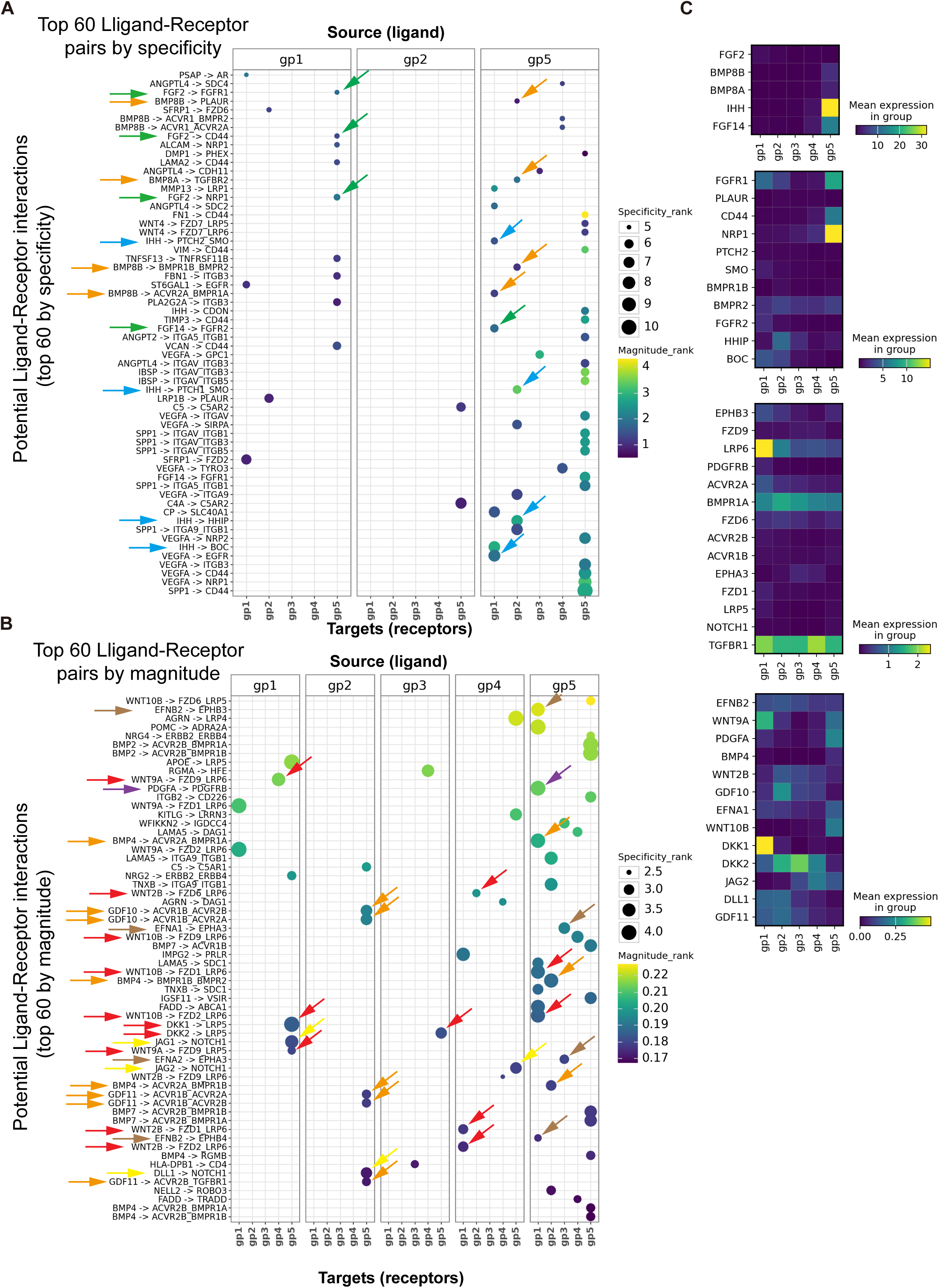
Predicted ligand-receptor interactions between different clusters within the human growth plate. (A, B) Bubble plots showing the top 60 predicted ligand-receptor pairs across clusters, ranked by (A) specificity and (B) magnitude. Clusters GP3 and GP4 do not include predicted interactions among the top 60 ligand-receptor pairs based on specificity and are omitted in A. Bubble size represents the specificity of the interaction, and bubble color indicates the computed strength of interaction. Arrows indicate cluster-separated ligand-receptor pairs within the following signaling pathways: Hedgehog (blue), BMP/TGFβ (orange), Wnt (red), FGF (green), Ephrin (brown), PDGF (violet), and Notch (yellow). (C) Heatmaps showing RNA expression of selected ligand-receptor pairs across chondrocyte subsets, without scaling between genes, and the heatmap is divided into four sections according to the relative expression scale for visualization purposes.

**Figure S6.**
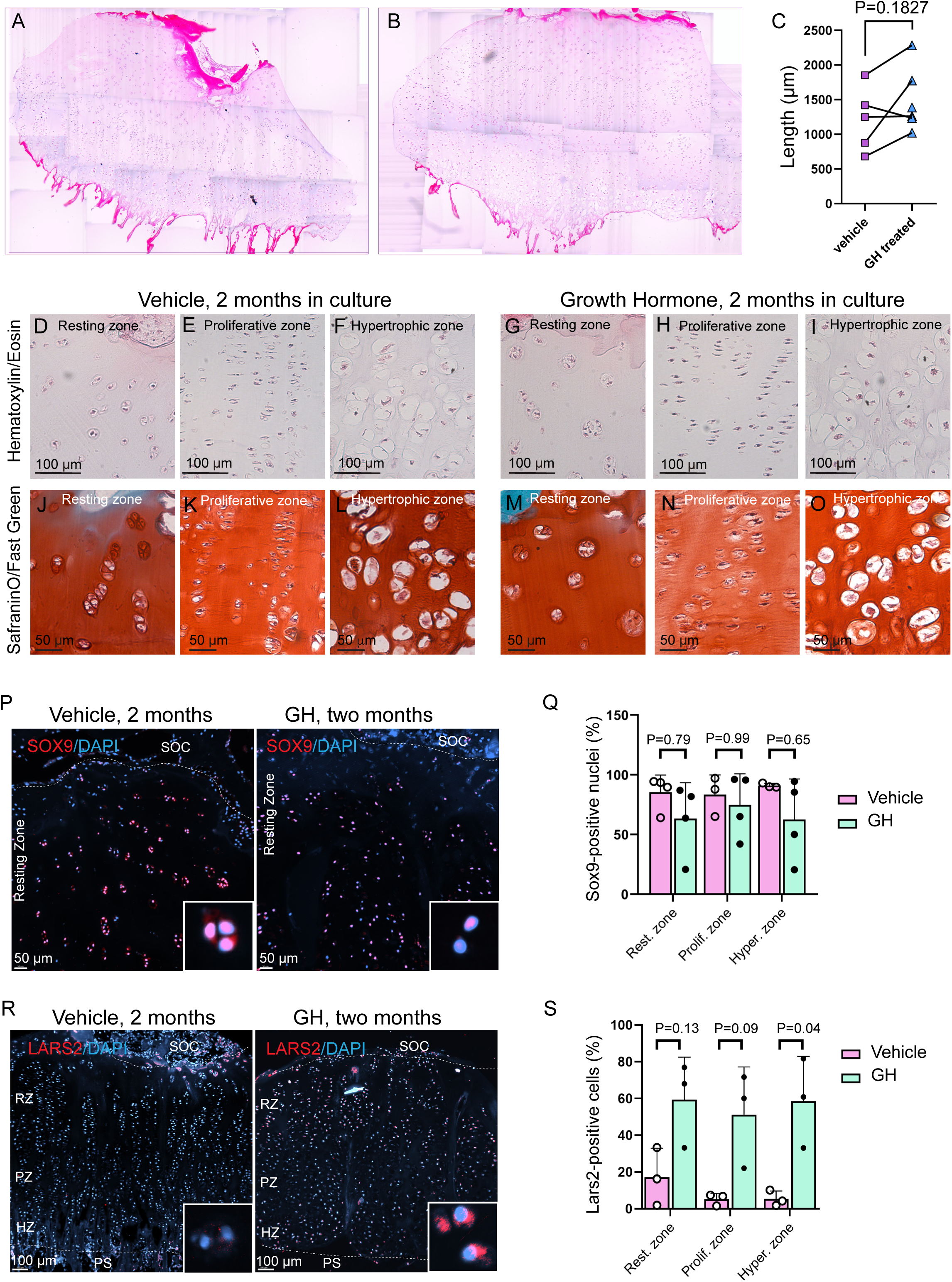
The effect of long-term GH treatment on the human growth plate. (A, B) Explants of adolescent human growth plates cultured for two months in the presence of (A) vehicle or (B) growth hormone (GH). Tile scans of entire growth plate sections stained with Hematoxylin and Eosin show no morphological malformations in either group. (C) Quantification of growth plate height, measured from the secondary ossification center (SOC) to the primary spongiosa, for vehicle- and GH-treated explants (n = 5 patients). (D-O) High-magnification histological images of indicated growth plate zones from sections stained with Hematoxylin and Eosin (D-I) or Safranin-O/Fast Green (J-O). Images show vehicle-treated explants (D-F, J-L) and GH-treated explants (G-I, M-O). (P, Q) Immunofluorescence staining of SOX9 (red) and quantification of SOX9-positive nuclei in each growth plate zone for vehicle- and GH-treated groups (n = 4 patients/group). (R, S) Immunofluorescence staining of LARS2 (red) and quantification of LARS2-positive cells in vehicle- and GH-treated groups (n = 3 patients/group). Statistical analyses were performed using a paired Student’s *t*-test for (C) and ONE-WAY-ANOVA with Turkey’s correction for multiple comparisons for (Q, S). SOC – Secondary Ossification Center; RZ – Resting Zone; PZ – Proliferative Zone; HZ – Hypertrophic Zone; PS – Primary Spongiosa.

**Figure S7.**
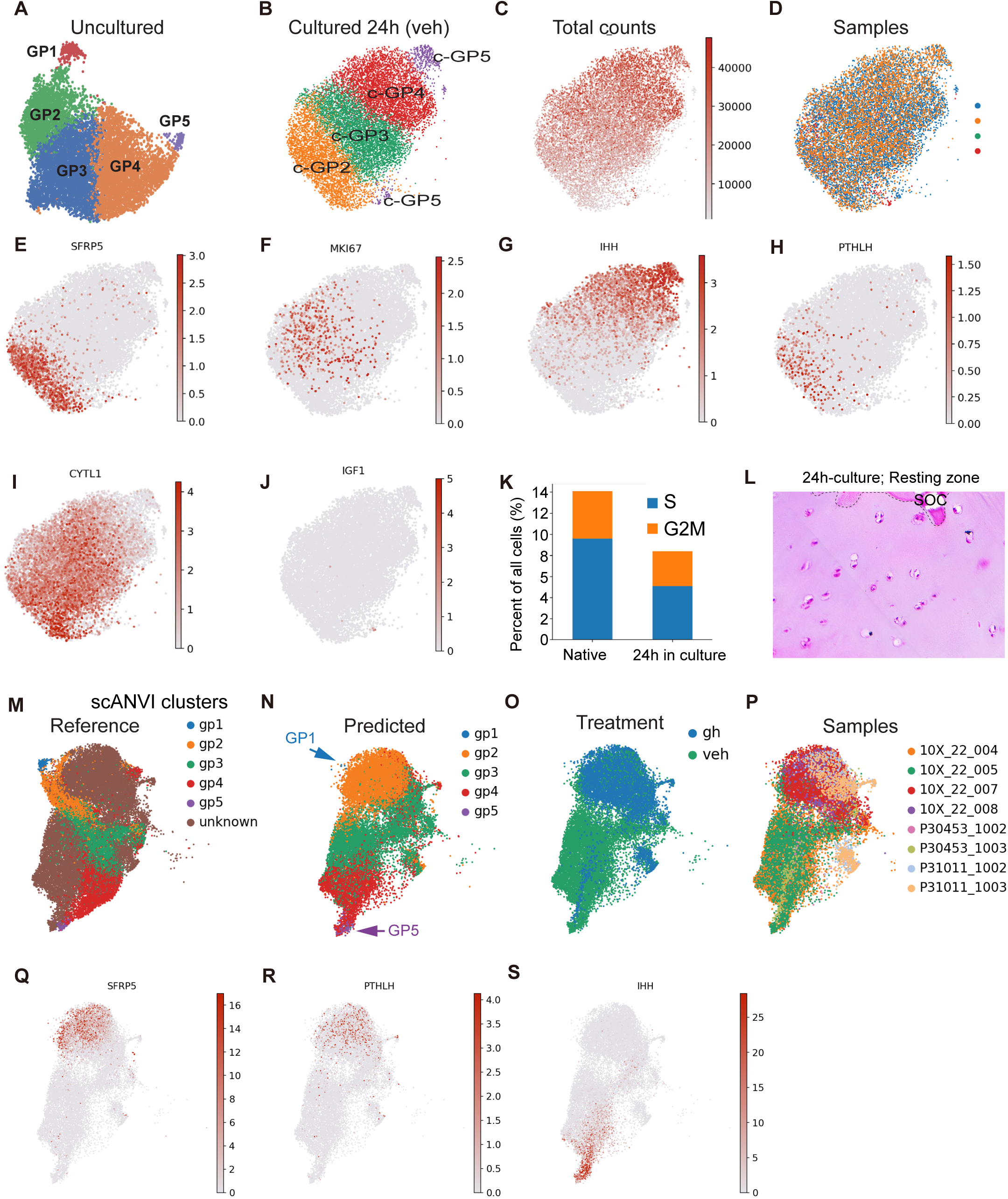
Validation and molecular characterization of short-term culture model. (A, B) UMAP clustering of (A) uncultured human growth plate chondrocytes, reproduced from Fig. 1E, and (B) chondrocytes obtained from growth plate explants cultured for 24 hours, for direct comparison of cellular distributions. (C, D) Feature plots showing (C) total RNA counts in explant cultures and (D) biological replicates color-coded for visualization. (E-J) Feature plots illustrating mRNA expression of (E) *SFRP5*, (F) *MKI67*, (G) *IHH*, (H) *PTHLH*, (I) *CYTL1*, and (J) *IGF1* in explant cultures. (K) Bar graph comparing the relative percentage of cells in S phase (blue) and G2M phase (orange) between naïve (uncultured) and short-term cultured chondrocytes. (L) Representative H&E-stained image of sections from short-term cultured explants, showing preserved cell morphology. (M) scANVI embedding combining known clusters from uncultured chondrocytes (color-coded GP1-GP5) with cells from cultured explants (unknown, brown), including both vehicle- and GH-treated samples. (N, O) scANVI predicted clusters for cultured explants with (O) treatment representation and (P) biological replicates. (Q-S) Feature plots showing mRNA expression of (Q) *SFRP5*, (R) *PTHLH*, and (S) *IHH* in combined cultured vehicle- and GH-treated samples.

**Figure S8.**
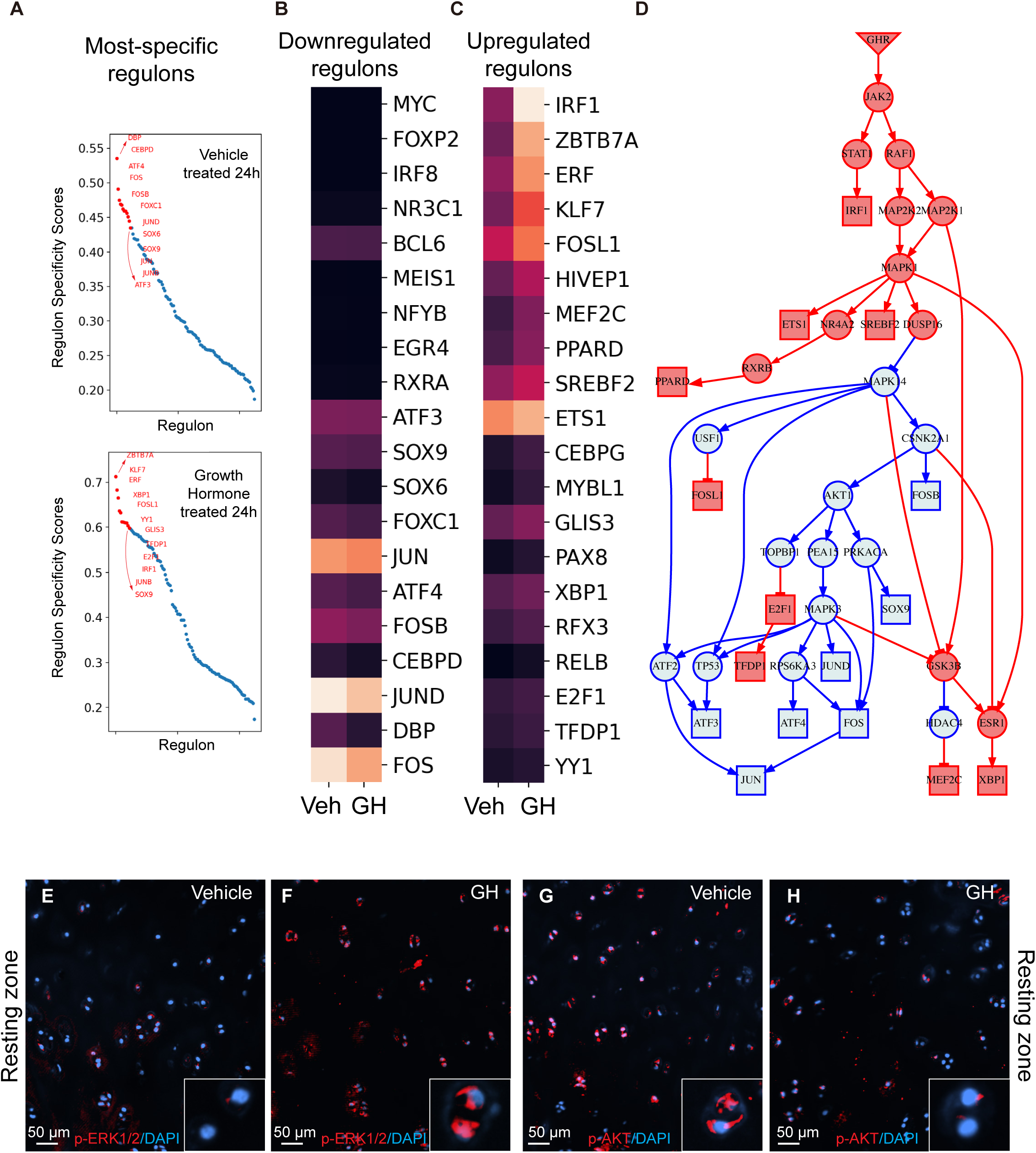
Transcription factors and intracellular signaling pathways activated by GH within 24 hours. (A) Scatter plots ranking the specificity of regulons within the cultured GP2 cluster predicted by SCENIC for vehicle- and GH-treated explants. (B, C) Heatmaps displaying the (B) most downregulated and (C) most upregulated regulons by GH in the GP2 cluster of cultured explants. The color code represents average regulon activity scores in relative units, with transcription factors (TFs) sorted by the absolute difference between control and treatment groups. (D) Inferred protein-protein interaction network linking activated GHR with regulons identified in (B) and (C). The inverted triangle denotes the known activation of GHR, squares represent known transcription factors identified in B and C, and circles indicate predicted intermediate proteins. Red denotes activation, and blue denotes inhibition. (E-H) Immunofluorescence staining of (E, F) phosphorylated extracellular signal-regulated kinases (p- ERK1/2) and (G, H) phospho-AKT in (E, G) vehicle-treated and (F, H) GH-treated explants. Images focus on the resting zone.

**Figure S9.**
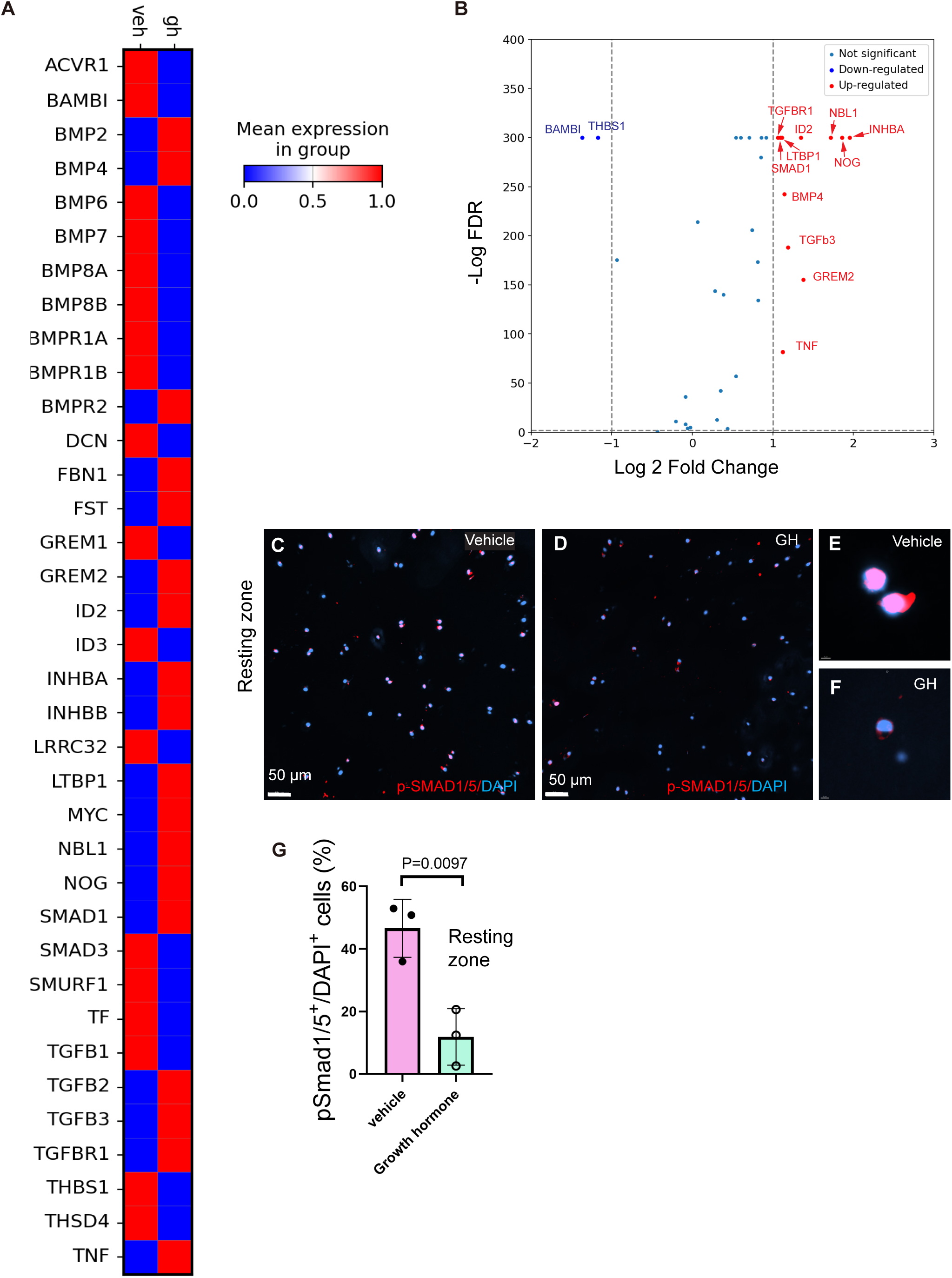
Regulation of TGFβ pathway by GH. (A) Heatmap showing the expression of genes belonging to the TGFβ/BMP pathways in the GP2 cluster from vehicle- and GH-treated growth plate explants after 24 hours of treatment. (B) Volcano plot depicting significant differences in gene expression for genes shown in (A) between vehicle and GH-treated samples. Red dots indicate upregulated genes, and blue dots indicate downregulated genes. FDR – False Discovery Rate. (C, D) Immunofluorescence staining of phospho-SMAD1/5 (p-SMAD1/5, red) in (C) vehicle- and (D) GH-treated samples, demonstrating signal distribution. (E, F) Magnified images of p-SMAD1/5 staining from (C) and (D), highlighting nuclear localization. (G) Quantification of p-SMAD1/5-positive nuclei in the resting zone of vehicle-treated (pink) and GH-treated (green) samples. Data represent measurements from 3 patients per group. Statistical analyses were performed using an unpaired Student’s *t*-test.

**Table S1.**
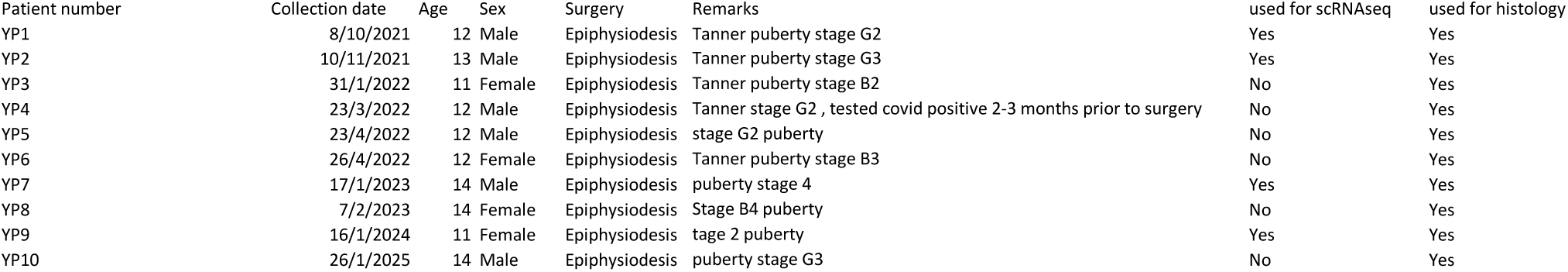
Patient information.

**Table S2.**
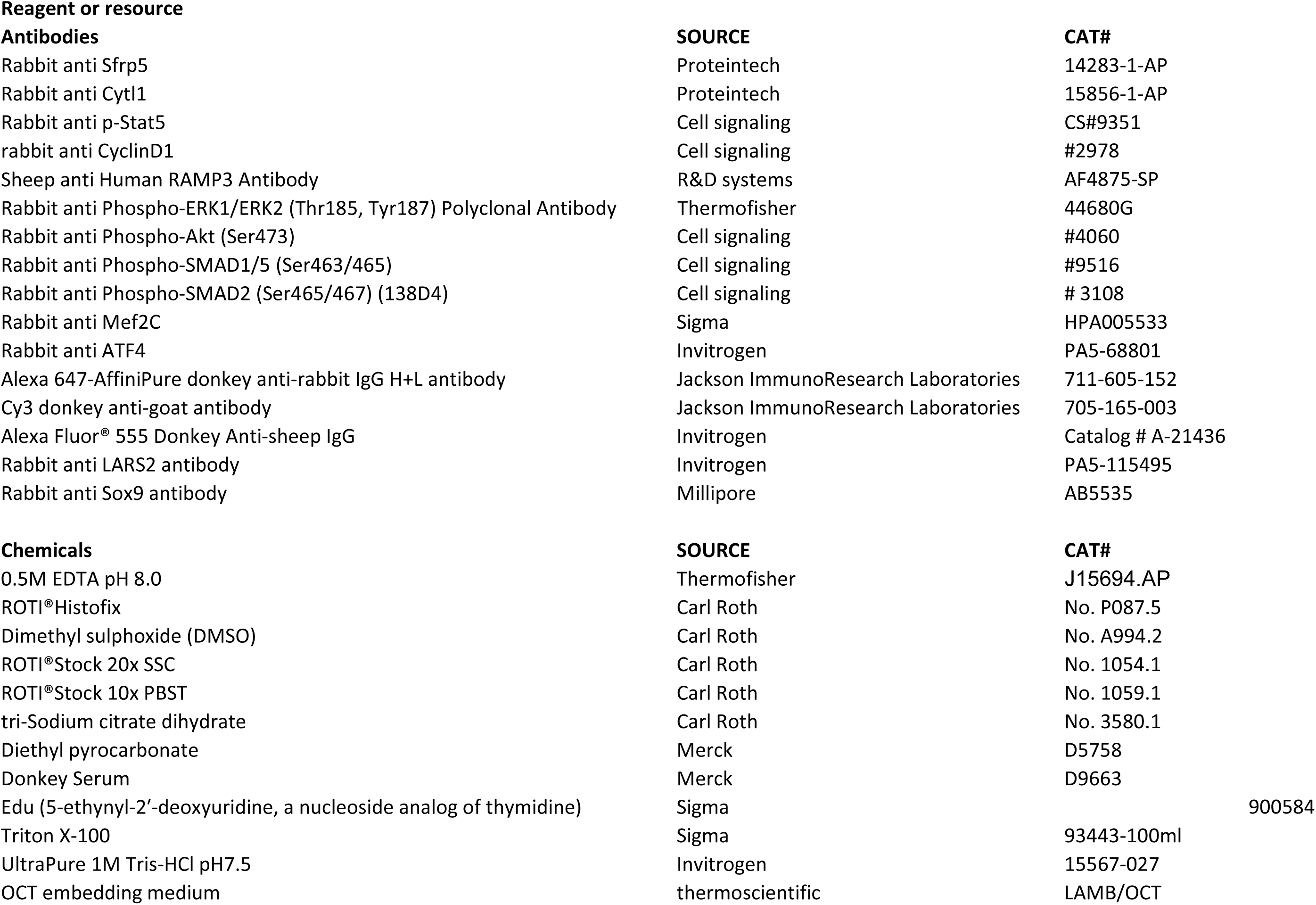
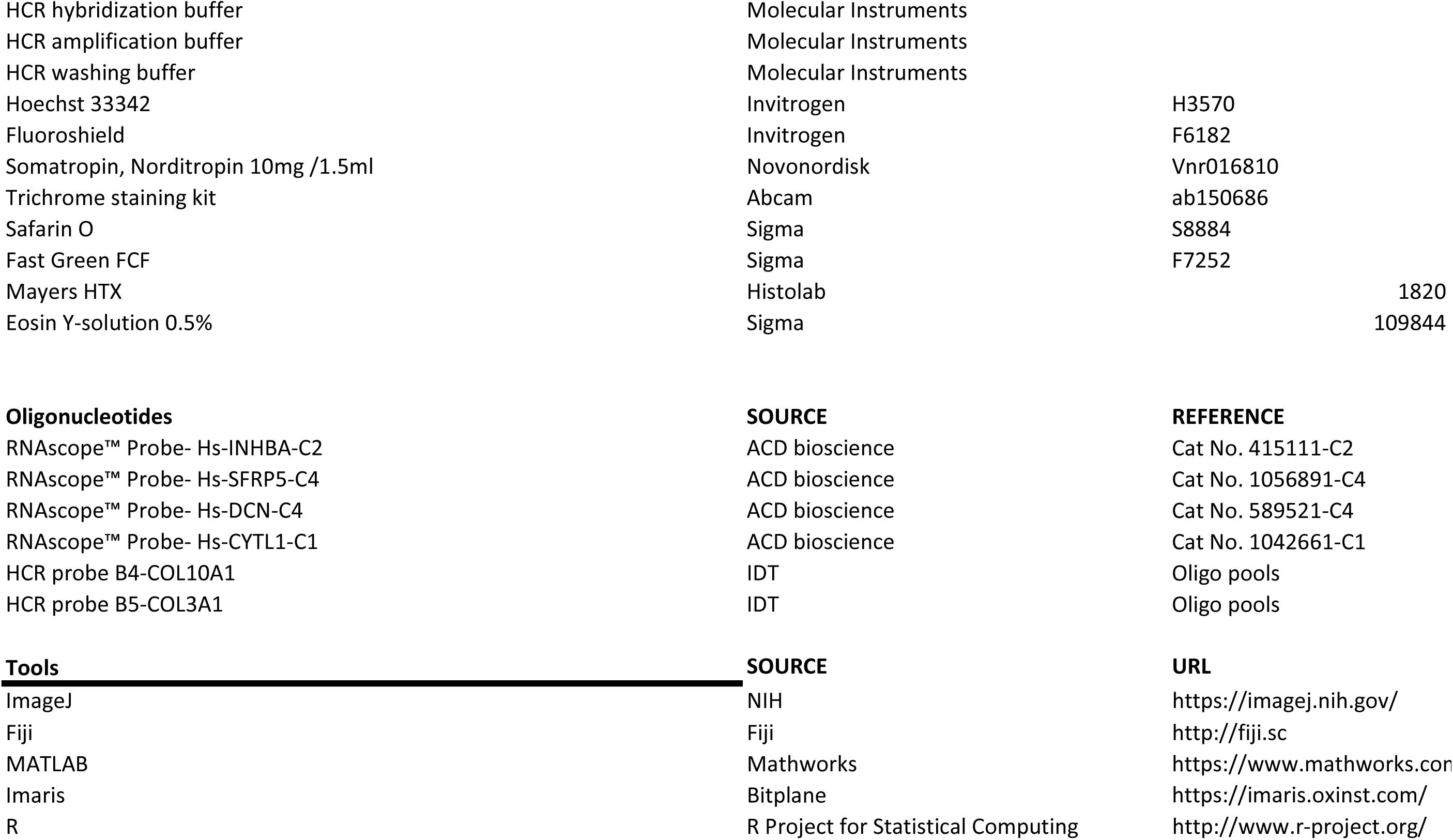
Source of Reagents.

## Materials and Methods

### Study Design

### Sample Size

Due to the scarcity of fresh adolescent human growth plate donors, only four out of ten patients were sequenced using the 10X Chromium platform. Other samples were used for long- and short-term experiments and for validation purposes. Samples from at least three patients were used for quantified histological analysis and validation experiments.

### Data Inclusion/Exclusion Criteria

No data were excluded from statistical tests.

### Outliers

No outliers were excluded from the analysis.

### Research Objectives

The primary objective of this study was to investigate whether pubertal human growth plate chondrocytes can directly respond to growth hormone treatment. Additionally, the study aimed to identify the cellular and molecular targets of growth hormone. This objective further motivated an extensive characterization of the native pubertal human growth plate, as it had not been comprehensively characterized in previous studies.

### Research Subjects

The surgical removal of the growth plates, known as epiphysiodesis, was performed on healthy adolescent children with predicted tall stature in accordance with local pediatric guidelines. The study included ten subjects aged 11–14 years, encompassing both sexes. Detailed patient information is provided in **Table S1**. All patients were monitored, and all surgeries were performed at Karolinska University Hospital, Sweden. The collection of human growth plate biopsies post-surgery was approved by the local human ethics committee (Karolinska Institutet Research Ethics Committee North, Karolinska University Hospital, Stockholm, Sweden). Informed consent was obtained from all patients and their parents, and the process was documented in hospital records.

### Experimental Design

#### Human Growth Plate Biopsies

Human growth plate biopsies were harvested during epiphysiodesis surgery by an experienced orthopedic surgeon under live X-ray guidance. The biopsies were immediately transported on ice to the laboratory for subsequent analysis. Under sterile conditions and on ice, the biopsies were sliced perpendicularly to their long axis into 1–2 mm sections to obtain slices encompassing the secondary ossification center (SOC), the growth plate, and the primary spongiosa. The slices were then allocated for either immediate analysis (fixed or digested for single-cell RNA sequencing [scRNA-seq]) or for explant culture.

#### Preparing Cells for single-cell RNA sequencing (scRNA-seq)

The obtained slices were dissected from bone under a stereo microscope, ground into small fragments, transferred to fresh, ice-cold HBSS, and thoroughly pipetted to remove residual blood. Then HBSS was replaced with a digestion solution comprising Collagenase P (3 U/mL, Roche) and TrypLE™ Select Enzyme (1X, Thermo Fisher Scientific, Catalog No: A1217701) in HBSS. The tubes were incubated for 30–40 minutes at 37°C with shaking at 120 rpm. During digestion, the tissue fragments were disrupted every 10 minutes using a 1-mL pipette. The digestion was terminated by the addition of 10 mL of 2% FBS in cold HBSS. The mixture was centrifuged at 300 rpm for 10 minutes at 4°C. The supernatant was carefully removed, and the pellet was resuspended in 100 µL of 2% FBS in 1X PBS. Ice-cold methanol (400 µL) was added dropwise to the suspension, and the sample was placed on ice for 10 minutes. The cells were stored at -80°C for a maximum of six months prior to library preparation.

#### Human Growth Plate Explant Culture

For organ culture experiments, the explants were incubated at 37°C with 5% carbon dioxide in a culture medium consisting of DMEM/F12 (without phenol red), gentamycin (20 µg/ml), 0.2% bovine serum albumin (BSA), β-glycerophosphate (1 mM), ascorbic acid (5 µg/ml), and 2% fetal bovine serum (FBS), as previously described (*1, 2*).

*Long-term cultures:* To provide proof-of-principle evidence of direct growth hormone (GH) action, the slices were cultured in medium supplemented with or without 40 ng/ml of human recombinant GH (Norditropin, Novo Nordisk) for 2 months. The medium was replenished every 2–3 days using 3 ml per well in a 6-well dish. At the end of the culture period, the slices were washed with ice-cold 1X PBS, fixed overnight in ROTI®Histofix (Carl Roth, Art. No. 8907.1), and processed for histological analysis.

*Short-term cultures:* To investigate the mechanisms underlying GH action, human growth plate slices were cultured for 24 hours with or without 40 ng/ml of GH (Norditropin, Novo Nordisk). Following the culture period, the slices were washed with ice-cold PBS, dissected from bone under a stereo microscope, and either subjected to single-cell preparation or fixed for histological analysis as described above.

### Methods

#### Preprocessing for Single-Cell Library Preparation

Prior to single-cell library preparation, frozen cell suspensions were thawed from -80°C. The suspension was pelleted at 1000 rpm, resuspended, and washed twice with 3X SSCT in 2% FBS containing 50 U of RNaseOUT™ Recombinant Ribonuclease Inhibitor (Thermo Fisher Scientific, Catalog No: 10777019) to remove methanol. The suspension was filtered through 70 µm Cell-Strainer Caps to remove extracellular matrix debris, centrifuged at 1000 rpm, and resuspended in 1X PBS with 3% BSA. For purification, the cell suspension was sorted using a BD FACS Aria Fusion with a 100 µm nozzle (specific for human cells). Debris was gated out based on FSC-H versus SSC-H parameters. Cell viability and density were quantified using the Thermo Scientific Invitrogen Countess II Automated Cell Counter (Catalog No: AMQAX1000R) according to the manufacturer’s instructions. The processed cells were immediately used for single-cell library preparation.

#### 10X Genomics scRNA-seq

Single-cell libraries were prepared following the manufacturer’s protocol using the Chromium Next GEM Single Cell 3’ Kit v3.1 (Cat. No. 1000269) and the Chromium Next GEM Chip G Single Cell Kit (Cat. No. 1000127). Samples where emulsion preparation failed were excluded from this study. Uncultured, vehicle-treated, and growth hormone-treated cells were separately indexed and pooled for sequencing on the Illumina NovaSeq 6000 or NovaSeq X platform at a depth of 20,000 reads per cell. Quality control of the libraries was performed using a Bioanalyzer to ensure library integrity and sequencing readiness.

#### Tissue Processing

Fixed samples were processed for either paraffin or frozen sectioning. Paraffin embedding and sectioning were performed standardly. For frozen sections, tissues were dehydrated using a methanol gradient (Methanol Puriss 99.85%, Histolab, Cat. No. 02095) at concentrations of 20%, 40%, 60%, 80%, and 100%. Each step involved incubation for 1 hour at 4°C with agitation. After dehydration, tissues were delipidated using dichloromethane (Merck, Cat. No. D65100) for 3 hours at 4°C. This was followed by rehydration through a reverse methanol gradient (100%, 80%, 40%) and incubation in 2X SSCT (ROTI®Stock 20x SSC, Carl Roth, Cat. No. 1054.1) for 1 hour per step at 4°C. Subsequently, the tissues were decalcified in 10% EDTA (Merck, Cat. No. E6511) for two nights at 4°C with gentle agitation, replacing the solution daily. Decalcified tissues were then incubated overnight in 30% sucrose (Merck, Cat. No. S0389) solution prepared in 1X PBS until the tissues sank to the bottom of the container. Finally, the processed human growth plate samples were embedded in OCT compound (Tissue-Tek® O.C.T. Compound, Sakura, Cat. No. 4583), frozen on dry ice, and stored until sectioning.

#### Multiplex Fluorescence In Situ Hybridization with RNAscope

Multiplex fluorescence in situ hybridization (FISH) was performed using the RNAscope Multiplex Fluorescent V2 Assay (ACD; Cat. No. 323100) and the RNAscope® 4-plex Ancillary Kit (ACD; Cat. No. 323120) according to the manufacturer’s guidelines, with minor modifications as described below. Tissue sections were rehydrated, treated with hydrogen peroxide supplied in the kit, and heated in a target retrieval buffer. The sections were then digested with pepsin before the application of FISH target probes. The probes were incubated overnight at 40°C. The following probes were used:

Hs-CYTL1 (ACD; Cat. No. 1042661-C1), Hs-INHBA (ACD; Cat. No. 415111-C2), Hs-SFRP5 (ACD; Cat. No. 1056891-C4), Hs-DCN (ACD; Cat. No. 589521-C4). Signal development was achieved using TSA Plus fluorophore kits (PerkinElmer; Cat. Nos. NEL744001KT and NEL745001KT) at a dilution of 1:800, followed by counterstaining with DAPI. Fluorescence signals were detected and visualized using a Nikon spinning disk confocal microscope.

#### Multiplex Fluorescence In Situ Hybridization with Hybridization Chain Reaction (HCR)

Detection of Collagen mRNA transcripts was performed using Hybridization Chain Reaction (HCR) according to the manufacturer’s protocol (Molecular Instruments, USA). HCR was used instead of RNAscope for abundant transcripts due to economic reasons. Briefly, tissue sections were dehydrated using an ethanol gradient and digested with Proteinase K to unmask mRNA transcripts. Pre-hybridization was performed, followed by hybridization with 2 µL of oligo pool probes in hybridization buffer (Molecular Instruments, USA) at 37°C overnight. After hybridization, probes were washed three times with probe wash buffer, each for 5 minutes. Amplifier pairs were prepared by heating in separate PCR tubes to 95°C in a thermocycler and cooling to 25°C for 30 minutes. Amplification was carried out using HCR amplifiers in amplification buffer (Molecular Instruments, USA) overnight at room temperature. Following amplification, tissue sections were washed four times with 5X SSCT for 10 minutes each. The sections were then mounted in Fluoroshield containing Hoechst 34580 for nuclear counterstaining.

#### Histological Staining: Hematoxylin, Eosin, Safranin-O, Fast Green, and Trichrome

Hydrated tissue sections were first washed with 1X PBS, followed by distilled water. The sections were stained with Mayer’s Hematoxylin (Histolab) for 1 minute, then washed three times with distilled water. The sections were developed to a bluish color using 1X PBS. After hematoxylin staining, the sections were counterstained with Eosin G (Histolab) for 30 seconds, dehydrated through an ethanol series, and mounted using Depex (Sigma, Cat. No. 06522).

For Safranin-O and Fast Green staining, hydrated sections were washed with distilled water and stained with Weigert’s Iron Hematoxylin for 5 minutes to label nuclei. The sections were then rinsed four times with water, followed by a quick dip (2 seconds) in Acid-Alcohol solution (1% HCl in 70% ethanol). Next, the sections were stained with 0.02% Fast Green for 1 minute, followed by immersion in 1% acetic acid (prepared in 70% ethanol) for 30 seconds. The sections were then stained with 1% Safranin-O for 10 minutes to visualize cartilage. Finally, the sections were dehydrated through absolute ethanol, cleared with HistoClear (Histolab), and mounted.

Trichrome staining was performed using the Trichrome Staining Kit (Abcam, Cat. No. ab150686) according to the manufacturer’s instructions.

#### Immunofluorescence Staining

*Phospho-Protein Immunostaining:* Tissue sections (6 µm) were thawed and dried at room temperature for 30 minutes. Slides were then incubated in 1X PBS twice, for 5 minutes each, to remove residual OCT compound. Antigen unmasking was performed using Antigen Unmasking Solution, Citrate-Based (Vector Labs, Cat. No. H-3300-250) at 55°C overnight, which was used instead of boiling to prevent section detachment. Following antigen unmasking, citrate buffer was washed off with 1X PBS. Hydrophobic barriers were created around the tissue sections using a hydrophobic barrier pen. The sections were blocked with 2.5% Normal Horse Serum (ImmPRESS® HRP Anti-Rabbit IgG Polymer Detection Kit, Peroxidase, Vector Labs, Cat. No. MP-7401). Primary antibodies targeting specific antigens (rabbit-hosted) were incubated overnight at 4°C in 3% BSA prepared in 1X TBS. After incubation, the sections were washed three times in 1X TBS, each wash lasting 3 minutes. The sections were subsequently incubated with ImmPRESS Peroxidase Polymer Anti-Rabbit IgG Reagent for 1 hour at room temperature, followed by three 3-minute washes in 1X TBS. Fluorescent signals were developed using the TSA Cyanine 5 System (Akoya Biosciences, Cat. No. NEL705A001KT) according to the manufacturer’s instructions. Sections were mounted with Fluoroshield containing Hoechst 34580 (Thermo Fisher Scientific, Cat. No. H21486) for nuclear fluorescence visualization.

*Non-Phospho Protein Immunolabeling:* For immunolabeling of non-phospho proteins, sections were washed with 1X PBS and blocked using 3% BSA in 1X TBS (blocking buffer). Primary antibodies, diluted in blocking buffer, were applied to the sections and incubated overnight at 4°C. After primary antibody incubation, the sections were washed three times in 1X PBS for 3 minutes each. Secondary antibodies conjugated with fluorescence and matching the host species of the primary antibodies were applied to the sections and incubated for 1 hour at room temperature. Afterward, the sections were washed three times in 1X PBS for 3 minutes each at room temperature. Finally, the sections were mounted using Fluoroshield containing Hoechst 34580 for nuclear fluorescence.

Details of the antibodies and their sources used in this manuscript are provided in **Table S2**.

#### EdU (5-Ethynyl-2-deoxyuridine) Labelling for Proliferating Cells in Organ Cultures

EdU, a thymidine analog that incorporates into DNA during replication, was used to label proliferating cells in human growth plate (hGP) explants. EdU was added to the hGP explant culture medium at a concentration of 10 mM, 24 hours prior to tissue collection to label proliferating cells. For EdU detection on tissue sections, the sections were permeabilized using 0.5% Triton X-100 in 1X PBS for 20 minutes, followed by two washes with 3% BSA in 1X PBS. The EdU labeling reaction mixture was prepared as follows: 100 µL of 1 M Tris (pH 7.5), 20 µL of 100 mM CuSO₄, 0.1 µL of Alexa-azide 647, 680 µL of ddH₂O. Immediately before initiating the reaction, 200 µL of 0.5 M ascorbic acid was added to the mixture. The reaction was developed on the sections for 30 minutes at room temperature. Following the reaction, the sections were washed twice with 3% BSA in 1X PBS for 5 minutes each. Finally, the sections were mounted using Fluoroshield containing Hoechst 34580, and images were captured using a confocal microscope.

#### Image Acquisition and Processing

Fluorescence images were acquired using a Nikon Ti2-E inverted microscope equipped with a CrEST X-light V3 spinning disk. Fluorophores, including Hoechst 34580, Alexa Fluor 488, 546, 647, Cy3, and Cy5, were excited with 405, 477, 520, 546, and 638 nm lasers, respectively. Images were captured using a 20x Nikon CFI60 Plan Apochromat Lambda objective. Z-stack images were taken at 4 µm intervals, covering a depth of 20–40 µm, with tissue sections having a thickness of 6 µm. Tile images were automatically stitched during acquisition and exported in ND2 file format. For image processing and visualization, ND2 files were converted to IMARIS format (IMS) using Imaris Converter 10.0. Images intended for the manuscript were subsequently exported as TIF files.

#### Bioinformatic Analysis

scRNA-seq FASTQ reads were initially processed using CellRanger. To separate nascent and mature mRNA reads, Velocyto was employed. The output matrices from Velocyto were analyzed using the Scanpy Python package. Only protein-coding genes were retained for this stage. Reads were filtered according to quality control (QC) parameters, normalized for sequencing depth, log-transformed, and scaled. Potential doublets were removed using Scrublet, and highly variable genes were identified. Dimensionality reduction via PCA was conducted, followed by batch correction using Harmony. A neighbor graph was constructed based on PC coordinates, which was used for Leiden clustering and UMAP embedding to locate and isolate cartilage cells for further analysis. To achieve better integration, isolated cartilage cells were processed using the scVI suite, which leverages an autoencoder neural network. The latent representation from this process was used to construct a neighbor graph for Leiden clustering and UMAP embedding. Covariates such as heat shock response genes and cell cycle scores were included during model training.

*Analysis Pipelines (references are provided in the main text)*

1. RNA Velocity RNA velocity analysis was performed using the UniT Velo package to define differentiation directions and root cells.
2. SNP Analysis The Monopogen pipeline was employed to infer SNPs from scRNA-seq data. Genes from the first 20 chromosomes were scanned, and QC filters were applied. The resulting table contained SNP positions as rows and individual cells as columns. SNPs were annotated as follows: R for reference allele reads, A for alternative nucleotide (SNP), B for heterozygous variants, 0 if the position was not sequenced.
3. Differentiation Trajectory Analysis ScFates was used to infer a principal graph on 2D cell embeddings, with cells softly assigned to neighboring nodes. The previously determined differentiation trajectory was utilized to set root cells, after which pseudotime scoring was calculated along the graph. Features, genes, or regulons were fitted to the trajectory, enabling the identification of significant changes over the differentiation course.
4. Prediction of Transcription Factor Activity Active transcription factors (TFs) were inferred using Decoupler, which applies a univariate linear model to test whether known downstream genes are upregulated for each TF.
5. SCENIC Analysis Regulons (TFs with their putatively regulated genes) were identified using pySCENIC. Filters were applied to retain only activating and annotated motifs, resulting in 219 active TFs. After overlap with Decoupler results, 147 TFs remained. Cell-wise enrichment scores (AUC) were calculated, and average regulon activity scores were determined for each cluster. Top 20 regulons, sorted by absolute differences in average AUC scores, were visualized in heatmaps.
6. Gene Set Enrichment Analysis (GSEA) Scanpy’s built-in rank_genes_groups method (Wilcoxon statistics) identified highly expressed genes for each cluster. p-values were corrected for multiple testing using the Benjamini-Hochberg approach. Enriched gene sets were identified using GProfiler against the Enrichr Reactome_Pathways_2024 database.
7. Differential Gene Expression (DEG), Volcano Plots, and Heatmaps Differentially expressed genes between gp1 and gp2 were identified using DESeq2, and results were visualized in volcano plots. DEGs (log fold change > 1, adjusted p-value < 0.05) were analyzed with GProfiler for enrichment terms.
8. Ligand-Receptor Pair Analysis The LIANA library was used to identify ligand-receptor (LR) pairs expressed across clusters. Depth-normalized and log-transformed expression matrices were input for analysis. LIANA’s consensus approach integrated multiple prediction methods, ranking LR-pair magnitude and specificity to highlight promising interactions.
9. Prediction of Cell Clusters in Culture Cultured samples were annotated by matching them to populations defined in directly sequenced tissues using the scANVI library, which employs a machine learning-based autoencoder neural network.
10. Bioinformatic Assessment of Cultured and Uncultured Samples Vehicle-treated explants were processed with the same pipeline as uncultured samples to assess the effects of cultivation.
11. Protein-Protein Interaction Analysis Corneto was used to investigate potential pathways linking GHR stimulation to TFs identified by pySCENIC and Decoupler. Protein-protein interactions from the OmniPath database were queried using GHR and the top 20 TFs up- and down-regulated upon GH treatment. The Gurobi solver (academic license) was employed for imputation.
12. Ambient RNA Correction Ambient RNA contamination was corrected using Cell Bender. Genes highly expressed in “empty” 10X beads were identified from cultured samples and excluded from DEG analysis results.

### Statistical Analysis

All values are presented as mean ± standard deviation (SD) unless otherwise stated. Comparisons between two groups (e.g., vehicle-treated and GH-treated slices) were performed using either a paired or unpaired Student’s *t*-test, depending on whether the samples were paired or independent. While experiments were performed in the way that growth plate slices from each patient were cultured with either vehicle or GH (paired samples), during the analysis pipeline and blinding of the samples, analyzed histological sections were not always from the same patients and were regarded as independent samples in this case. For grouped data, one-way ANOVA with Tukey’s correction for multiple comparisons was applied.

All statistical analyses were conducted using GraphPad Prism (version 8.0) unless otherwise specified. A minimum of three independent biological replicates were included in each statistical analysis. Details regarding the specific statistical tests employed for individual experiments are provided in the corresponding figure legends. The following *p*-value thresholds were used to denote statistical significance: *P* < 0.05: represented by *, *P* < 0.005: represented by **, *P* < 0.0005: represented by ***, *P* < 0.00005: represented by ****.

To statistically estimate the proliferation score in scRNA-seq data, a chi-squared contingency test was employed. Each cell was assigned a phase score based on the expression of known cell cycle genes. Cells identified as being in the S or G2/M stages were classified as cycling, rendering the parameter binary. For each cluster, the absolute number of cycling and non-cycling cells was calculated. A pairwise chi-squared contingency test was conducted to compare gp1 against all other clusters. The null hypothesis (*H₀*) posited that the groups compared had the same effect on proliferation, and this hypothesis was tested.

