## Supplementary figures and images for "A transcriptional atlas of the pubertal human growth plate reveals direct stimulation of cartilage stem cells by growth hormone"

### Supplementary file 1

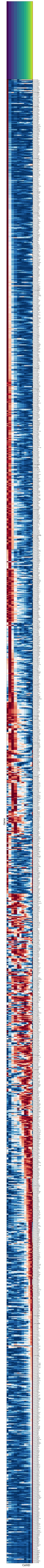

### Supplementary file 2

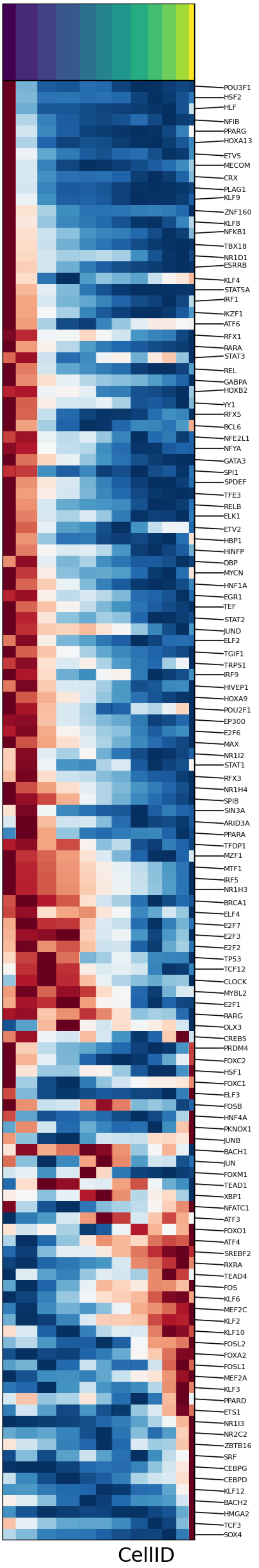

### Supplementary file 4

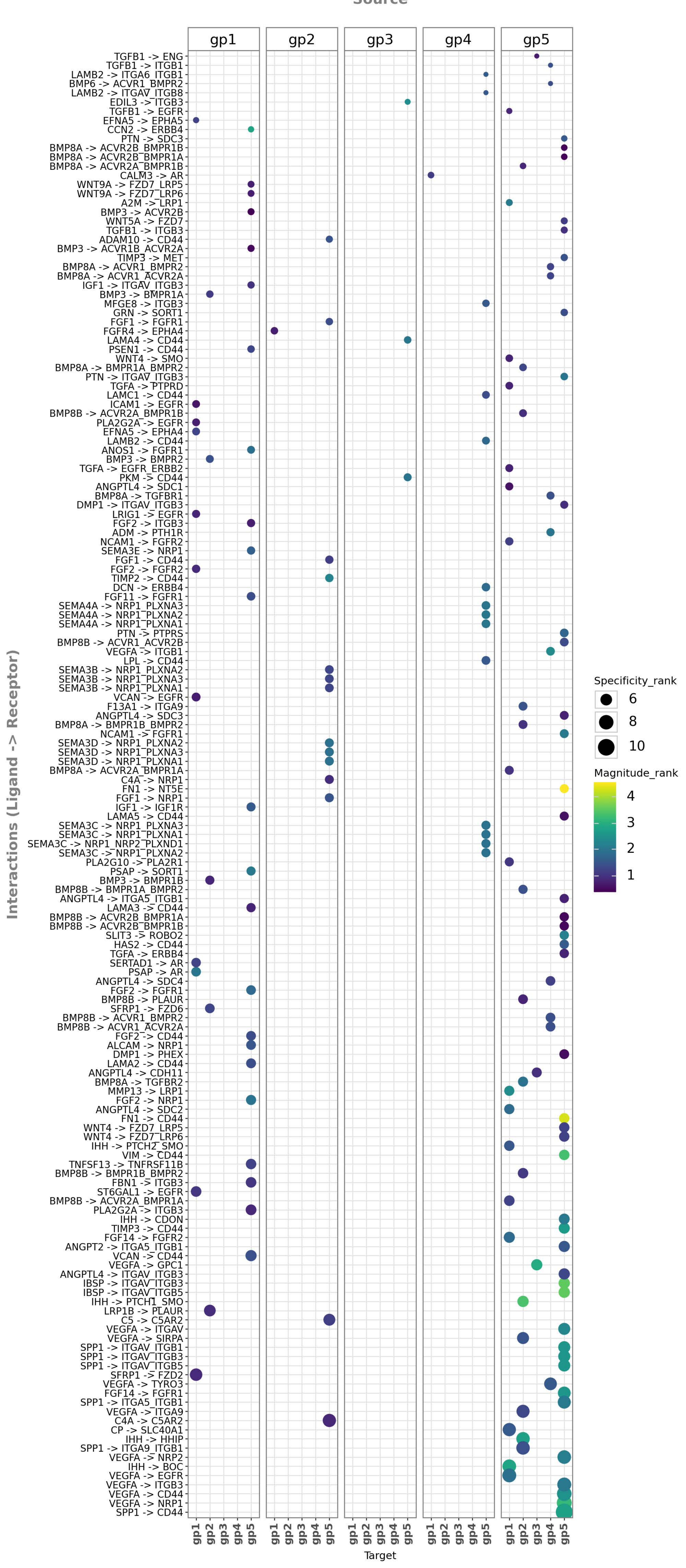

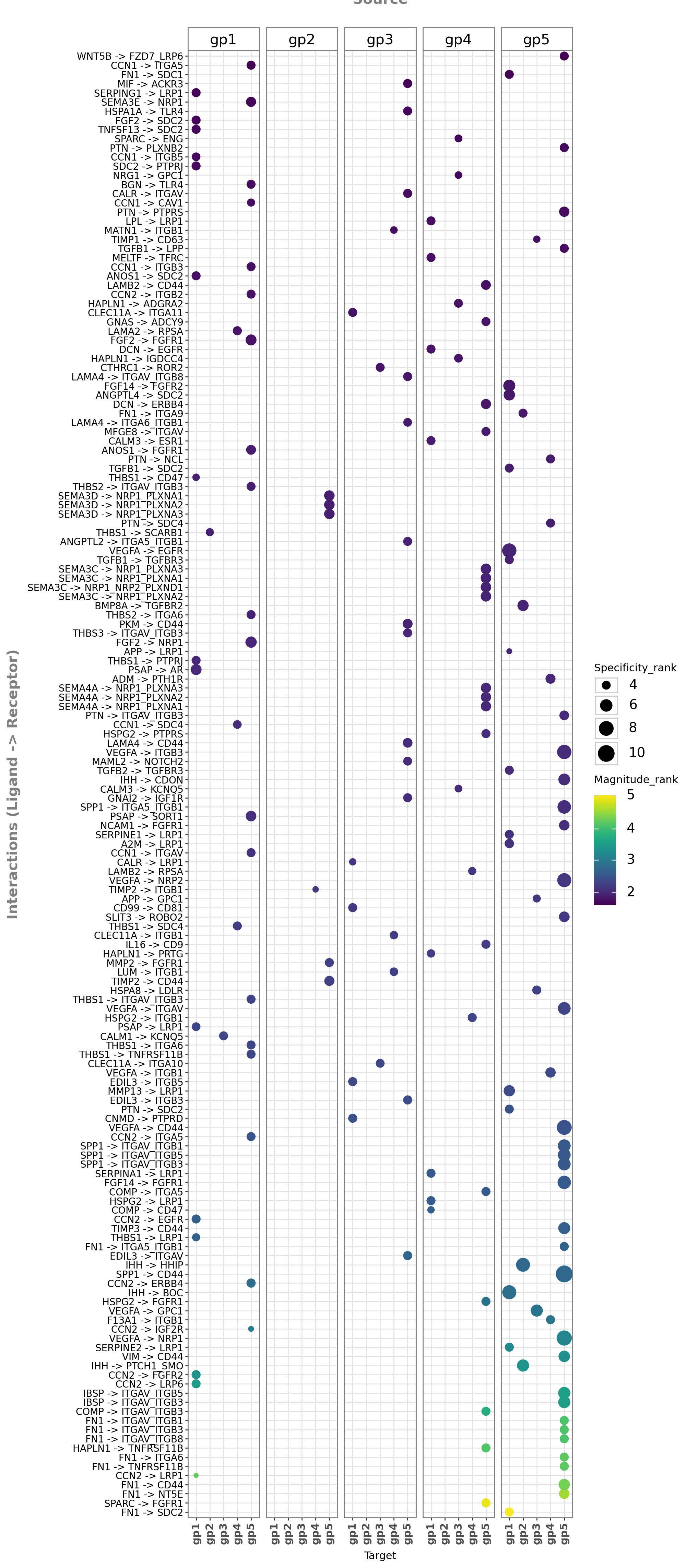
