## Supplementary file 3 for "A transcriptional atlas of the pubertal human growth plate reveals direct stimulation of cartilage stem cells by growth hormone"

In [ ]:

In [197…
```
for reg in adata.var.index:
    scf.pl.single_trend(adata,reg, basis='umap')
```

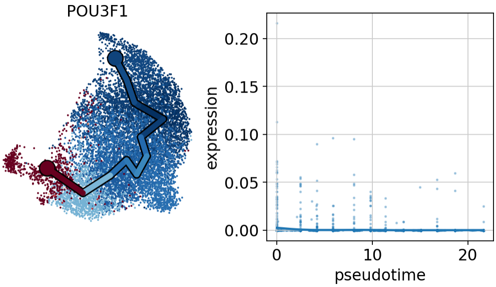


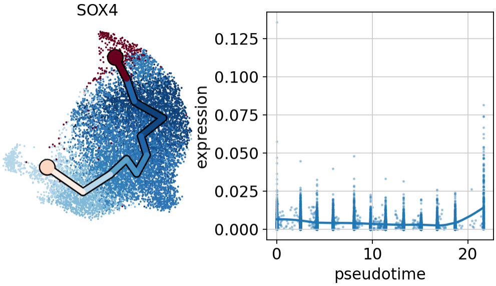


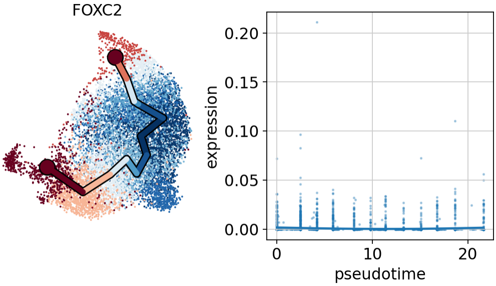


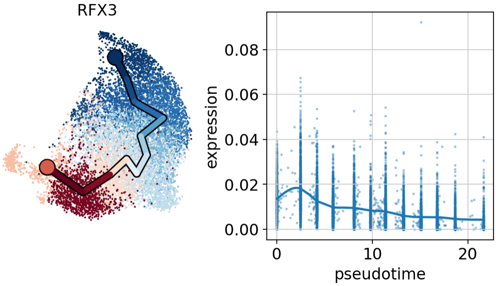

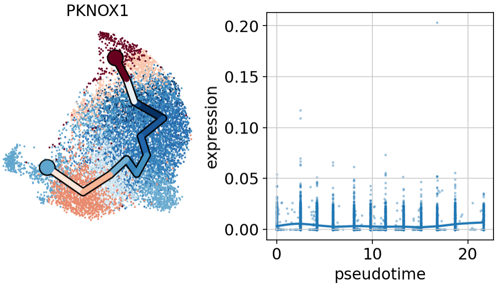
PKNOX1
expression
pseudotime
0.00
0.05
0.10
0.15
0.20
0
10
20

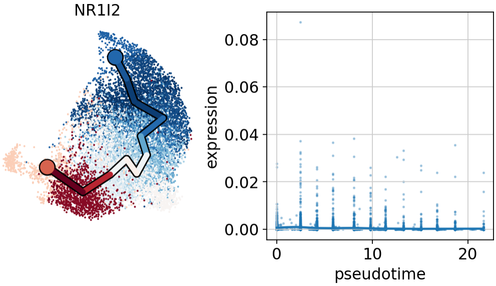
NR1I2
expression
pseudotime
0.00
0.02
0.04
0.06
0.08
0
10
20

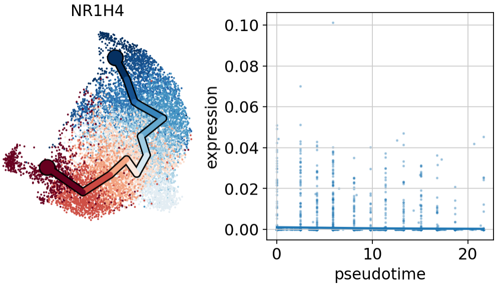
NR1H4
expression
pseudotime
0.00
0.02
0.04
0.06
0.08
0.10
0
10
20

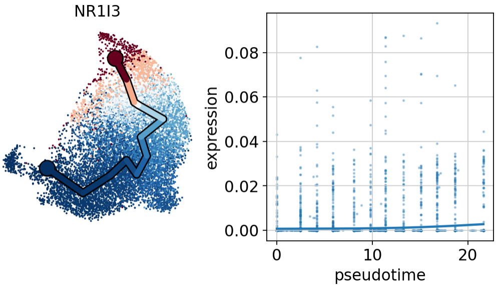
NR1I3
expression
pseudotime
0.00
0.02
0.04
0.06
0.08
0
10
20

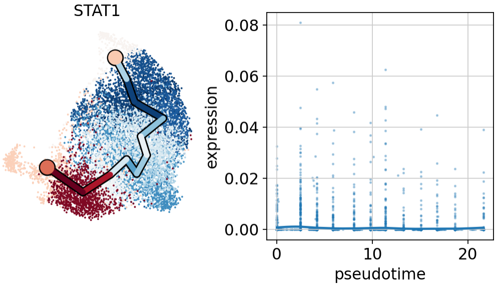
STAT1
expression
pseudotime

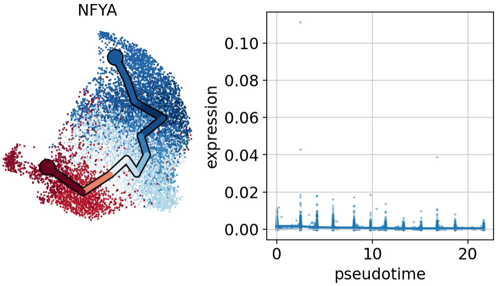
NFYA
expression
pseudotime

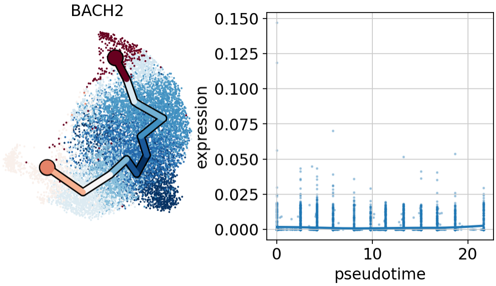
BACH2
expression
pseudotime

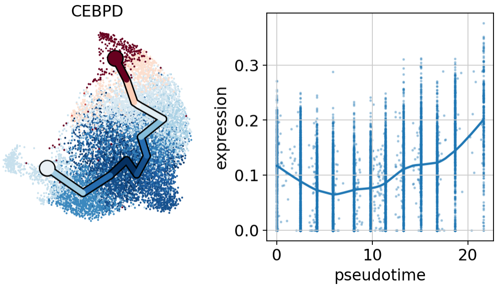
CEBPD
expression
pseudotime

HNF1A

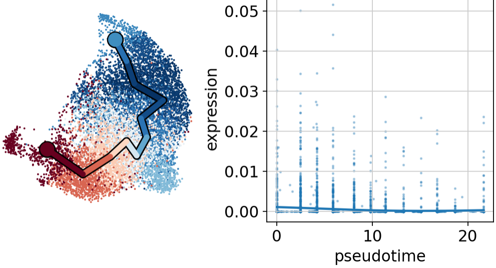


BRCA1

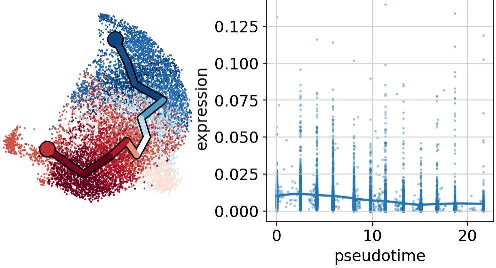


NFATC1

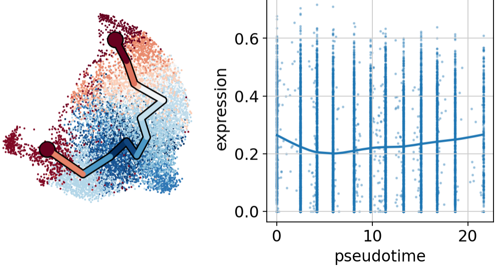


ESRRB

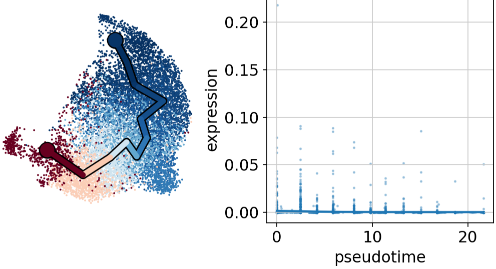

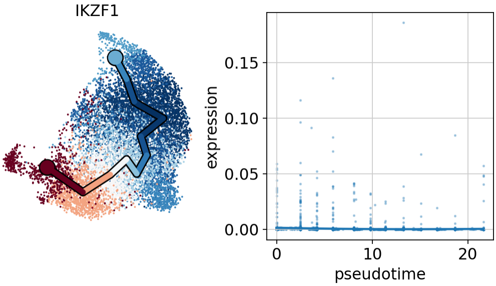
IKZF1
expression
0.15
0.10
0.05
0.00
0
10
20
pseudotime

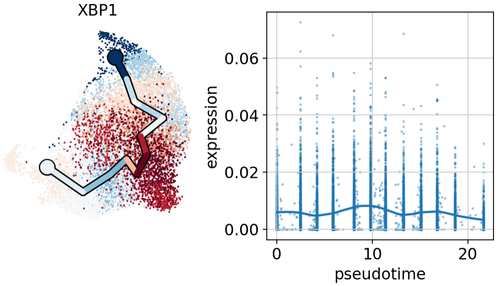
XBP1
expression
0.06
0.04
0.02
0.00
0
10
20
pseudotime

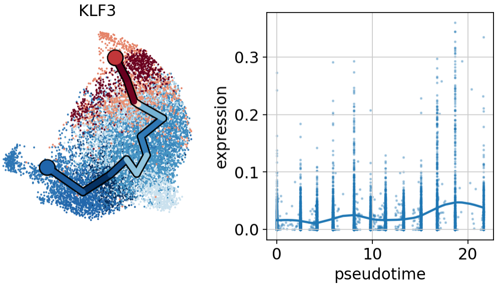
KLF3
expression
0.3
0.2
0.1
0.0
0
10
20
pseudotime

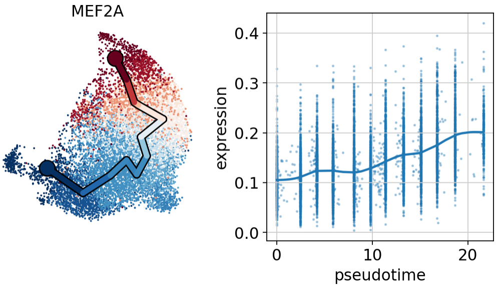
MEF2A
expression
0.4
0.3
0.2
0.1
0.0
0
10
20
pseudotime

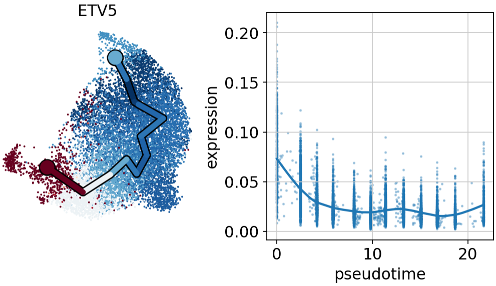
ETV5
0.20
0.15
0.10
0.05
0.00
expression
0
10
20
pseudotime

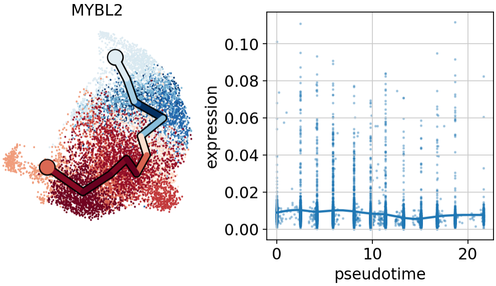
MYBL2
0.10
0.08
0.06
0.04
0.02
0.00
expression
0
10
20
pseudotime

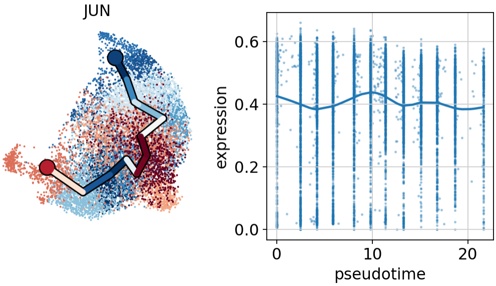
JUN
0.6
0.4
0.2
0.0
expression
0
10
20
pseudotime

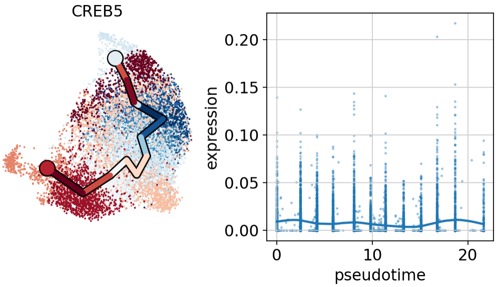
CREB5
0.20
0.15
0.10
0.05
0.00
expression
0
10
20
pseudotime

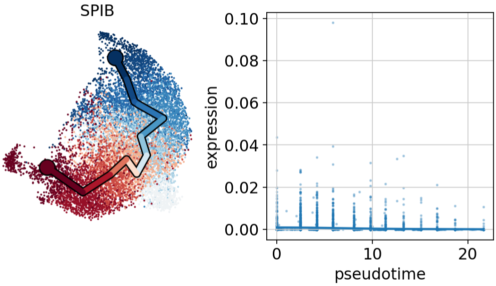
SPIB
0.10
0.08
0.06
0.04
0.02
0.00
expression
0
10
20
pseudotime

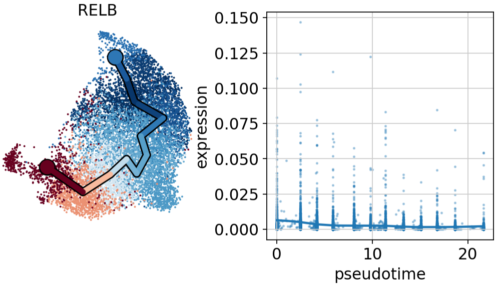
RELB
0.150
0.125
0.100
0.075
0.050
0.025
0.000
expression
0
10
20
pseudotime

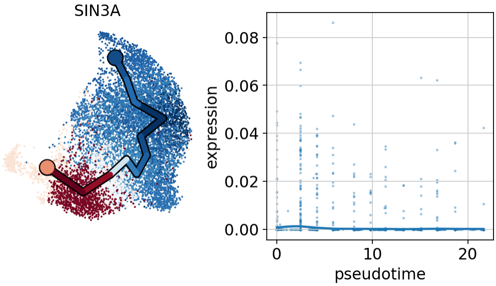
SIN3A
0.08
0.06
0.04
0.02
0.00
expression
0
10
20
pseudotime

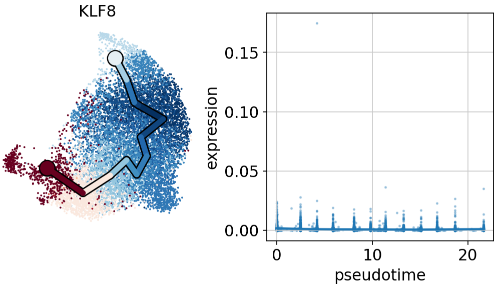
KLF8
0.15
0.10
0.05
0.00
expression
0
10
20
pseudotime

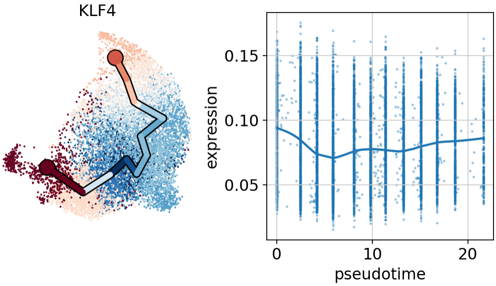
KLF4
0.15
0.10
0.05
expression
0
10
20
pseudotime

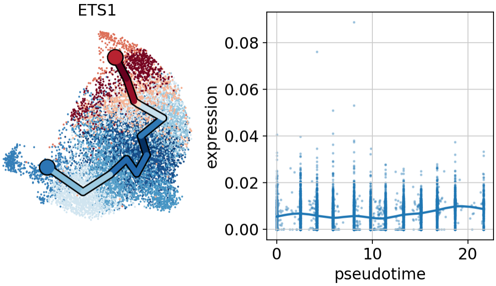
ETS1
0.08
0.06
0.04
0.02
0.00
expression
0
10
20
pseudotime

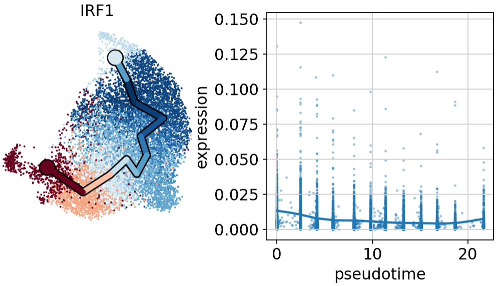
IRF1
0.150
0.125
0.100
0.075
0.050
0.025
0.000
expression
0
10
20
pseudotime

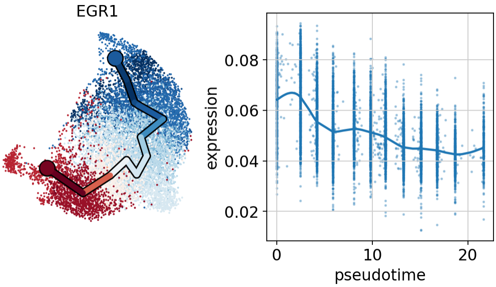
EGR1
0.08
0.06
0.04
0.02
expression
0
10
20
pseudotime

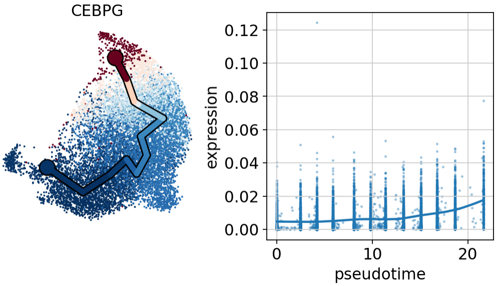
CEBPG
0.12
0.10
0.08
0.06
0.04
0.02
0.00
expression
0
10
20
pseudotime

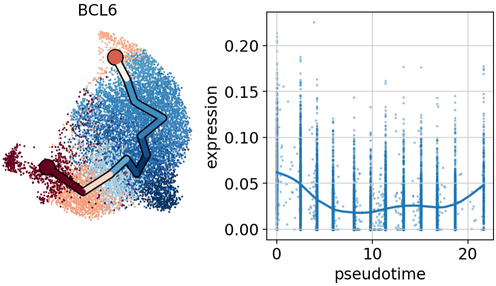
BCL6
0.20
0.15
0.10
0.05
0.00
expression
0
10
20
pseudotime

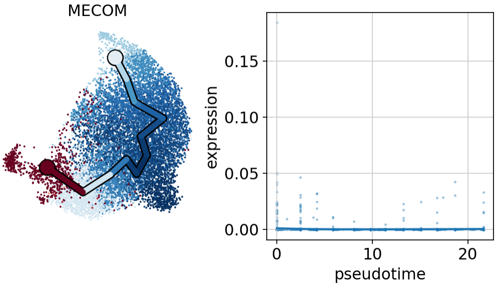
MECOM
0.15
0.10
0.05
0.00
expression
0
10
20
pseudotime

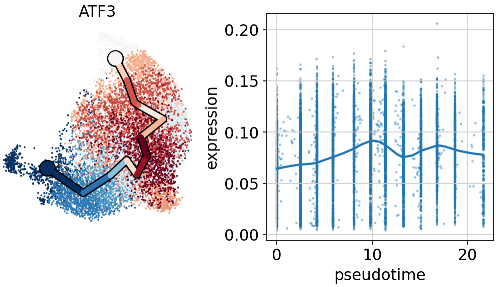
ATF3
0.20
0.15
0.10
0.05
0.00
expression
0
10
20
pseudotime

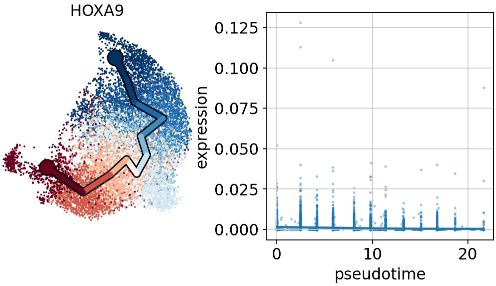
HOXA9
0.125
0.100
0.075
0.050
0.025
0.000
expression
0
10
20
pseudotime

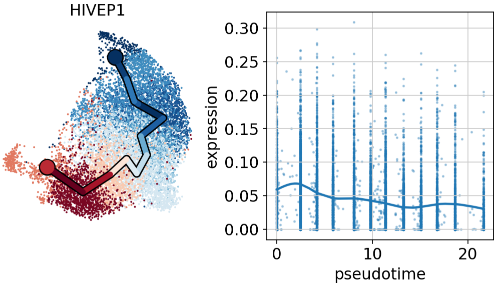
HIVEP1
0.30
0.25
0.20
0.15
0.10
0.05
0.00
expression
0
10
20
pseudotime

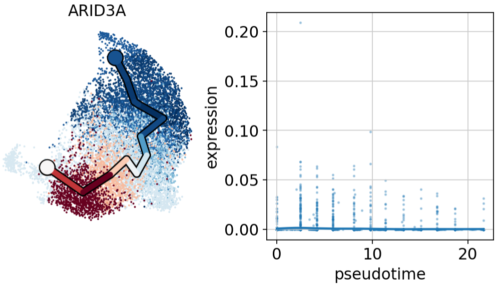
ARID3A
0.20
0.15
0.10
0.05
0.00
expression
0
10
20
pseudotime

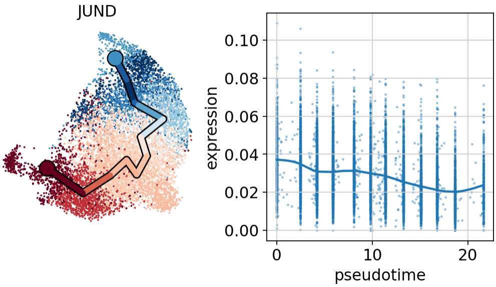
JUND
0.10
0.08
0.06
0.04
0.02
0.00
expression
0
10
20
pseudotime

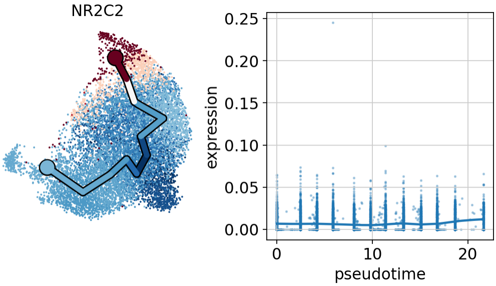
NR2C2
0.25
0.20
0.15
0.10
0.05
0.00
expression
0
10
20
pseudotime

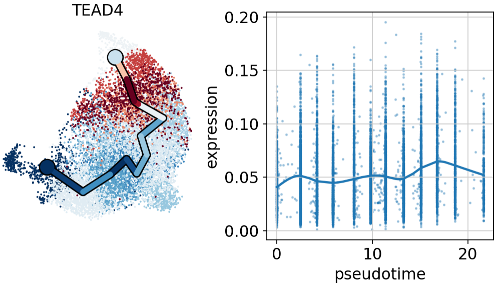
TEAD4
0.20
0.15
0.10
0.05
0.00
expression
0
10
20
pseudotime

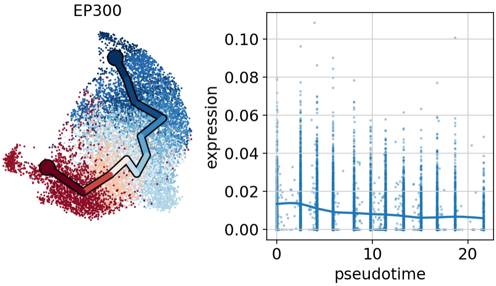
EP300
0.10
0.08
0.06
0.04
0.02
0.00
expression
0
10
20
pseudotime

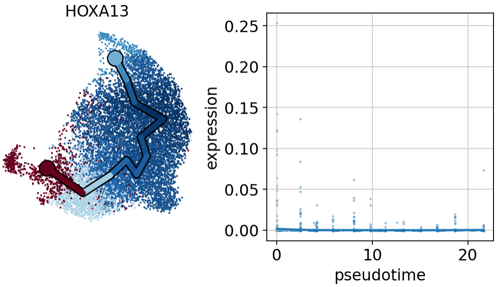
HOXA13
0.25
0.20
0.15
0.10
0.05
0.00
expression
0
10
20
pseudotime

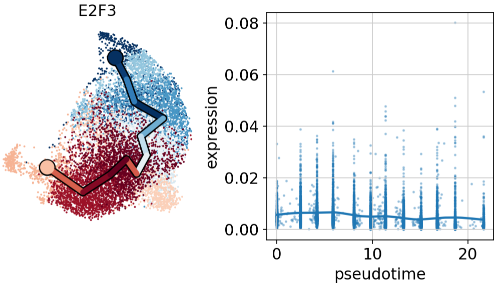
E2F3
expression
0.08
0.06
0.04
0.02
0.00
0
10
20
pseudotime

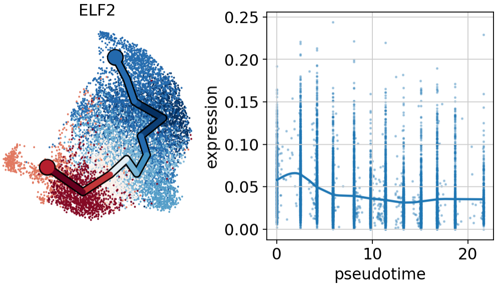
ELF2
expression
0.25
0.20
0.15
0.10
0.05
0.00
0
10
20
pseudotime

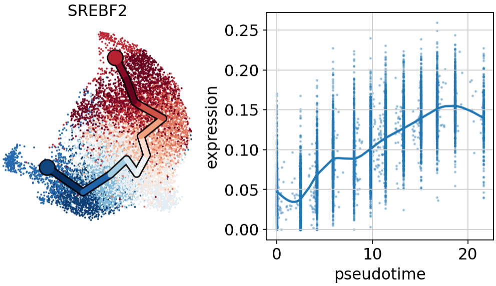
SREBF2
expression
0.25
0.20
0.15
0.10
0.05
0.00
0
10
20
pseudotime

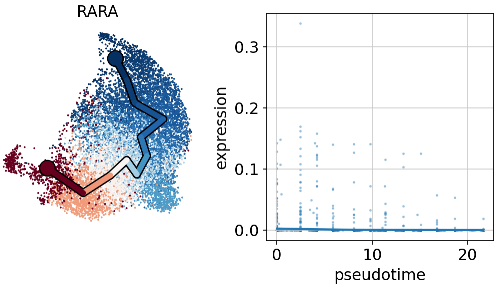
RARA
expression
0.3
0.2
0.1
0.0
0
10
20
pseudotime

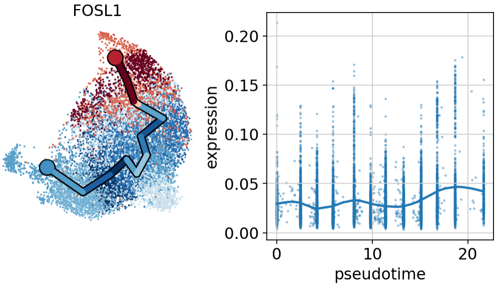
FOSL1
0.20
0.15
0.10
0.05
0.00
expression
0
10
20
pseudotime

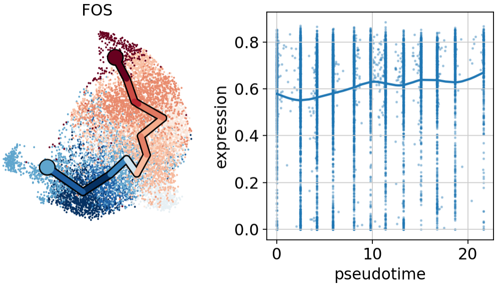
FOS
0.8
0.6
0.4
0.2
0.0
expression
0
10
20
pseudotime

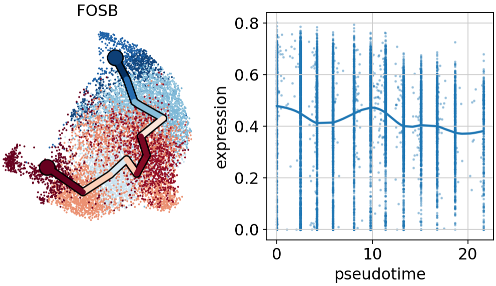
FOSB
0.8
0.6
0.4
0.2
0.0
expression
0
10
20
pseudotime

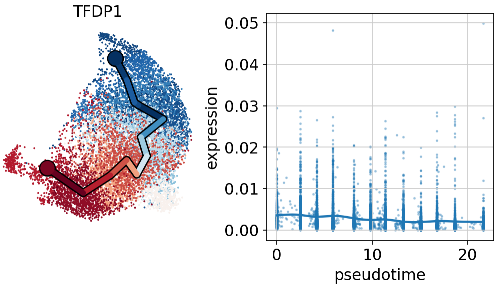
TFDP1
0.05
0.04
0.03
0.02
0.01
0.00
expression
0
10
20
pseudotime

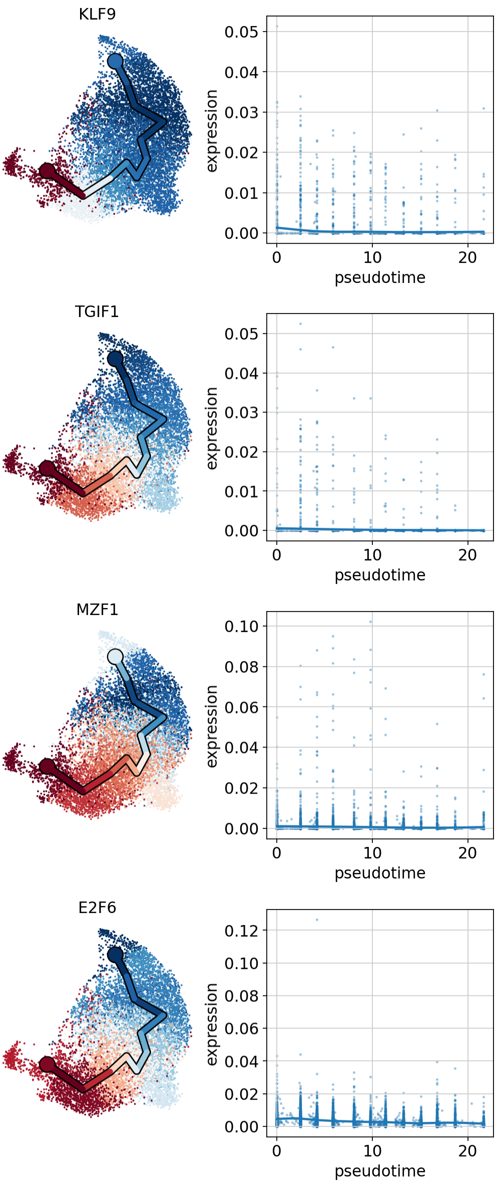
KLF9
expression
pseudotime
TGIF1
expression
pseudotime
MZF1
expression
pseudotime
E2F6
expression
pseudotime

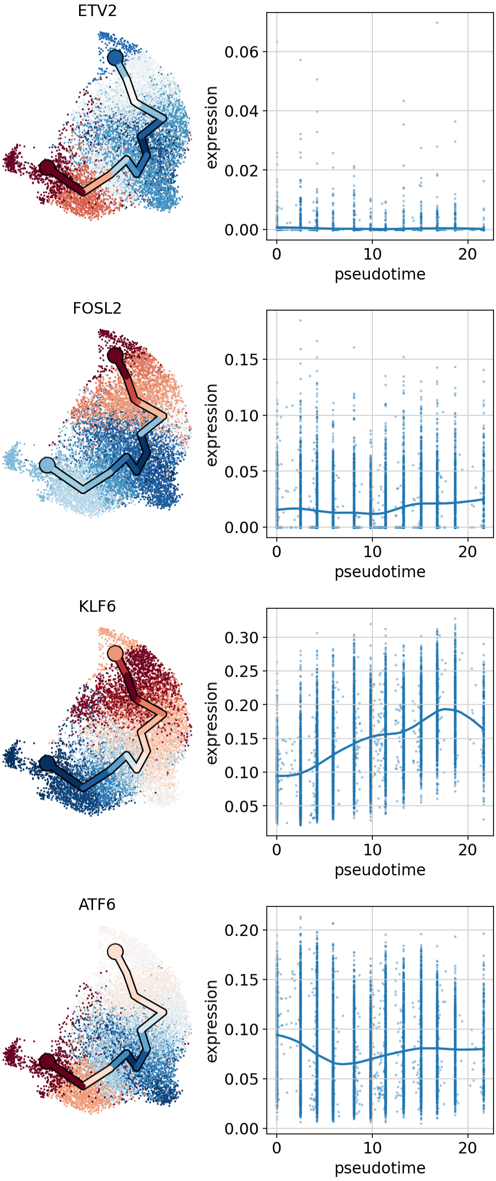
ETV2
0.06
0.04
0.02
0.00
expression
0
10
20
pseudotime
FOSL2
0.15
0.10
0.05
0.00
expression
0
10
20
pseudotime
KLF6
0.30
0.25
0.20
0.15
0.10
0.05
expression
0
10
20
pseudotime
ATF6
0.20
0.15
0.10
0.05
0.00
expression
0
10
20
pseudotime

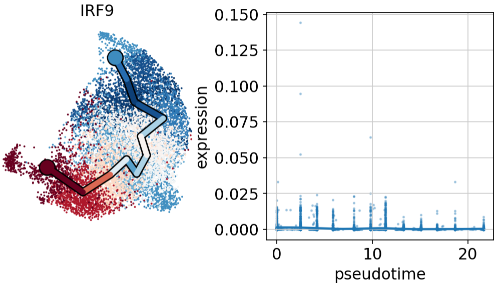
IRF9
0.150
0.125
0.100
0.075
0.050
0.025
0.000
expression
0
10
20
pseudotime

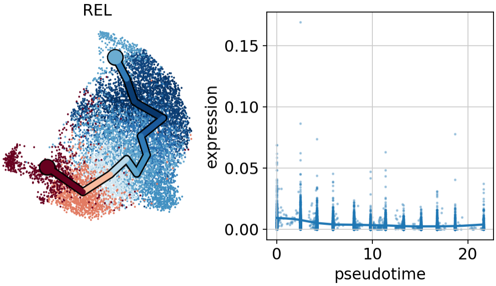
REL
0.15
0.10
0.05
0.00
expression
0
10
20
pseudotime

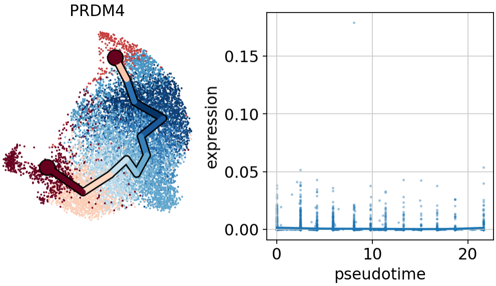
PRDM4
0.15
0.10
0.05
0.00
expression
0
10
20
pseudotime

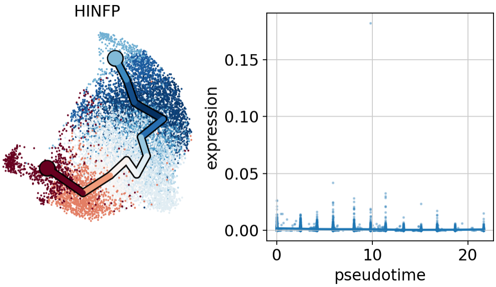
HINFP
0.15
0.10
0.05
0.00
expression
0
10
20
pseudotime

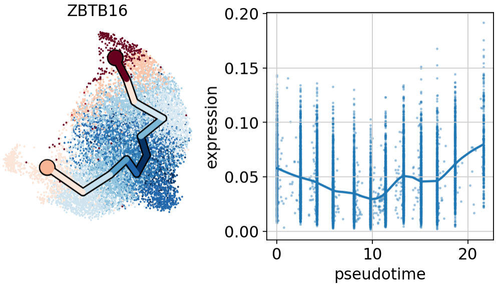
ZBTB16
0.20
0.15
0.10
0.05
0.00
expression
0
10
20
pseudotime

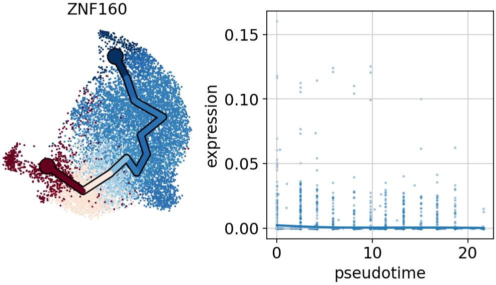
ZNF160
0.15
0.10
0.05
0.00
expression
0
10
20
pseudotime

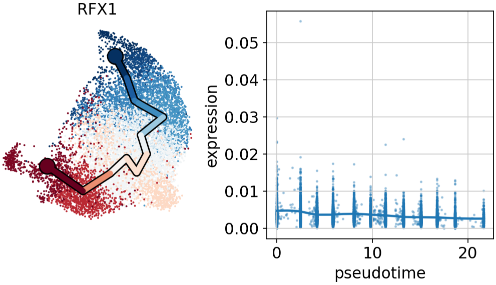
RFX1
0.05
0.04
0.03
0.02
0.01
0.00
expression
0
10
20
pseudotime

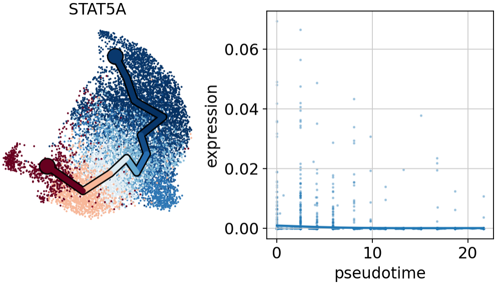
STAT5A
0.06
0.04
0.02
0.00
expression
0
10
20
pseudotime

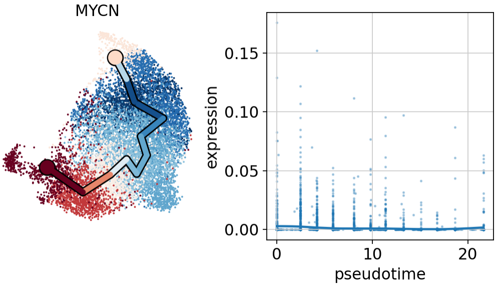
MYCN
0.15
0.10
0.05
0.00
expression
0
10
20
pseudotime

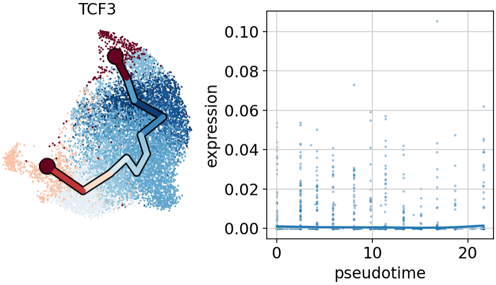
TCF3
0.10
0.08
0.06
0.04
0.02
0.00
expression
0
10
20
pseudotime

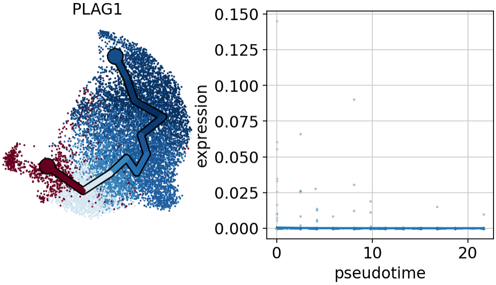
PLAG1
0.150
0.125
0.100
0.075
0.050
0.025
0.000
expression
0
10
20
pseudotime

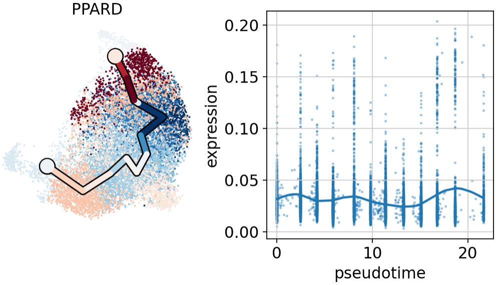
PPARD
0.20
0.15
0.10
0.05
0.00
expression
0
10
20
pseudotime

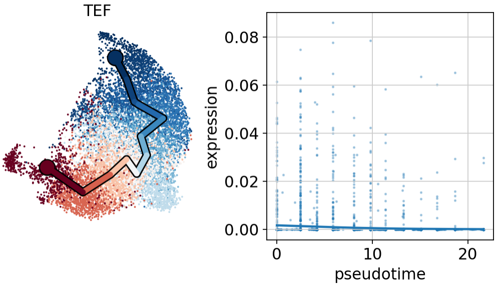
TEF
0.08
0.06
0.04
0.02
0.00
expression
0
10
20
pseudotime

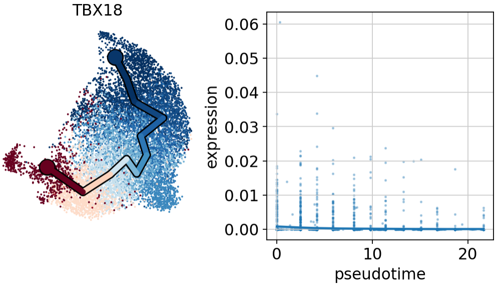
TBX18
0.06
0.05
0.04
0.03
0.02
0.01
0.00
expression
0
10
20
pseudotime

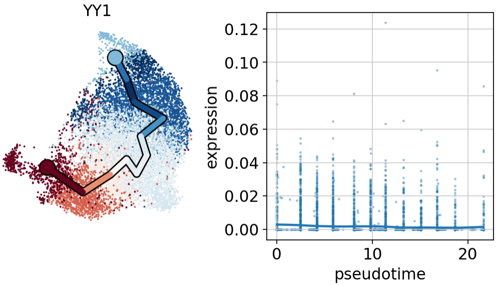
YY1
0.12
0.10
0.08
0.06
0.04
0.02
0.00
expression
0
10
20
pseudotime

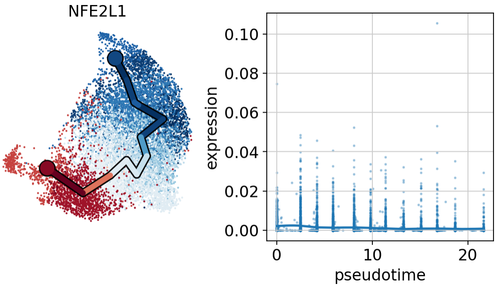
NFE2L1
0.10
0.08
0.06
0.04
0.02
0.00
expression
0
10
20
pseudotime

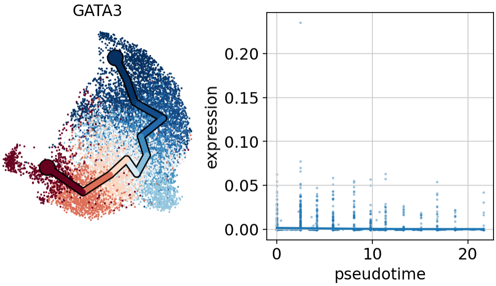
GATA3
0.20
0.15
0.10
0.05
0.00
expression
0
10
20
pseudotime

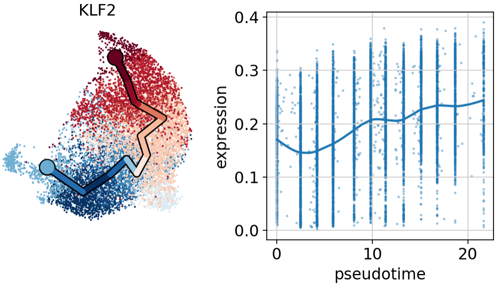
KLF2
0.4
0.3
0.2
0.1
0.0
expression
0
10
20
pseudotime

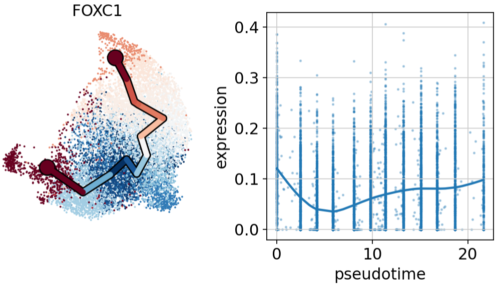
FOXC1
0.4
0.3
0.2
0.1
0.0
expression
0
10
20
pseudotime

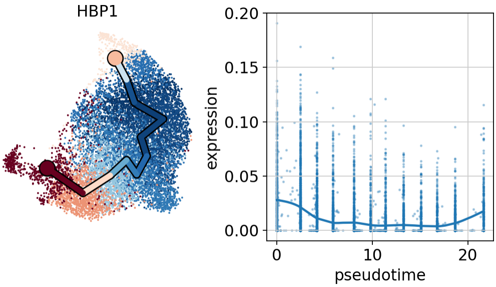
HBP1
0.20
0.15
0.10
0.05
0.00
expression
0
10
20
pseudotime

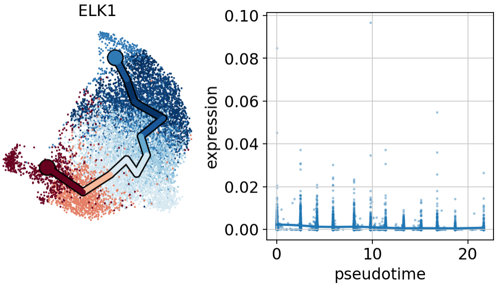
ELK1
0.10
0.08
0.06
0.04
0.02
0.00
expression
0
10
20
pseudotime

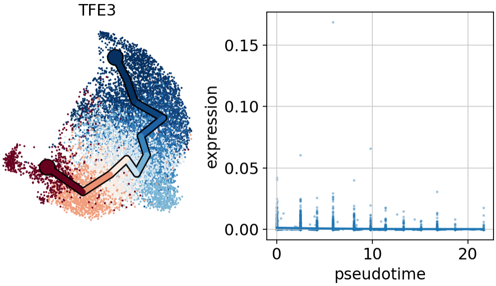
TFE3
0.15
0.10
0.05
0.00
expression
0
10
20
pseudotime

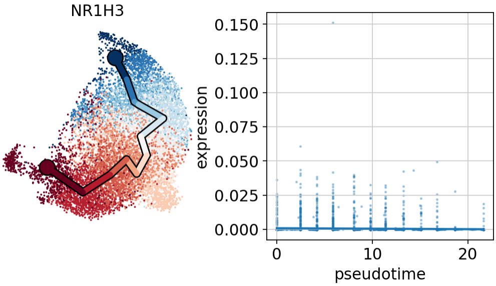
NR1H3
0.150
0.125
0.100
0.075
0.050
0.025
0.000
expression
0
10
20
pseudotime

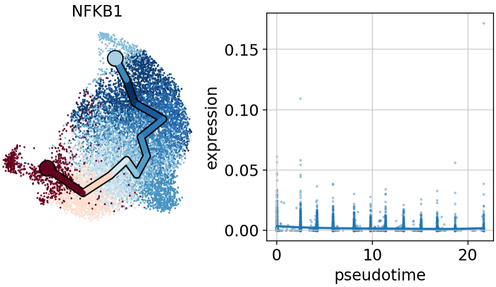
NFKB1
0.15
0.10
0.05
0.00
expression
0
10
20
pseudotime

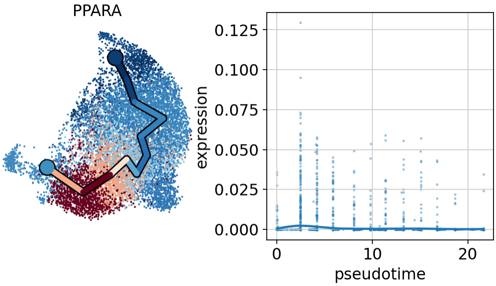
PPARA
0.125
0.100
0.075
0.050
0.025
0.000
expression
0
10
20
pseudotime

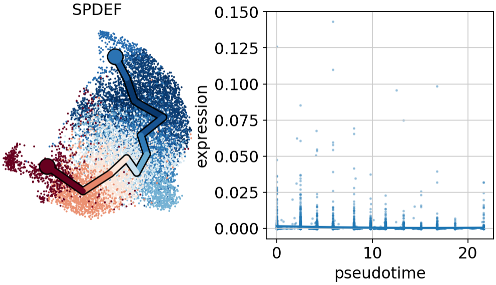
SPDEF
0.150
0.125
0.100
0.075
0.050
0.025
0.000
expression
0
10
20
pseudotime

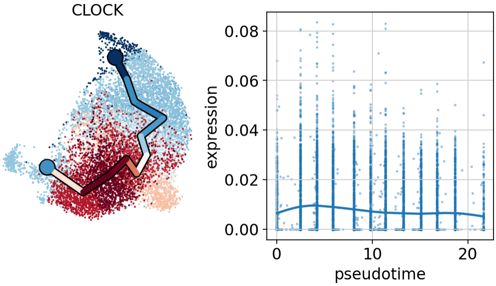
CLOCK
0.08
0.06
0.04
0.02
0.00
expression
0
10
20
pseudotime

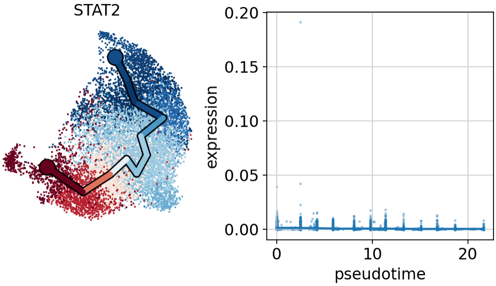
STAT2
0.20
0.15
0.10
0.05
0.00
expression
0
10
20
pseudotime

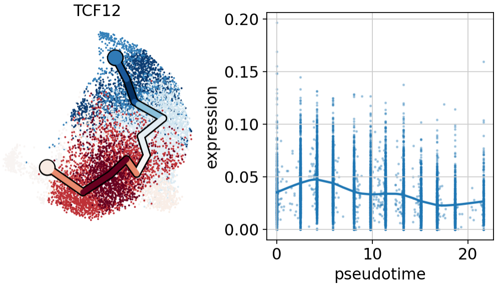
TCF12
0.20
0.15
0.10
0.05
0.00
expression
0
10
20
pseudotime

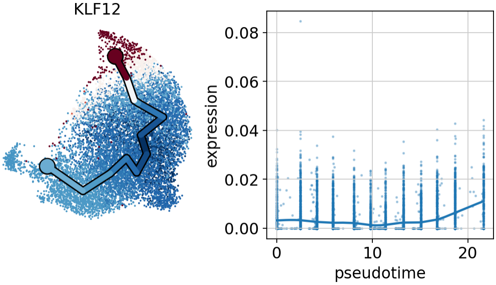
KLF12
0.08
0.06
0.04
0.02
0.00
expression
0
10
20
pseudotime

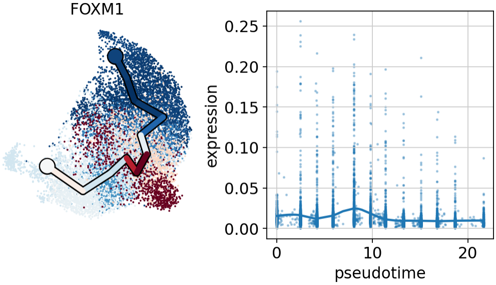
FOXM1
0.25
0.20
0.15
0.10
0.05
0.00
expression
0
10
20
pseudotime

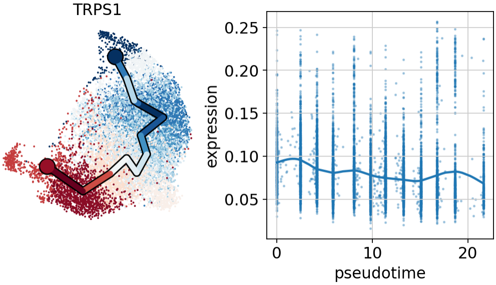
TRPS1
0.25
0.20
0.15
0.10
0.05
expression
0
10
20
pseudotime

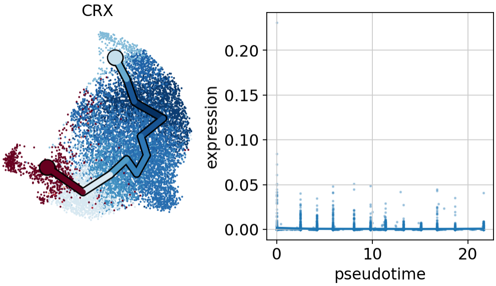
CRX
0.20
0.15
0.10
0.05
0.00
expression
0
10
20
pseudotime

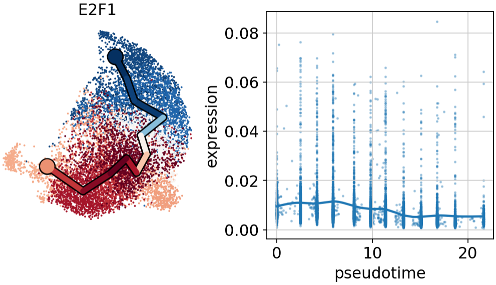
E2F1
0.08
0.06
0.04
0.02
0.00
expression
0
10
20
pseudotime

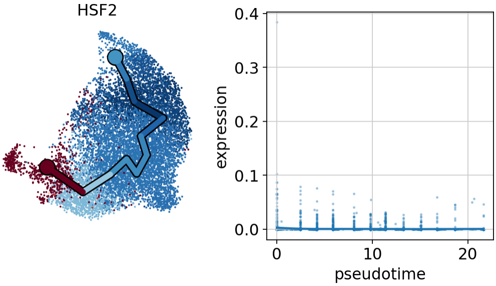
HSF2
0.4
0.3
0.2
0.1
0.0
expression
0
10
20
pseudotime

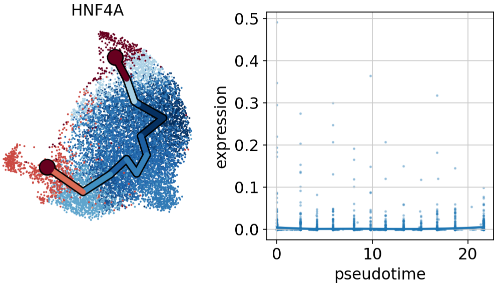
HNF4A
0.5
0.4
0.3
0.2
0.1
0.0
expression
0
10
20
pseudotime

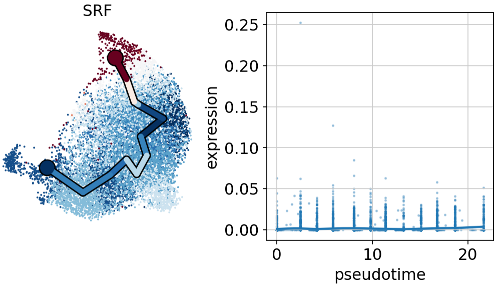
SRF
expression
0.25
0.20
0.15
0.10
0.05
0.00
0
10
20
pseudotime

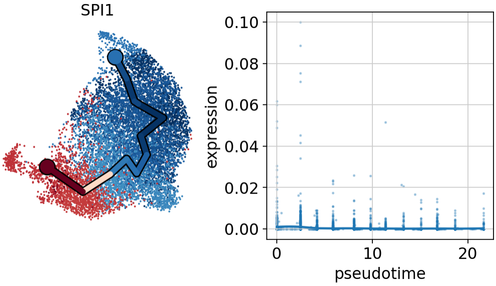
SPI1
expression
0.10
0.08
0.06
0.04
0.02
0.00
0
10
20
pseudotime

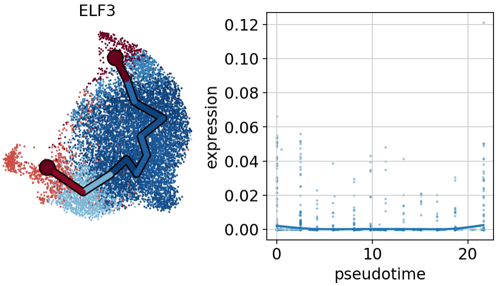
ELF3
expression
0.12
0.10
0.08
0.06
0.04
0.02
0.00
0
10
20
pseudotime

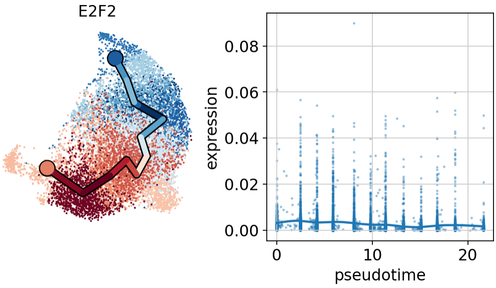
E2F2
expression
0.08
0.06
0.04
0.02
0.00
0
10
20
pseudotime

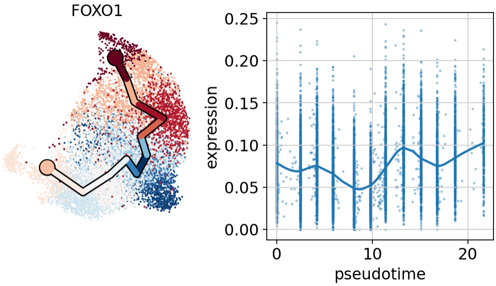
FOXO1
0.25
0.20
0.15
0.10
0.05
0.00
expression
0
10
20
pseudotime

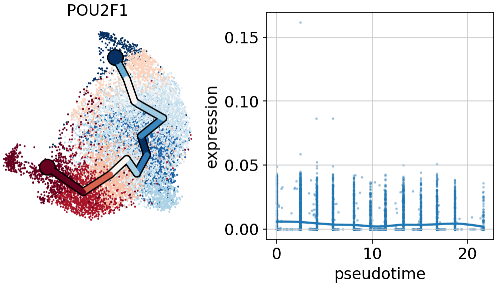
POU2F1
0.15
0.10
0.05
0.00
expression
0
10
20
pseudotime

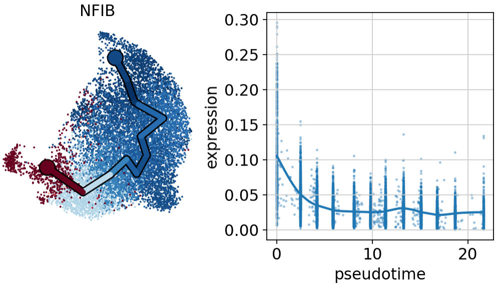
NFIB
0.30
0.25
0.20
0.15
0.10
0.05
0.00
expression
0
10
20
pseudotime

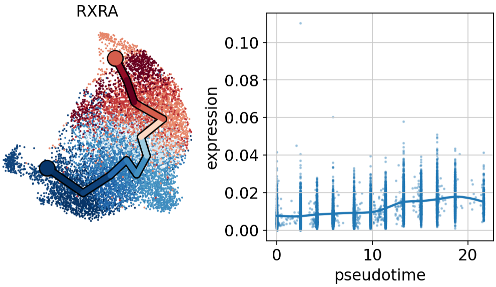
RXRA
0.10
0.08
0.06
0.04
0.02
0.00
expression
0
10
20
pseudotime

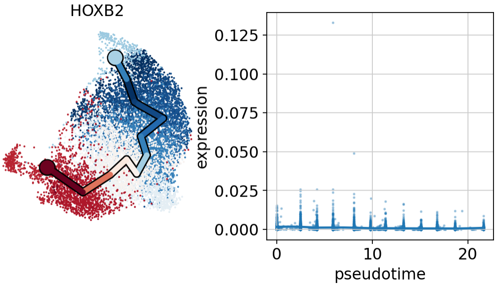
HOXB2
expression
0.125
0.100
0.075
0.050
0.025
0.000
0
10
20
pseudotime

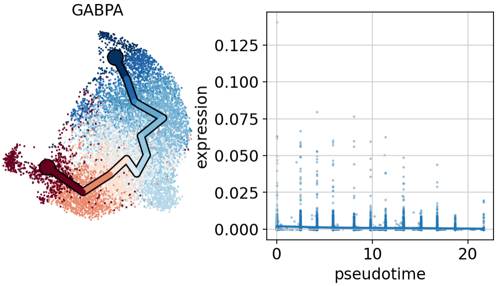
GABPA
expression
0.125
0.100
0.075
0.050
0.025
0.000
0
10
20
pseudotime

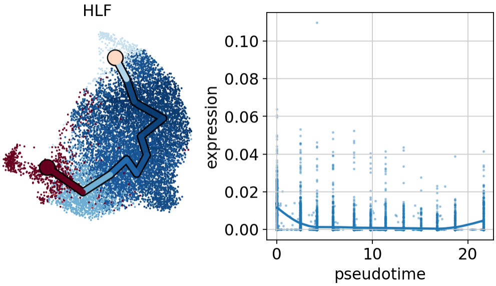
HLF
expression
0.10
0.08
0.06
0.04
0.02
0.00
0
10
20
pseudotime

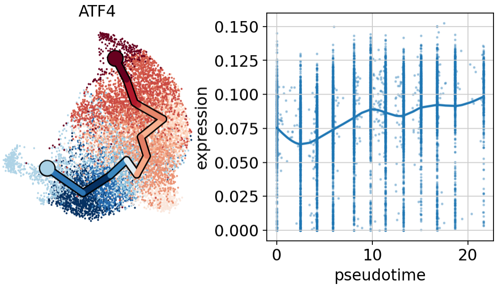
ATF4
expression
0.150
0.125
0.100
0.075
0.050
0.025
0.000
0
10
20
pseudotime

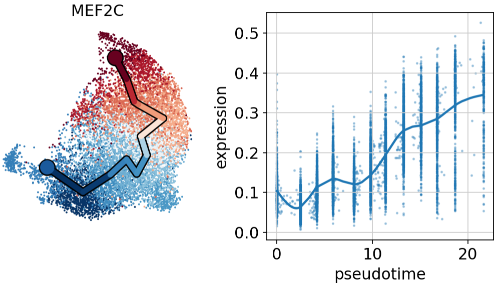
MEF2C
0.5
0.4
0.3
0.2
0.1
0.0
expression
0
10
20
pseudotime

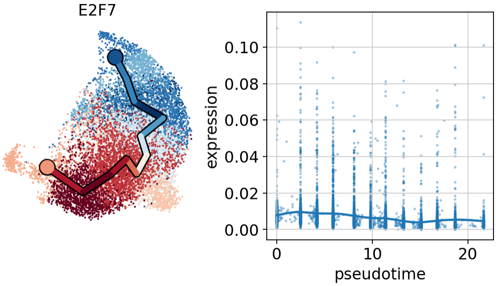
E2F7
0.10
0.08
0.06
0.04
0.02
0.00
expression
0
10
20
pseudotime

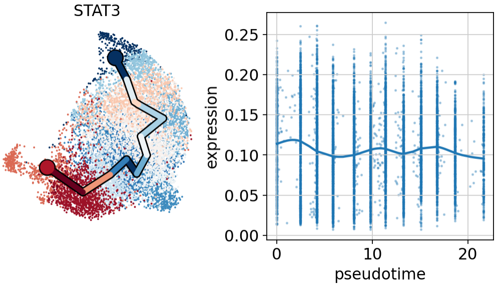
STAT3
0.25
0.20
0.15
0.10
0.05
0.00
expression
0
10
20
pseudotime

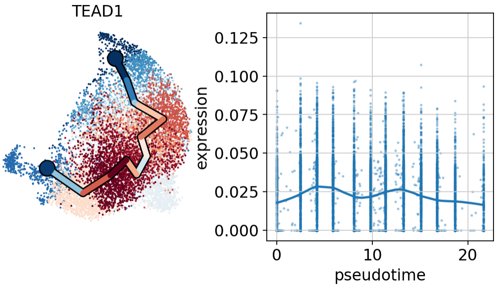
TEAD1
0.125
0.100
0.075
0.050
0.025
0.000
expression
0
10
20
pseudotime

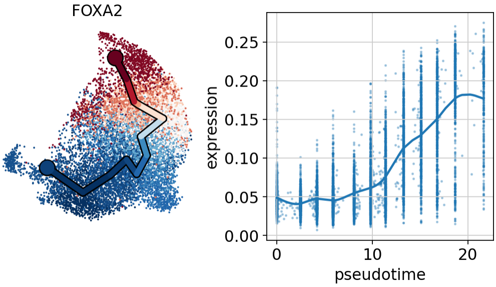
FOXA2
0.25
0.20
0.15
0.10
0.05
0.00
expression
0
10
20
pseudotime

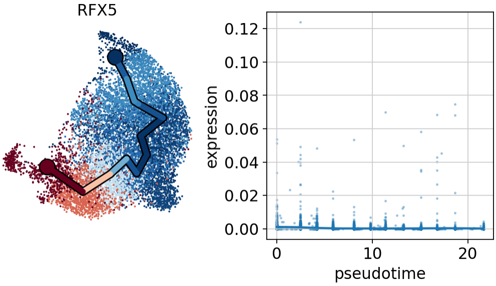
RFX5
0.12
0.10
0.08
0.06
0.04
0.02
0.00
expression
0
10
20
pseudotime

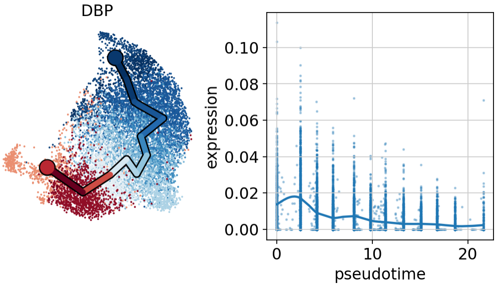
DBP
0.10
0.08
0.06
0.04
0.02
0.00
expression
0
10
20
pseudotime

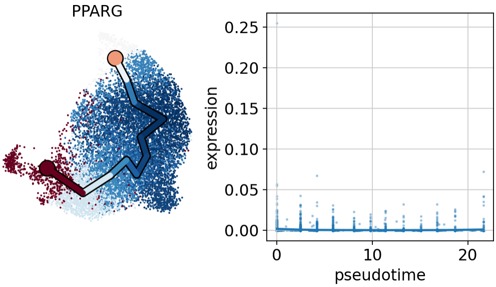
PPARG
0.25
0.20
0.15
0.10
0.05
0.00
expression
0
10
20
pseudotime

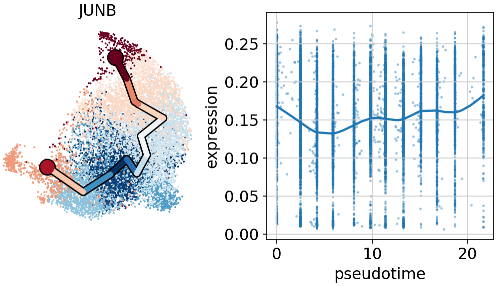
JUNB
expression
0.00
0.05
0.10
0.15
0.20
0.25
0
10
20
pseudotime

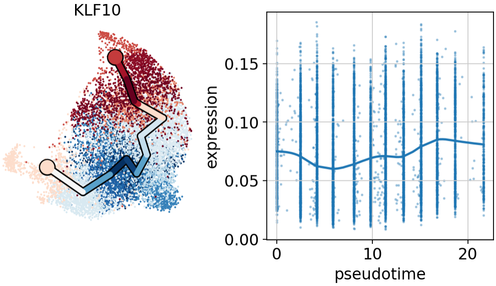
KLF10
expression
0.00
0.05
0.10
0.15
0
10
20
pseudotime

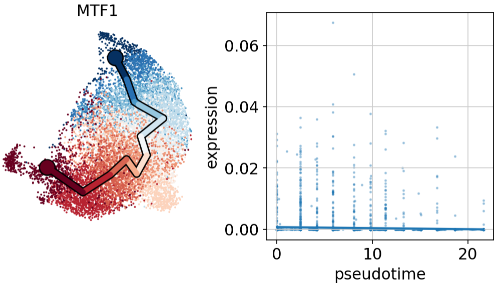
MTF1
expression
0.00
0.02
0.04
0.06
0
10
20
pseudotime

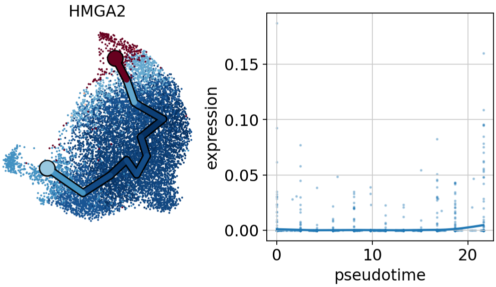
HMGA2
expression
0.00
0.05
0.10
0.15
0
10
20
pseudotime

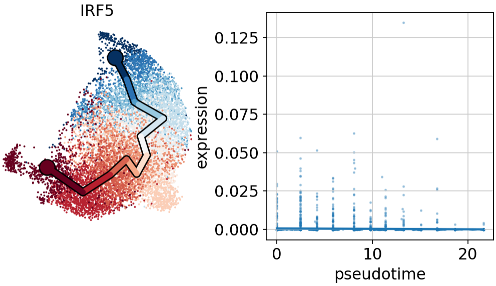
IRF5
0.125
0.100
0.075
0.050
0.025
0.000
expression
0
10
20
pseudotime

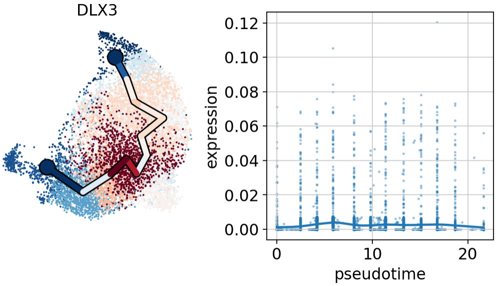
DLX3
0.12
0.10
0.08
0.06
0.04
0.02
0.00
expression
0
10
20
pseudotime

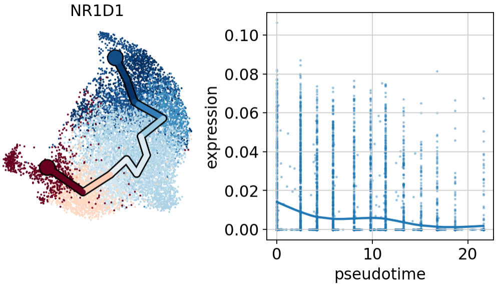
NR1D1
0.10
0.08
0.06
0.04
0.02
0.00
expression
0
10
20
pseudotime

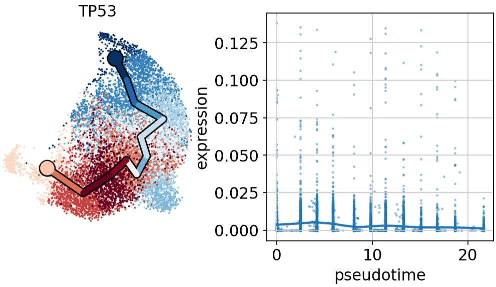
TP53
0.125
0.100
0.075
0.050
0.025
0.000
expression
0
10
20
pseudotime

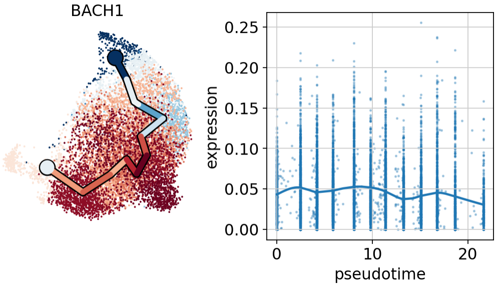
BACH1
0.25
0.20
0.15
0.10
0.05
0.00
expression
0
10
20
pseudotime

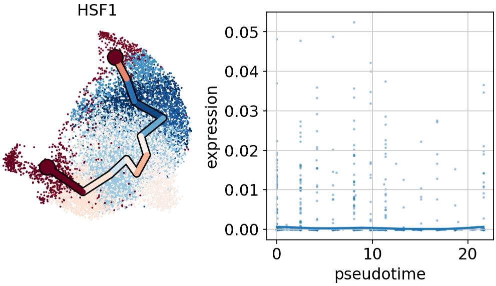
HSF1
0.05
0.04
0.03
0.02
0.01
0.00
expression
0
10
20
pseudotime

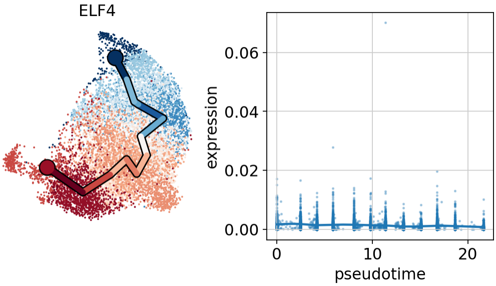
ELF4
0.06
0.04
0.02
0.00
expression
0
10
20
pseudotime

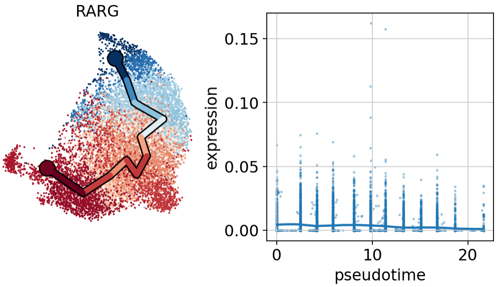
RARG
0.15
0.10
0.05
0.00
expression
0
10
20
pseudotime

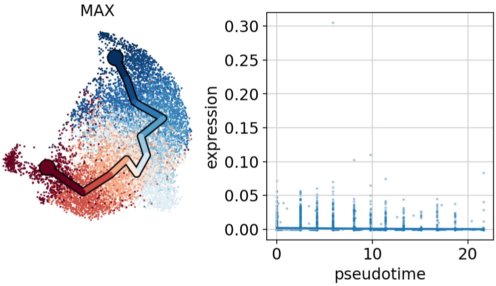
MAX
0.30
0.25
0.20
0.15
0.10
0.05
0.00
expression
0
10
20
pseudotime
